# Cryo-EM elucidation of stoichiometric plasticity, asymmetric ligand recognition and allosteric coupling in human P2X2/3 heterotrimeric channels

**DOI:** 10.64898/2026.04.04.716525

**Authors:** Dong-Ping Wang, Wanbiao Chen, Xiao-Na Yang, Meng-Yang Sun, Ai-Xin Zhang, Yuan Gao, Xi Chen, Bin Cui, Xing Zhou, Yu Gao, Bei-Bei Ding, Yun Tian, Michael X. Zhu, Chang-Zhu Li, Chang-Run Guo, Chongyuan Wang, Ye Yu

## Abstract

P2X receptors are trimeric ATP-gated ion channels that assemble as homo- or heterotrimers, with heteromeric forms exhibiting intrinsic asymmetry that influences function. Here, we report four high-resolution cryo–EM structures of human P2X2/3 heterotrimers representing distinct functional states, including ATP-bound assemblies (P2X332 and P2X223), the apo form, and a ligand/ATP-bound closed conformation. The three ATP-binding sites show asymmetric recognition of MgATP²⁻ and ATP⁴⁻, and channel activation requires occupancy of only two MgATP²⁻ molecules. Gefapixant binds a single allosteric site and selectively inhibits MgATP²⁻, but not ATP⁴⁻, binding, indicating orthosteric–allosteric coupling within the heterotrimer. Structural features of the transmembrane domain define ion permeation, particularly for Ca²⁺. Despite asymmetric ligand interactions, gating remains largely symmetric, with minor differences in desensitization. These findings provide a structural framework linking asymmetry to coordinated channel function and open avenues for subtype-selective therapeutic intervention.

## Introduction

Extracellular ATP acts as an important signaling molecule and neurotransmitter via purinergic P2X receptors, mediating intercellular communication and sensory transmission^1^. Seven P2X receptor subtypes (P2X1–P2X7) have been identified^1^. Like other ligand-gated ion channels, P2X receptors assemble as oligomeric complexes, with trimers representing the minimal functional unit. The functional properties of P2X receptors, including P2X3, are not restricted to homomeric assemblies^2^. Heterologous expression studies indicate that P2X3 and P2X2 subunits can co-assemble into a P2X2/3 heterotrimer composed of two P2X3 and one P2X2 subunits^3^. However, it remains unclear whether this reflects their assembly in native sensory neurons, or dynamic stoichiometries exist in their assembly depending on relative subunit expression and other factors^4–5^. The P2X2/3 heterotrimers are widely expressed in sensory and autonomic neurons and are involved in multiple physiological and pathological processes, including taste perception, bladder reflexes, and visceral hypersensitivity^2, 6–14^. They are considered as an early mediator of nociceptive signaling^15^. Antagonists such as Gefapixant (AF-219) have demonstrated efficacy in treating refractory chronic cough (RCC), but their inhibition of P2X2/3 receptors also causes taste disturbances^16–17^. Consequently, P2X2/3 heterotrimers have attracted increasing attention.

A mechanistic analysis of P2X2/3 heterotrimers is required because of their biophysical differences from P2X3 homotrimers. Both receptor types are activated by ATP and its analogues but differ in ligand sensitivity: P2X3 homotrimers recognize both MgATP^2-^ and ATP⁴⁻, whereas P2X2 homotrimers respond preferentially to ATP⁴⁻ (ref.18). In heterotrimers, ATP-binding sites are located at heteromeric interfaces, suggesting a distinct mechanism of ligand discrimination that remains unresolved. P2X3 homotrimers contain three symmetric binding sites, whereas P2X2/3 heterotrimers have asymmetric sites^2^. P2X2/3 heterotrimers also exhibit slower desensitization kinetics and more sustained activity in the presence of Ca^2+^ compared with P2X3 homotrimers^19^. Incorporation of P2X2 subunits alters key gating determinants, including pore-lining residues, ion permeation pathways, and residues involved in desensitization^2^. Other ligand-gated ion channels, such as GABA_A_, glutamate, and nicotinic acetylcholine receptors, and have been of tremendous value in supporting pharmacological development^20–21^. P2X2/3 heterotrimers similarly differ from P2X3 homotrimers in multiple functional properties^5, 22–24^. Understanding these mechanisms is important for elucidating their physiological and pathological roles and for developing selective therapeutics.

Despite extensive studies on structure resolution, existing structures across multiple subtypes (P2X1, 2, 3, 4, 7) represent only homomeric architectures (P2X1, 2, 3, 4, 7) represent only homomeric architectures^25–42^. Heterotrimeric assemblies have remained largely uncharacterized due to challenges in protein production and high-resolution analysis. Therefore, the three-dimensional (3D) architecture of native heterotrimers such as P2X2/3 remains unknown, limiting mechanistic understanding and therapeutic development of the P2X channels. In this study, we use cryo-EM combined with functional and computational approaches to determine the structures and possible gating mechanisms of human P2X2/3 heterotrimers in ATP-bound P2X332 (2.68 Å) and P2X223 (2.95 Å) states, the *apo* P2X332 form (2.87 Å), and the antagonist- and ATP-bound closed conformation (3.3 Å). Endogenous expression and stoichiometry in dorsal root ganglion (DRG) neurons are confirmed. By integrating concatemer engineering, covalent modification, fluorescent noncanonical amino acid (flUAA) incorporation, voltage-clamp fluorometry (VCF), and membrane protein molecular dynamics (MD) simulations, we characterize ATP recognition, ligand specificity, ion permeation, and gating behaviors. Our findings provide a structural and mechanistic framework for P2X2/3 heterotrimer function.

## Results

### Cryo-EM structures of two assembly forms of ATP-bound human P2X2/3 heterotrimers: P2X332 and P2X322

The P2X2/3 heterotrimer consists of three subunits assembled from two distinct subtypes. Heterotrimer formation occurs stochastically in the expression system and is accompanied by the formation of P2X2 and P2X3 homotrimers, complicating sample purification and structural analysis. The high sequence identity between P2X2 and P2X3 (∼45%, Fig. S1) also introduces pseudo-symmetry, making subunit assignment challenging during post analysis. To address these issues, we used a BacMam mammalian expression system and a dual-tag, sequential affinity purification strategy to obtain highly purified human P2X2/3 (hP2X2/3) heterotrimeric proteins for cryo-EM analysis (Fig. 1a).

**Figure 1.**
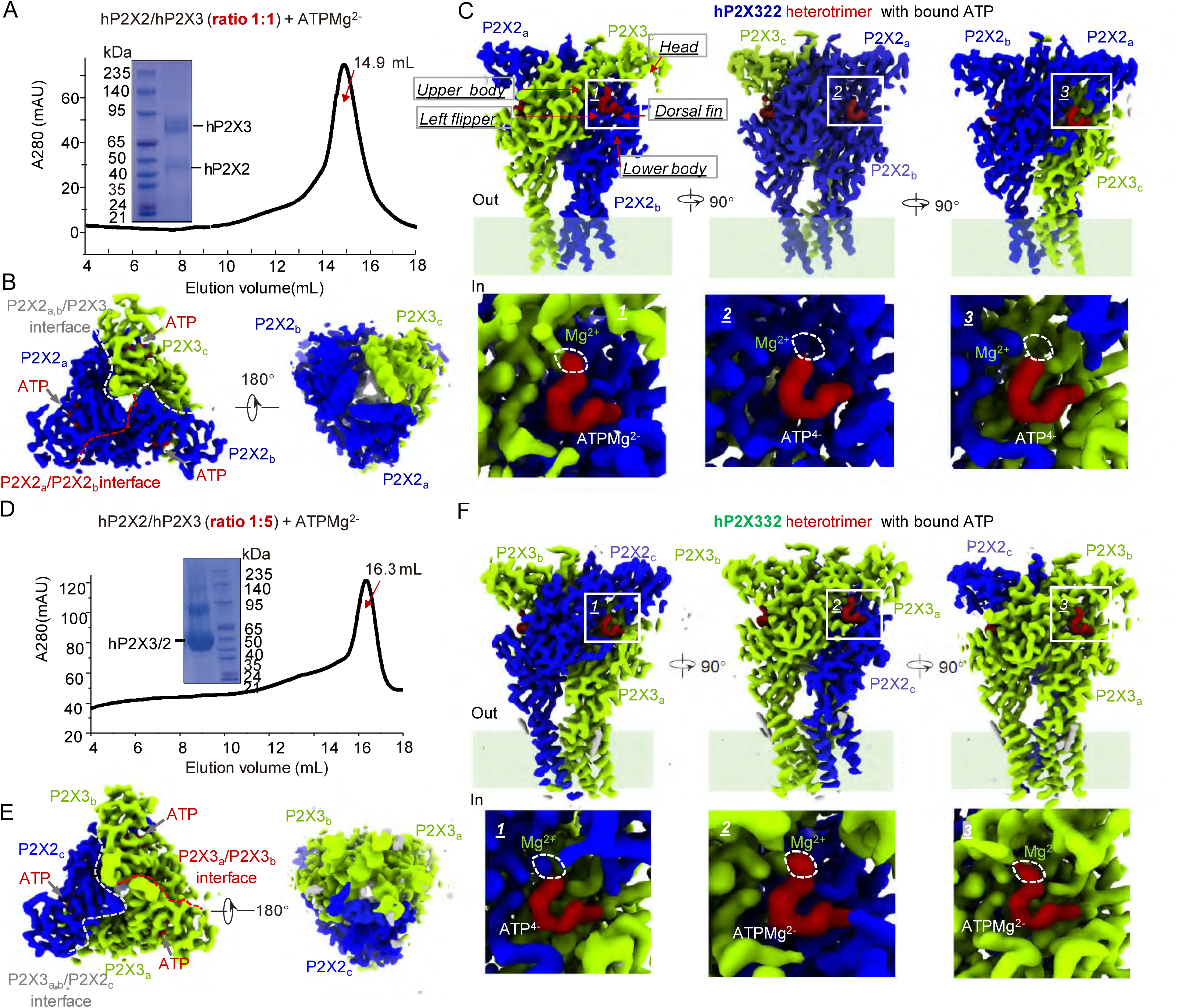
Cryo-EM structures of human P2X2/3 heterotrimers with bound ATP. (**A**) Size-exclusion chromatography (SEC) profile and SDS-PAGE analysis of the purified P2X2/P2X3 heterotrimeric complex co-expressed at a 1:1 ratio in the presence of ATP and Mg²⁺. (**B, C**) Cryo-EM density map of the hP2X322 heterotrimer bound to ATP, shown from the membrane plane (B, left), cytoplasmic side (B, right), and parallel to the membrane plane (C, upper). Insets in (C, lower) highlight the three orthosteric sites, illustrating the ATP binding conformations. (**D**) SEC profile and SDS-PAGE analysis of the P2X2/P2X3 heterotrimer co-expressed at a 1:5 ratio with ATP and Mg²⁺. (**E, F**) Cryo-EM density map of the hP2X332 heterotrimer bound to ATP, displayed from the membrane plane (E, left), cytoplasmic side (E, right), and parallel to the membrane plane (F, upper). Insets in (F, lower) show the ATP-bound orthosteric sites. In all cryo-EM maps, P2X2 and P2X3 subunits are colored blue and green, respectively, and ATP molecules are shown in red.

For structural determination, hP2X2 and hP2X3 were co-expressed in HEK293S GnTI⁻ cells to obtain heterotrimeric hP2X2/3 protein complexes. To improve protein expression and minimize glycosylation-induced heterogeneity, the C-terminus of hP2X2 after S390 was truncated to produce ΔhP2X2-M1–S390 (Figs. S1 and S2A), and a C-terminal twin-Strep II tag was introduced to facilitate purification and detection. Electrophysiological analyses confirmed that ATP affinity and desensitization kinetics were preserved relative to those of the full-length subunits (Fig. S2A, B, E). Similarly, guided by a previous structural study on hP2X3 ^26^, a truncated hP2X3 construct (hP2X3-MFCslow (hP2X3-MFC-T13P/S15V/V16I): D6–T364 with N- and C-terminal truncations and three N-terminal substitutions) was employed, incorporating a C-terminal FLAG tag. Functional characterization indicated unaltered ATP affinity and heightened sensitivity to allosteric modulators such as AF-353 in hP2X3-MFCslow, while current kinetics shifted from rapid to slow desensitization (Fig. S2C, D, E), confirming retained functionality.

High-resolution cryo-EM structures of multiple hP2X2/3–ATP complexes were determined using single-particle analysis (Table S1; Figs. S3–S4). To overcome the inherent pseudo-symmetry associated with P2X2/3 heterotrimer reconstruction, we implemented a multi-round local 3D classification approach, effectively mitigating symmetry-induced averaging and enabling isolation of authentic asymmetric conformations (Fig. S3–S4; see Methods). Within the Aspartate (D) and Glutamate (E)-enriched region, designated here as the *D/E*-loop (see below), we observed distinct structural divergence between subunits: P2X3 displays partial β-strand disruption into loop conformations, whereas P2X2 predominantly maintains an intact β-sheet structure. This differential structural feature served as a reliable determinant for subunit assignment during density interpretation (see Methods).

Intriguingly, cryo-EM analysis of hP2X2/3–ATP complexes under varying co-expression ratios revealed two discrete stoichiometric assemblies: hP2X322/ATP at 2.95 Å (Fig. 1A–C) and hP2X332/ATP at 2.87 Å (Fig. 1D–F). At an equimolar ratio of hP2X2:hP2X3 (1:1), only the hP2X322 assembly was observed (Fig. 1A), whereas a 1:10 ratio yielded exclusively the hP2X332 assembly (see Methods). At an intermediate ratio of 1:5 (Fig. 1D), both assemblies coexisted within the particle population and were computationally separated to yield distinct reconstructions of hP2X332/ATP (Fig. 1E, F) and hP2X322/ATP complexes (Fig. S3). These results indicate that heterotrimer assembly is dynamic and dependent on the relative abundance of the two subunits. hP2X2 appears to favor P2X322 formation at lower abundance/ratio, whereas higher levels of hP2X3 are required to generate P2X332.

To assess ATP binding and heterotrimer stability, 2.2-μs conventional molecular dynamics (CMD) simulations were conducted for hP2X322/ATP complex (Figs. S5–S7). The simulations substantiated the complex’s stability, as reflected in consistent root mean square deviation (RMSD) values for both protein and ATP, complemented by ATP dihedral angle analyses (Figs. S5B, C; S6B, C; S7B, C). Key residues interacting with the three asymmetrically arranged ATP molecules remained consistently engaged throughout the simulation (Figs. S5A, C; S6A, C; S7A, C). A longer 4.2-μs simulation for hP2X332/ATP (Figs. S9–S11) confirmed analogous stability: RMSD and dihedral analyses (Figs. S8B, C; S9B, C; S10B, C) as well as the positioning of asymmetrically arranged ATP-interacting residues (Figs. S8A; S9A; S10A). Overall, these results indicate that both heterotrimeric assemblies adopt stable, non-transient conformations suitable for mechanistic and functional analyses.

### Differential recognitions of endogenous MgATP²⁻ and ATP⁴⁻ by hP2X332 and hP2X322 heterotrimeric assemblies

The ability of hP2X332 and hP2X322 assemblies to distinguish between MgATP²⁻ and ATP⁴⁻ represents a fundamental feature of P2X receptor function. Both ligands are physiologically relevant, and P2X receptor subtypes segregate according to ligand sensitivity: P2X1 and P2X3 are responsive to both MgATP²⁻ and ATP⁴⁻, whereas P2X2, P2X4, and P2X7 display selective responsiveness to ATP⁴⁻ (ref.18). Accumulating evidence indicates that, despite simultaneous occupation of the orthosteric binding pocket, MgATP²⁻ and ATP⁴⁻ can induce distinct downstream effects via subtle differences in their coordination environments, analogous to biased agonism in GPCR systems^39, 43–44^. The hP2X2/3 heterotrimerization of subunits from these two ligand-sensitivity classes introduces pronounced structural and functional heterogeneity. As a result, ligand recognition in hP2X332 and hP2X322 assemblies cannot be predicted solely based on homomeric receptor properties, necessitating direct structural and functional interrogation.

Interestingly, cryo-EM structures of ATP-bound human P2X2/3 heterotrimers reveal a clear and reproducible architectural logic governing MgATP²⁻ and ATP⁴⁻ binding. In hP2X322 (Fig. 1C), three structurally distinct ATP-binding sites are delineated: (i) a hybrid site comprising the head and upper body domains of P2X3 together with the lower body and partial dorsal fin and left flipper regions of P2X2; (ii) a fully P2X2-derived site encompassing the head, upper body, lower body, dorsal fin, and left flipper; and (iii) a second hybrid site formed by the head and upper body of P2X2 in conjunction with the lower body and associated structural elements of P2X3. The hP2X332 assembly exhibits a complementary configuration (Fig.1F), with these domain contributions reciprocally arranged. This structural heterogeneity directly dictates ligand selectivity. Binding pockets incorporating P2X3-derived head and upper body domains preferentially coordinate MgATP²⁻, as evidenced by pronounced Mg²⁺ densities consistent with prior structural analyses of P2X homotrimers (Fig. 1C, left; Fig. 1F, middle and right). In contrast, pockets lacking these domains preferentially accommodate ATP⁴⁻ (Fig. 1C, middle and right; Fig. 1F, left). Accordingly, hP2X332 binds two MgATP²⁻ and one ATP⁴⁻, whereas hP2X322 binds one MgATP²⁻ and two ATP⁴⁻. The functional consequences of this asymmetric ligand occupancy under physiological and pathological conditions remain to be elucidated.

### Human, rat, and mouse P2X3/2 heterotrimers exhibit dynamic stoichiometric plasticity across heterologous expressions, native DRG neurons, and inflammatory states *in vivo*

Differential recognition of MgATP²⁻ and ATP⁴⁻ (Fig.1), together with structural differences in the D/E-loops of P2X2 and P2X3 (see below), allows discrimination of hP2X2/3 heterotrimer stoichiometry. However, current understanding of their dynamic assembly is largely derived from truncated constructs (ΔhP2X2-M1–S390 and ΔhP2X3-MFCslow) examined under detergent-solubilized conditions, raising uncertainty regarding their physiological relevance in intact cellular and in vivo environments. Co-expression of ΔhP2X2 and ΔhP2X3 in HEK293 cells at defined ratios (1:1 and 1:5) (Fig. S2F–G) revealed P2X322 currents, as well as currents from P2X332 and homomeric P2X3. Stoichiometric classification was based on pharmacological and electrophysiological criteria: receptors unresponsive to low ATP concentrations (0.1–0.3 μM) and insensitive to AF-353 inhibition were assigned as hP2X322, whereas responsive receptors were categorized as homomeric hP2X3 or heteromeric hP2X332. These were further distinguished by current kinetics at 1 μM ATP, with hP2X332 exhibiting intermediate properties between homomeric hP2X3 and homomeric P2X2 channels. These results demonstrate that heterotrimer assembly, including P2X332 and P2X322, occurs under both detergent-solubilized structural and live-cell conditions, although P2X332 has previously been considered the predominant heteromeric form^10, 45–47^.

P2X3 homomeric and P2X2/3 heteromeric channels are endogenously expressed in DRG neurons. DRG neurons from wild-type (WT) mice were here analyzed under basal physiological conditions and following inflammatory stimulation (Fig. 2A). Quantitative real-time PCR (qPCR) indicated that *P2rx3* expression exceeds *P2rx2* under normal conditions (1.46 ± 0.17 vs. 0.0783 ± 0.0092, p < 0.0001, unpaired t-test, n = 7; Fig. 2B). Inflammatory challenge significantly increased the *P2rx3*/*P2rx2* expression ratio (23.8 ± 4.2 vs. 38.1 ± 5.3, p < 0.05, unpaired t-test, n = 13; Fig. 2C), suggesting dynamic regulation of subunit stoichiometry in physiological and pathological contexts.

**Figure 2.**
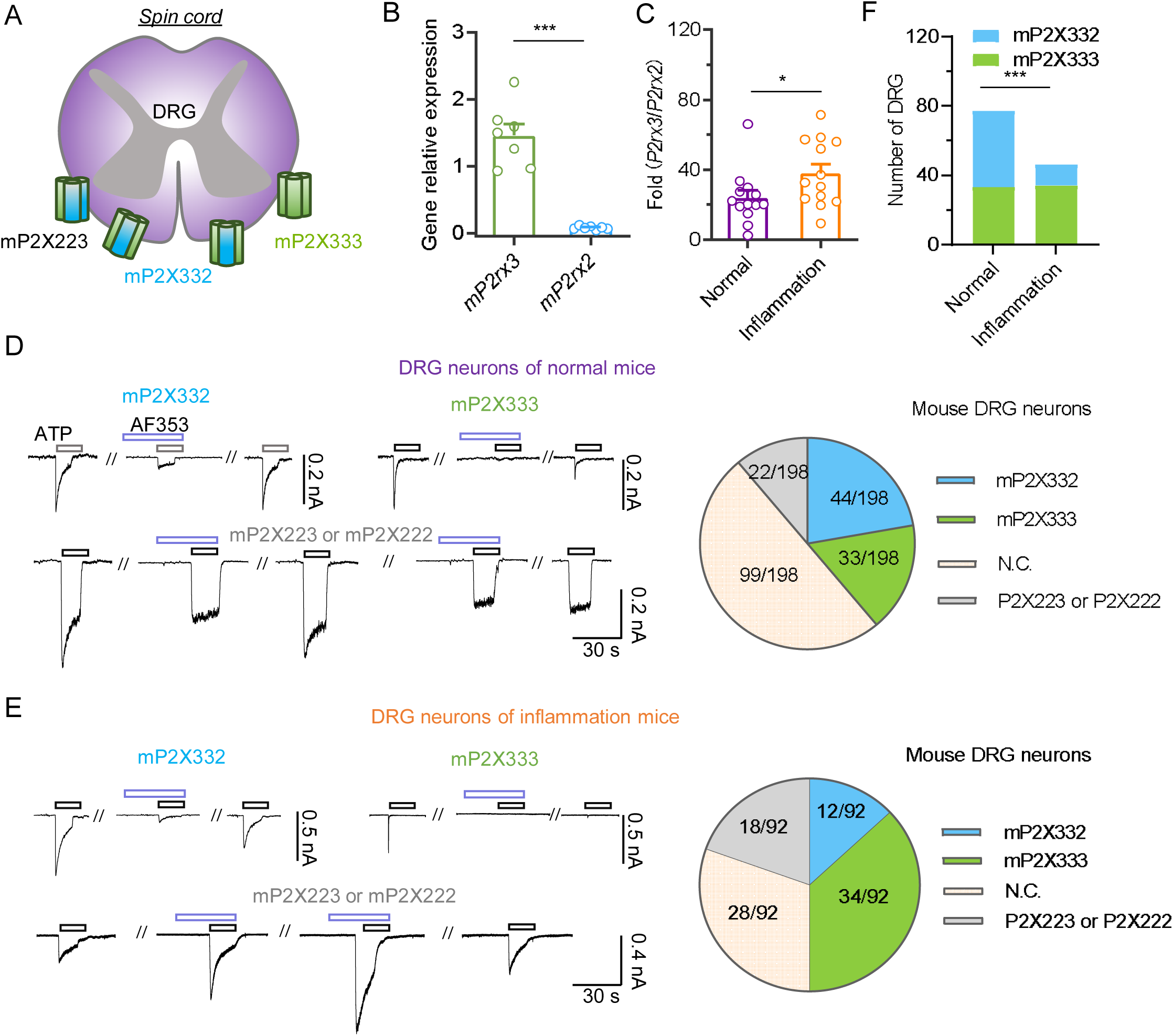
Distribution of P2X2 and P2X3 homo- and heterotrimers in mouse dorsal root ganglia under physiological and inflammatory conditions. (**A**) Schematic representation of mouse dorsal root ganglion (DRG) structure. (**B**) Expression levels of *P2rx3* and *P2rx2* genes in DRG of naïve mice. Data were analyzed using an unpaired two-tailed t-test (***p < 0.001).(**C)** Ratio of *P2rx3* to *P2rx2* expression in DRG of normal versus inflamed mice; unpaired two-tailed t-test (*p < 0.05).(**D, E**) Proportions of P2X3 homotrimers, P2X332 heterotrimers, P2X2 homotrimers, and P2X322 heterotrimers in DRG neurons of normal (D) and inflamed (E) mice. Homo- and heterotrimers were distinguished based on ATP-evoked current shapes and sensitivity to AF-353 (D and E, left). Pie charts depict the number of neurons expressing each channel type (D and E, right). N.C., no current detected. (**F**) The prevalence of P2X332 heterotrimers was significantly elevated in DRG neurons of inflamed mice compared with normal controls (Chi-squared test, ***p < 0.001).

DRG neurons were harvested from mice, cultured for 16–20 hours, and analyzed electrophysiologically in the presence of AF-353, which selectively inhibits ATP currents mediated by homomeric mP2X3 and heteromeric mP2X332 channels (Fig. 2D-E). Combining pharmacological inhibition with current desensitization kinetics enabled the discrimination of mP2X3 homotrimers, mP2X332 heterotrimers, and AF-353-insensitive mP2X322 heterotrimers. Across both basal and inflammatory conditions, mP2X3 homotrimers and mP2X332 predominated, whereas mP2X322 represented a minor fraction (∼10–20%) of neurons (Fig. 2D–F). Under homeostatic conditions, mP2X332 homotrimers and mP2X3 homotrimers were similarly represented (22.2% vs. 16.7%; Fig. 2D, F), while inflammation favored mP2X3 homotrimers (13.0% vs. 37.0%; Fig. 2E, F; p < 0.001), consistent with qPCR measurements (Fig. 2C). These data confirm that all three receptor assemblies are expressed in DRG neurons and exhibit dynamic regulation. Notably, the mP2X322 population emerges even when *P2rx2* expression is much lower than *P2rx3*.

To further validate the dynamic assembly of P2X2/3 heterotrimers observed in human and mouse systems, molecular approaches, including P2X2/3 heteromeric concatemers and co-transfection of full-length P2X2 and P2X3 cDNAs in HEK293 cells, are required. This framework allows investigation of full-length heteromeric formation, the stoichiometry of MgATP²⁻ and ATP⁴⁻ binding at the three asymmetric orthosteric sites, and the interaction of allosteric modulators at the asymmetric allosteric sites (see below). Rat-derived sequences are advantageous for assembling rP2X332, rP2X322, and rP2X222 concatemers. Evolutionary analysis shows that P2X2 and P2X3 genes are highly conserved across human, mouse, and rat, with substantial sequence homology (Fig. S11A). Functional characterization of ATP and allosteric ligand recognition (ATP, AF-219, AF-353) in both homomeric P2X3 and heteromeric P2X2/3 channels reveals no significant interspecies differences (EC_50_ of ATP on hP2X3, rP2X3, mP2X3: 508 ± 84, 396 ± 68, 258 ± 22 nM; IC_50_ of AF-219: 197 ± 43, 205 ± 43, 93.6 ± 4.8 nM; IC_50_ of AF-353: 8.68 ± 2.22, 8.80 ± 0.42, 10.8 ± 1.1 nM; EC_50_ of ATP on hP2X2/3 and rP2X2/3: 3.72 ± 0.12 and 1.24 ± 0.45 μM; IC_50_ of AF-219: 0.772 ± 0.319 and 1.11 ± 0.29 μM; IC_50_ of AF-353: 0.276 ± 0.157 and 0.165 ± 0.045 μM, n = 3–4; Fig. S11B–G). These results support the functional interchangeability of these orthologs. The high degree of sequence conservation and functional similarity across species permits their interchangeable use, supporting experimental consistency and reproducibility.

Then, full-length human and rat P2X2 and P2X3 subunits were co-transfected at a 1:2 ratio (h/rP2X2:h/rP2X3, Fig. S12) to replicate elevated in vivo P2X3 mRNA expression. ATP concentration-response relationships were assessed under continuous AF-353 application to prevent formation of h/rP2X3 homotrimers and h/rP2X332 heterotrimers, ensuring selective activation of h/rP2X322 heterotrimers. Both human and rat subunits assembled into P2X322 heterotrimers at measurable frequencies (human: 2/5; rat: 4/11; Fig. S12A–C), with EC_50_ values lower than those of h/rP2X2 homotrimers (EC_50_ of ATP (μM): h/rP2X322: 7.15 ± 0.29 and 7.63 ± 0.73; h/rP2X2: 15.1 ± 3.7 and 16.1 ± 1.5; n = 5–11; Fig. S12A–C).

To further corroborate these observations, we engineered an rP2X2-rP2X2-rP2X3 concatemer (rP2X2-2-3; Fig. S12D), positioning the three subunits adjacently to favor near-exclusive assembly into rP2X3-2-2 heterotrimers. The rP2X2-2-3 concatemer exhibited full activation upon 100 μM ATP stimulation (pA/pF = 65.9 ± 4.5, n = 4; Fig. S12E), with ATP affinity shifted leftward relative to rP2X2-2-2 concatemers (EC_50_ = 25.4 ± 3.8 μM vs 35.9 ± 3.2 μM; rP2X2 EC_50_ = 11.6 ± 0.6 μM, n = 3; Fig. S12F). Application of 10 μM AF-353 failed to inhibit ATP-evoked currents, and even at 100 μM, maximal inhibition reached only 47.3 ± 2.8%. AF-353 sensitivity was not significantly different between rP2X2-2-3 and rP2X2-2-2 concatemers (p > 0.05; two-way ANOVA with Sidak’s multiple comparisons, F(2,13) = 0.9072, n = 3–4; Fig. S12G–H). These data show that P2X322 heterotrimers, detected in detergent-mediated protein purification (Fig. 1A), can also form from full-length P2X2 and P2X3 sequences in HEK293 *in vitro* expression systems.

Thus, *in vivo* P2X2/3 heterotrimer assembly is modulated by P2X2 and P2X3 expression levels, resulting in the coexistence of P2X332 and P2X322 heterotrimers. Even low P2X2 expression is sufficient to drive P2X322 formation. Pathological ATP elevation and increased *P2rx3* transcription promote the generation of additional P2X3 homotrimers to facilitate rapid nociceptive signaling, whereas under physiological conditions, signaling is primarily mediated by P2X2/3 heterotrimer and P2X3 homotrimers.

### Cryo-EM structure of *apo* hP2X332 heterotrimers

Considering that P2X332 constitutes the major endogenous form of P2X3/2 heterotrimers in vivo, a predominance resulting from the 20–30-fold higher expression of P2X3 compared with P2X2 (Fig. 2B), we next investigated the structural and functional mechanisms that dictate this assembly. The cryo-EM structure of *apo* hP2X332 was determined at 2.87 Å resolution (Fig. 3A–B) using a BacMam mammalian expression system to co-express hP2X2 and hP2X3 at a 1:10 ratio (Figs. 3A, B and S13). CMD simulations (∼2.7 μs) confirmed the stability of the assembly (Fig. S14). Each subunit contains extracellular domains, including the head, dorsal fin, and left flipper, as well as transmembrane (TM) domains (Fig. 3C, left). The channel gate comprises two distinct constriction sites (Fig. 4A): an extracellular-facing gate formed by I343 (hP2X2) and I323 (hP2X3 ×2), and a cytoplasm-facing gate defined by T350 (hP2X2) and T330 (hP2X3 ×2), with pore radii of 0.78 Å and 0.63 Å, respectively. These dimensions preclude hydrated Na^+^ /Ca^2+^ permeation, indicating a closed pore. Despite minor residue differences in the pore region (lower middle panels of Fig. S15), the gating architecture (Fig. 4A) is consistent with that of homotrimeric P2X receptors.

**Figure 3.**
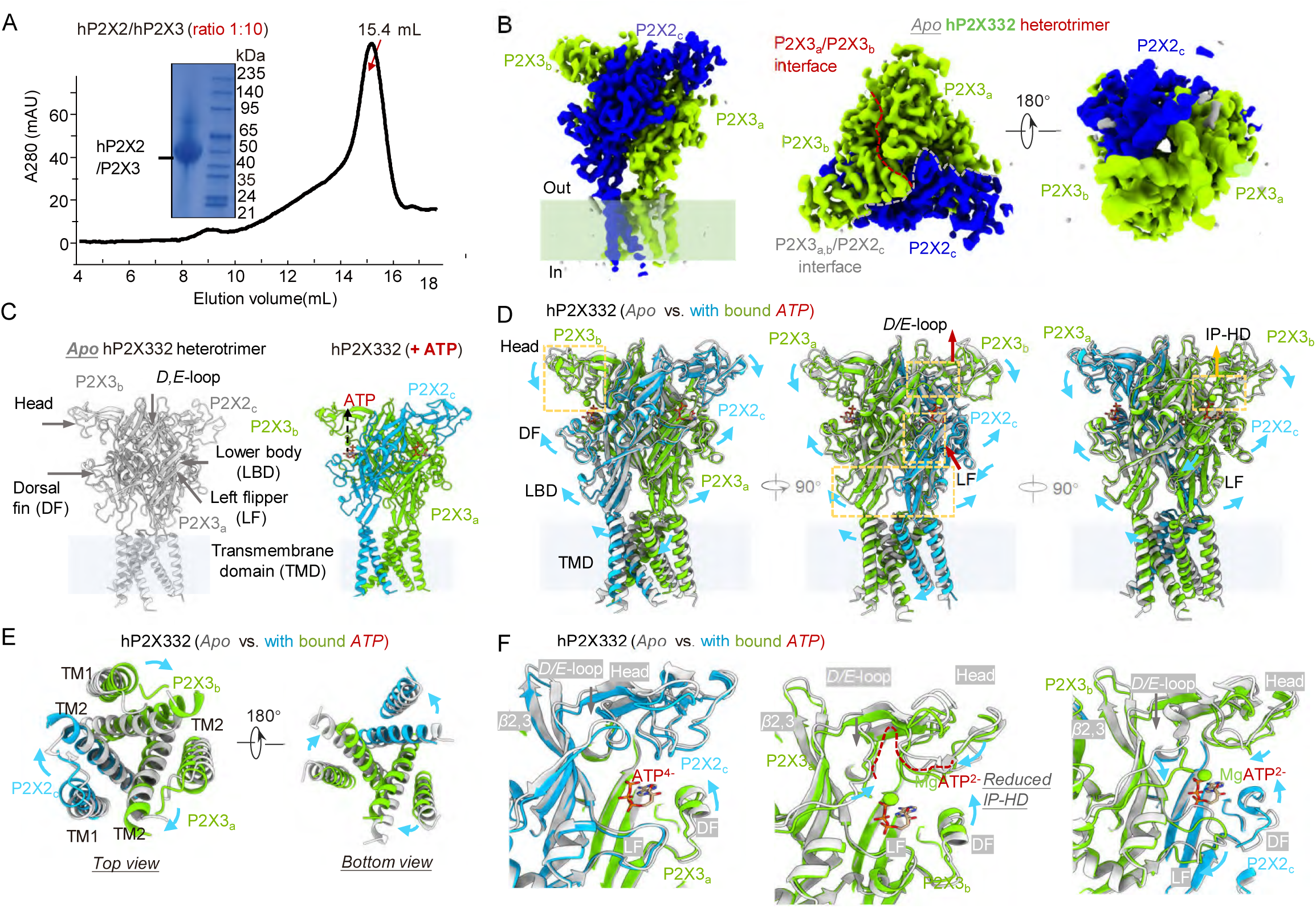
Cryo-EM structures of apo and ATP-bound hP2X332 heterotrimers and structural superimposition. **(A)** SEC profile and SDS-PAGE analysis of the purified P2X2/P2X3 heteromer co-expressed at a 1:10 ratio. **(B)** Cryo-EM density map of the hP2X322 heterotrimer, viewed parallel to the membrane plane (left), from the membrane plane (middle), and from the cytoplasmic side (right). Red and gray dashed lines indicate inter-subunit interfaces between P2X3_a_–P2X3_b_ and P2X3_a,b_–P2X2_c_, respectively. **(C)** Comparison of apo (gray) and ATP-bound (blue and green) hP2X332 structures. **(D–F)** Superimposition of apo and ATP-bound structures, shown parallel to the membrane plane (D), with close-up views of transmembrane domains 1 and 2 (TM1, TM2) (E), and orthosteric binding sites (F). Blue arrows indicate conformational changes from apo to ATP-bound states. LBD, ligand-binding domain; IP-HD, inner pocket of the head domain.

**Figure 4.**
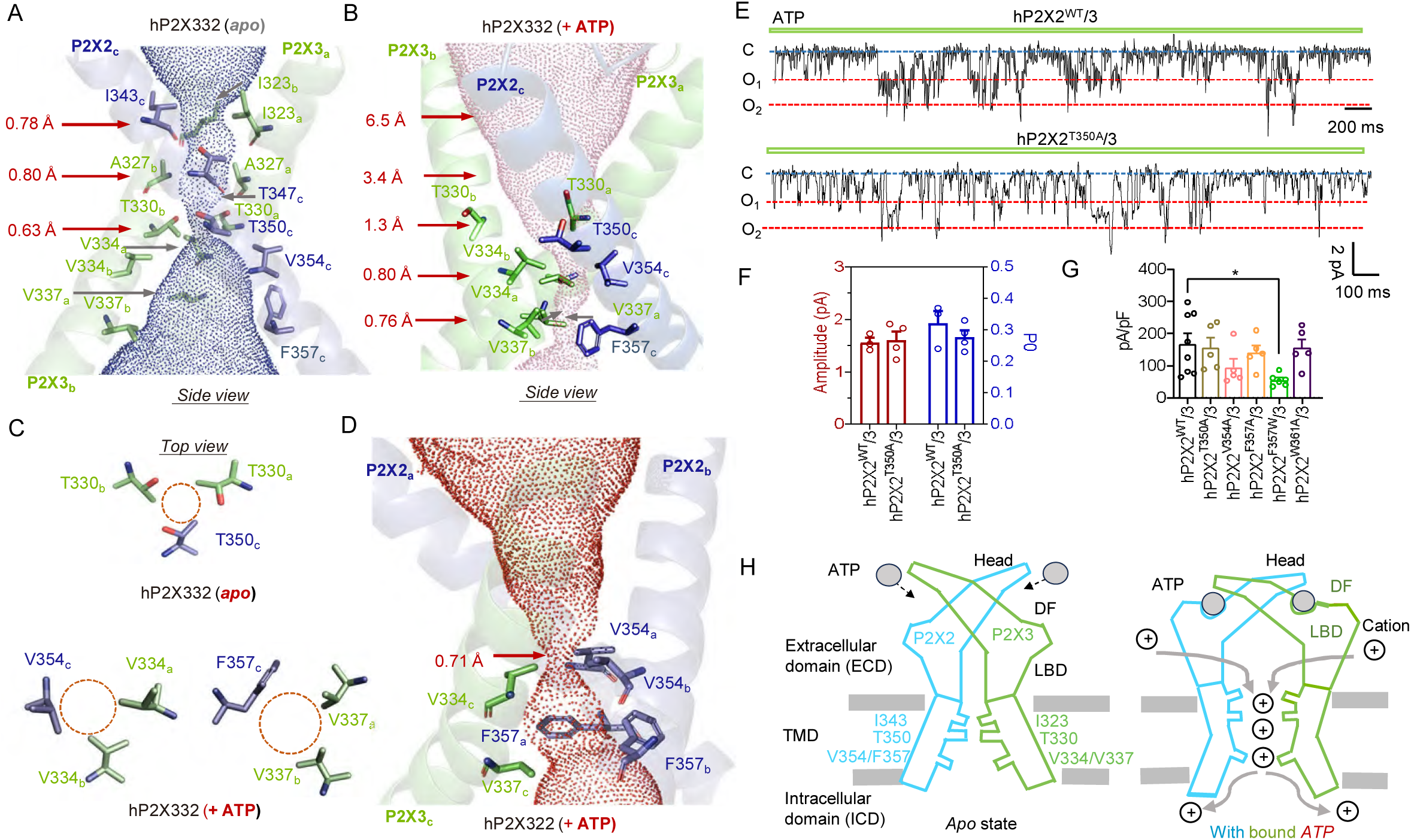
Key pore residues and functional analysis of hP2X332 heterotrimer during activation and desensitization. **(A, B)** Pore structures of the *apo* (A) and ATP-bound (B) states of hP2X332 heterotrimer. Blue (*apo*) and brick-red (ATP-bound) dots denote the pore-lining surface of hP2X3, with critical residues highlighted as sticks. Pore dimensions were calculated using MOLE. **(C)** Representation of the desensitization gate of hP2X332. Critical residues are highlighted as sticks. **(D)** Pore structure of the ATP-bound hP2X322 heterotrimer. Key residues are highlighted as sticks. **(E, F)** Unitary currents recorded from outside-out patches held at −60 mV in response to 0.1 μM ATP (E) and summary of amplitude and open probability (P₀) (F). Red and blue lines indicate channel opening (O) and closing (C), respectively. Each circle represents an independent experimental measurement. **(G)** Current density of hP2X332 and indicated mutants. Data points correspond to independent experiments. *p < 0.05, one-way ANOVA with Dunnett’s multiple comparison test (F(5,28) = 2.669). **(H)** Schematic cartoon summarizing the activation and ion permeation mechanisms of the hP2X332 heterotrimer.

As the *apo* hP2X3-3-2 cryo-EM structure was determined using truncated constructs (ΔhP2X2-M1–S390 and ΔhP2X3-MFCslow), potential effects of truncation on assembly heterogeneity were evaluated. Two pairs of engineered disulfide bonds were introduced at the dorsal fin/left flipper domain interfaces between hP2X2 and adjacent hP2X3 subunits in full-length hP2X332, generating hP2X3^L195C^/hP2X2^A295C^ and hP2X3^S275C^/hP2X2^I215C^ mutants (Fig. S16A). Non-reducing SDS–PAGE showed prominent trimeric bands, indicating successful disulfide bond formation (Fig. S16B). Electrophysiological recordings revealed reduced currents for hP2X3^S275C^/hP2X2^I215C^ compared to WT, which increased ∼2–5-fold following DTT treatment (current ratio (DTT/H_2_O_2_) = 5.07 ± 0.31, 1.89 ± 0.26, 1.01 ± 0.11 and 1.72 ± 0.53 for hP2X2^I215C^/3^S275C^, hP2X2^WT^/3^S275C^, hP2X2^I215C^/3^WT^ and hP2X2/3^WT^, respectively; p < 0.0001, hP2X2^I215C^/3^S275C^ vs. WT, p > 0.05, hP2X2^WT^/3^S275C^ and hP2X2^I215C^/3^WT^ vs. WT, one-way ANOVA with Dunnett’s multiple comparisons test, F (6, 16) = 13.00, n = 3-5; Fig. S16C-D). For hP2X3^L195C^/hP2X2^A295C^, currents increased 2–3-fold, while DTT slightly reduced WT currents (ratio = 2.95 ± 0.73, 1.31 ± 0.23, 1.12 ± 0.005 and 1.72 ± 0.53 for hP2X2^A295C^/3^L195C^, hP2X2^WT^/3^L195C^, hP2X2^A295C^/3^WT^ and hP2X2/3^WT^, respectively; p < 0.1, hP2X2^A295C^/3^L195C^ vs. WT, p > 0.05, hP2X2^WT^/3^L195C^ and hP2X2^A295C^/3^WT^ vs. WT, one-way ANOVA with Dunnett’s multiple comparisons test, F (6, 16) = 13.00, n = 3-5; Fig. S16C-D). These results indicate that the *apo* hP2X332 assembly is structurally valid and suggest that intersubunit relative movement at the P2X2–P2X3 interface is required for channel gating.

We next characterized the interfacial residues between P2X3 and P2X2 in the *apo* hP2X332 cryo-EM structure. In comparison with the P2X3 homotrimer, replacement of one P2X3 subunit with P2X2 introduces several non-conserved residues at the interface, including F173, L154, L147, and M191 in the head domain, and R286, I328, and H330 in the lower body domain of hP2X2 (Fig. S15). These interfacial differences may influence the coexistence and relative proportions of hP2X332 heterotrimers, hP2X3 and hP2X2 homotrimers, and hP2X322 heterotrimers in DRG neurons co-expressing both subunits. Differences in expression levels and neuronal distribution may further modulate these assembly ratios. The *apo* hP2X332 structure thus provides atomic-level insight into P2X3/2 heterotrimer composition and a structural basis for their dynamic assembly.

### Comparison of the *apo* and ATP-bound hP2X332 cryo-EM structures reveals asymmetric assembly while maintaining conserved symmetric gating

Upon resolving the *apo* hP2X332 structure, it is possible to compare it with the ATP-bound hP2X332 cryo-EM structure. The ATP-bound conformation appears to correspond to a desensitized state (Fig. 4A, B). In the desensitization configuration, the two constriction sites—the extracellular gate formed by I343 (hP2X2) and I323 (hP2X3 ×2), and the intracellular gate formed by T350 (hP2X2) and T330 (hP2X3 ×2)—are expanded to pore radii of 6.5 Å and 1.3-1.5 Å (Fig. 4B), respectively, allowing hydrated Na⁺ and Ca²⁺ permeation. In contrast, a desensitization gate, reminiscent of those observed in other P2X homotrimers, is established in the ATP-bound state (Fig. 4B, C). This gate, primarily composed of V354 (hP2X2) and V334 (hP2X3 ×2), and F357 (hP2X2) and V337 (hP2X3 ×2), exhibits pore radii of 0.76–0.8 Å (Fig. 4B, C), effectively blocking ion conduction.

Nonetheless, the comparative analysis of *apo* and ATP-bound hP2X332 cryo-EM structures provides essential mechanistic insights into hP2X332 gating, including both activation and desensitization. This inference is founded upon two critical observations: (1) P2X receptor subtypes maintain highly conserved gating architectures^25–42^, as demonstrated by high-resolution homotrimeric structures of P2X1, 2, 3, 4, and 7 in diverse functional states; and (2) detailed structural comparisons of desensitized and open states of those homotrimer structures^25–42^ indicate that, aside from desensitization gate closure and minor inward contraction of the TM region, ATP-driven extracellular domain (ECD) conformational rearrangements are fundamentally conserved^26–28, 31, 33, 39, 42^. Consistent with this framework, structural superposition of the ATP-bound hP2X3332 cryo-EM structure with the ATP-bound open-state hP2X3 homotrimer (5VJK) provides compelling corroboration (Fig. S17A). The ECD are virtually indistinguishable between the desensitized hP2X332 and the open hP2X3 homotrimer, as evidenced by the precise alignment of pore constriction sites within the ECD (Fig. S17B, middle). The principal structural divergence is confined to regions below the gate, corresponding to the desensitization transition (Fig. S17B, middle).

Superposition of the *apo* and ATP-bound hP2X332 conformations, together with prior analyses, reveals gating and desensitization mechanisms that mirror those of P2X homotrimers^26–27, 42^ (Fig. 3C–F). ATP binding at intersubunit pockets, despite heterogeneity, initiates downward displacement of the head domain. Upward motion of the dorsal fin domain closes the ATP-binding clefts (Fig. 3C, D) and propagates conformational changes that bend the lower body domain outward (Fig. 3D). The β-sheet layer of the lower body, directly coupled to the TM helices, transmits this motion as torque, producing a clockwise “twist-to-open” rotation of TM1 and TM2, thereby enabling the transition from resting to open conformation (Figs. 3D, E and 4B). The desensitization gate remains open during initial channel opening but closes upon sustained activation, a process previously documented in P2X3 and P2X1^26, 42^.

During desensitization, T350 (hP2X2) and T330 (hP2X3×2) largely maintain a symmetric, near–ion-conducting conformation (Fig. 4C, upper), whereas the residues forming the narrowest constriction—V354 (hP2X2) and V334 (hP2X3 ×2), and F357 (hP2X2) and V337 (hP2X3 ×2)—adopt an asymmetric orientation (Fig. 4C, lower). This indicates that asymmetrically arranged amino acids can support normal channel desensitization. A similar closure of the desensitization gate, mediated by V334 (hP2X3) and V354 (hP2X2 ×2), and V337 (hP2X3) and F357 (hP2X2 ×2), is also observed in the heterotrimeric ATP-bound hP2X322 structure (Fig. 4D).

Owing to the heterogeneity and asymmetry of the amino acid composition within the hP2X332 pore, we further examined the effect of incorporating P2X2 subunit residues on channel gating. Substitution of T350 with alanine (hP2X2^T350A^) did not disrupt pore integrity, and heterotrimer function remained comparable to WT hP2X332: single-channel conductance/amplitude (1.61 ± 0.2 vs. 1.56 ± 0.1 pA, p > 0.05), ATP-saturated open probability (P₀ = 0.278 ± 0.021 vs. 0.320 ± 0.04, p > 0.05), and whole-cell current density (156 ± 31 pA/pF vs. 170 ± 33, p > 0.05, n = 5, Fig. 4E–G). Alanine substitutions at V354, F357, and W361 also did not significantly affect ATP-evoked currents. These results indicate that pore heterogeneity is tolerated and that the hP2X332 pore retains its nonselective cation permeability, in contrast to the highly conserved amino acid requirements of voltage-gated Kv or Nav channels^48–49^.

Notably, augmentation of the F357 side chain via tryptophan substitution at the desensitization gate (hP2X2^F357W^/hP2X3) elicited a marked diminution in current density (pA/pF = 55.3 ± 7.5 versus 168 ± 33 for mutant vs.WT heterotrimers, p < 0.01, n = 6–8, one-way ANOVA with Dunnett’s multiple comparisons, F(5,28) = 2.522, Fig. 4G), highlighting its dual role as both a structural constituent of the desensitization gate and a modulator of pore constriction that limits ion conduction in the open channel.

In addition, despite the pronounced asymmetry of the three ATP-binding pockets, ATP-driven conformational rearrangements in the ECD—comprising inward displacements of the D/E-loop, coordinated motions of the left flipper and dorsal fin, and contraction of the inner pocket of the head domain (IP-HD) (Fig. 4H)—previously shown to be essential in P2X3 homotrimers^35, 50^, are largely retained in the ATP-bound hP2X332 heterotrimer, albeit with site-specific variations. Collectively, these data indicate that asymmetric subunit assembly does not compromise the overarching symmetric and conserved gating architecture.

### Activation of P2X332 heterotrimer is achieved through the engagement of merely two MgATP⁴⁻ molecules at orthosteric sites 2 and 3

Attention is subsequently directed toward the functional diversity conferred by the hP2X332 heterotrimeric assembly, including differences in ATP recognition, structural determinants of slower desensitization, and ion permeation via lateral pore pathways. These features provide a rationale for the physiological co-expression of homotrimeric and heterotrimeric P2X receptors. As delineated above, heteromeric hP2X332 differs from hP2X3 and hP2X2 homotrimers in its capacity to discriminate between MgATP²⁻ and ATP⁴⁻ (Fig.1). An outstanding question is whether activation of hP2X332 requires only two ATP molecules, as in hP2X2 homotrimers^51^, and whether these correspond to two MgATP²⁻ sites or involve a combination of MgATP²⁻ and ATP⁴⁻ binding at distinct sites.

We defined the three ATP-binding pockets of the hP2X332 heterotrimer as sites 1, 2 and 3 (Fig. 5A–C). Site 1 is formed at the interface between the P2X2 subunit’s head–upper body–left flipper and the P2X3 subunit’s lower body–dorsal fin (Fig. 5A), site 2 at the interface of the P2X3 head–upper body–left flipper and P2X2 lower body–dorsal fin (Fig. 5B), and site 3 lies between the two P2X3 subunits (Fig. 5C). While key ATP-recognition residues are conserved (Fig. 5A-C), important differences exist: P2X2-F174 versus P2X3-L198 increases ATP affinity approximately tenfold (Fig. 5A-C); D/E-loop conformations in P2X3 establish a D158-MgATP²⁻ binding mode distinct from P2X2-ATP⁴⁻(Fig. S17C-E), creating a unique coexistence pattern with head-domain residues; and hydrogen bonding between ATP’s β-phosphate and left-flipper residues P2X2-S296 (site 1) and P2X3-S275 (sites 2 and 3) is irregular, enhancing affinity by 3–5 fold. Overall, these findings demonstrate significant heterogeneity in MgATP²⁻/ATP⁴⁻ binding across the three pockets of the hP2X332 heterotrimer. A comparable pattern of ATP recognition (Fig. S18A–C) and the distinctive coexistence of D/E-loop and head-domain residues with MgATP²⁻ or ATP⁴⁻ (Fig. S18D–F) is also observed in ATP-bound hP2X322 heterotrimer.

**Figure 5.**
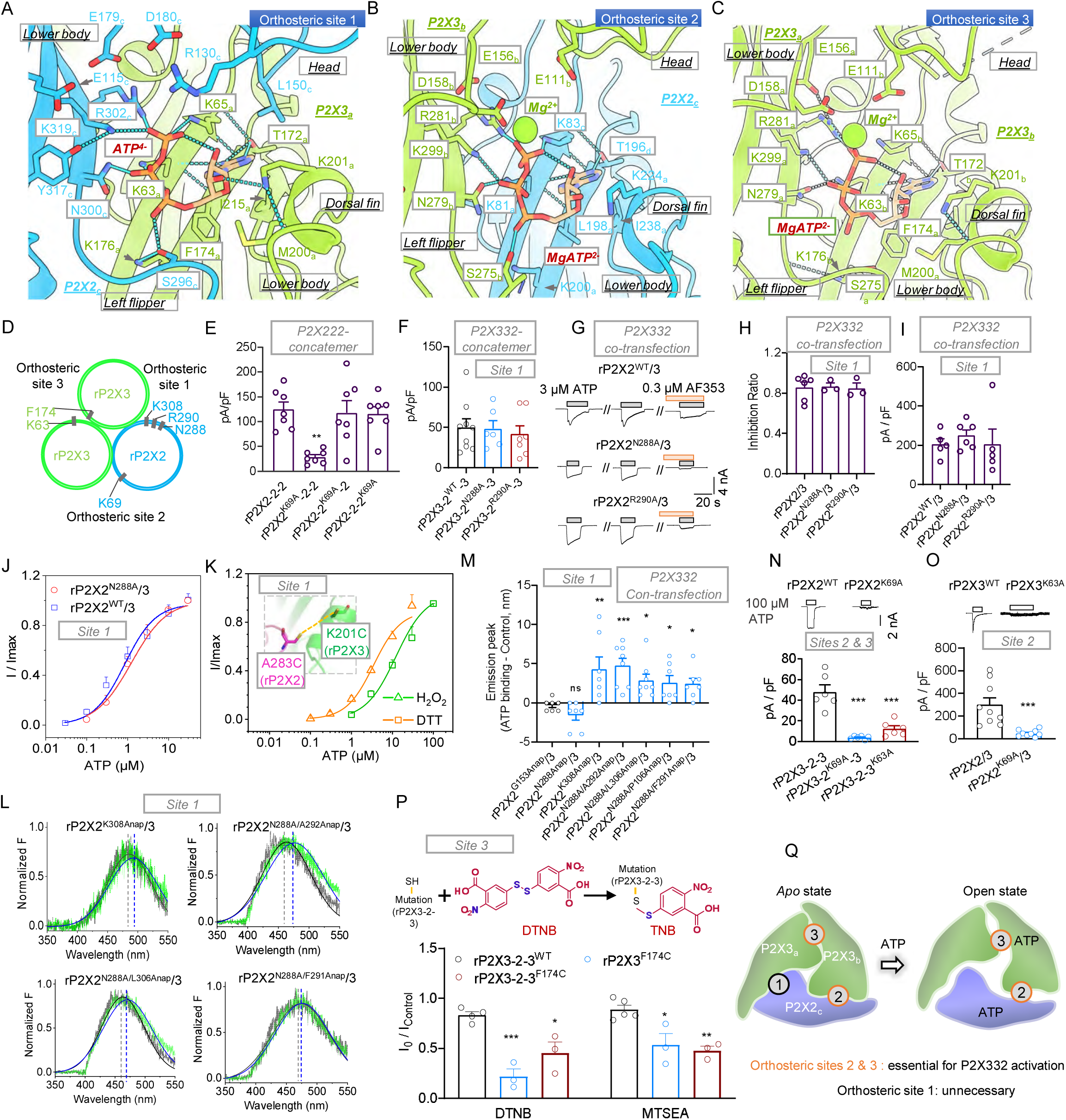
Distinct contributions of the three orthosteric sites to hP2X332 heterotrimer activation. **(A–C)** Close-up views of the three orthosteric sites in the hP2X332 heterotrimer. ATP and key residues are highlighted as stick models for clarity. **(D)** Schematic representation of the three orthosteric sites and their critical residues. **(E, F)** Current density measurements of hP2X222 concatemer (E), hP2X332 concatemer (F), and their respective mutants. **(G–I)** Activation and inhibition of the hP2X323 heterotrimer in the co-transfection system. Representative current traces (G), inhibition ratio by AF-353 (H), and ATP-evoked current density (I) are shown. **(J)** Concentration-response curves for ATP-evoked currents in rP2X332 (co-transfection) and mutants. **(K)** Shifts in ATP concentration-response curves following formation or disruption of a disulfide bond between A283C and K201C. H₂O₂ (0.3%) and dithiothreitol (DTT, 10 mM) were used to induce or reduce the crosslink, respectively. **(L)** Representative ANAP fluorescence emission spectra from mutant-expressing cells in the absence (black) or presence (green) of 1 mM ATP. **(M)** Summary of ANAP emission peak shifts with or without ATP; ns, p > 0.05; *p < 0.05; ***p < 0.001, compared to rP2X2^G153Anap^/3 (one-way ANOVA with Dunnett’s test, F(6,44) = 6.292). **(N, O)** Current density of hP2X332 and indicated mutants in concatemer (N) and co-transfection (O) systems; ***p < 0.001 (unpaired two-tailed t-test or one-way ANOVA with Dunnett’s test, F(2,15) = 30.39). **(P)** Modulation of mutated concatemer P2X3-2-3 currents by cysteine modification with DTNB (5,5’-dithiobis-(2-nitrobenzoic acid)) or MTSEA (N-Biotinylaminoethyl methanethiosulfonate); ***p < 0.001 compared to WT (one-way ANOVA with Dunnett’s test, F(2,8) = 40.92 and F(2,7) = 24.64). **(Q)** Schematic model summarizing the differential contributions of the three orthosteric sites to hP2X332 heterotrimer activation. Each data point represents an independent cell. Source data are provided.

To assess the functional relevance of the three MgATP²⁻/ATP⁴⁻ binding sites, we generated three concatenated receptor constructs: rP2X3-3-2, rP2X3-2-3, and rP2X2-3-3 (Fig. S19A). Among these, rP2X3-2-3 exhibited the largest current amplitudes relative to rP2X3-3-2 and rP2X2-3-3 (pA/pF = 222 ± 18, 105 ± 27, 15.8 ± 1.7, 55.6 ± 3.0, and 44.8 ± 6.0 for rP2X2, rP2X3, rP2X2-3-3, rP2X3-3-2, and rP2X3-2-3; p < 0.05, rP2X3-2-3 vs. rP2X3; p < 0.01, rP2X2-3-3 and rP2X3-3-2 vs. rP2X3; one-way ANOVA with Dunnett’s post hoc test, F(4,14) = 45.77; n = 3–4, Fig. S19C–F), showed relatively high ATP affinity (EC_50_ (μM) of rP2X2, rP2X3, rP2X2-3-3, rP2X3-3-2, and rP2X3-2-3: 14.7 ± 3.5, 1.61 ± 0.16, 24.1 ± 2.7, 9.78 ± 0.96, and 3.11 ± 0.22, respectively; n = 3, Fig. S19D, G), and was most sensitive to the inhibitor AF-353 (inhibition *ratio =* 0.0750 ± 0.0255, 0.970 ± 0.011, 0.886 ± 0.025, 0.234 ± 0.067, 0.866 ± 0.040 for rP2X2, rP2X3, rP2X2-3-3, rP2X3-3-2, rP2X3-2-3; rP2X3-3-2 vs. rP2X3, p < 0.001, one-way ANOVA with Dunnett’s post hoc test, F(4,10) = 117.0; n = 3, Fig. S19E, H). Accordingly, rP2X3-3-2 concatemer was selected as the concatenated construct for subsequent validation experiments.

We also developed a dual-fluorescence reporter system comprising P2X2-eGFP and P2X3-mCherry plasmids. By controlling the *in vitro* transfection ratio and conducting electrophysiological measurements, we optimized a heterologous expression system for rP2X2/3 heterotrimers (Fig. S19I–K). At a 1:5 expression ratio of rP2X2-eGFP to rP2X3-mCherry, rP2X2/3 heterotrimeric currents were detectable and displayed slow desensitization kinetics, distinct from the non-desensitizing rP2X2 and fast-desensitizing rP2X3 homotrimers (Fig. S19I). These heterotrimers were sensitive to AF-353 (inhibition *ratio* = 0.798 ± 0.096, 0.957 ± 0.017, 0.0720 ± 0.0292 for rP2X2/3, rP2X3, rP2X2; p > 0.05, rP2X2/3 vs. rP2X3; p < 0.001, rP2X2 vs. rP2X3; one-way ANOVA with Dunnett’s test, F(2,8) = 29.01; n = 3–5, Fig. S19J–K). The ATP EC_50_ of rP2X2/3 (1.88 ± 0.40 μM) was comparable to that of rP2X3 (1.61 ± 0.16 μM) and distinct from rP2X2 (14.7 ± 3.5 μM, n = 3, Fig. S19K), confirming the suitability of this co-expression system for studying P2X2/3 heterotrimers.

Using the two approaches described above, we introduced targeted mutations into key residues within the three ATP-binding pockets of the P2X332 heterotrimer to assess their contributions to ATP recognition (Fig. 5D). Consistent with prior findings^3, 51^, lysine-to-alanine substitutions in concatenated P2X trimers exhibit positional dependence, with mutations in the A subunit significantly reducing ATP-evoked currents (pA/pF = 124 ± 15, 28.1 ± 4.9, 118 ± 25, and 115 ± 15 for rP2X2-2-2, rP2X2^K69A^-2-2, rP2X2-2^K69A^-2, and rP2X2-2-2^K69A^, respectively; p < 0.01 for rP2X2^K69A^-2-2 versus rP2X2-2-2; p > 0.05 otherwise; one-way ANOVA with Dunnett’s post hoc test, F(3,24) = 7.617; n = 7, Fig. 5E). Accordingly, mutations probing site 1 in the P2X2/3 heterotrimer were introduced exclusively into the P2X2 subunit (Fig. 5D). The mutations rP2X2^N288A^ and rP2X2^R290A^ (corresponding to hP2X2-N300 and R302) were validated, showing significantly altered ATP responsiveness in homotrimers (Fig. S20A) while maintaining surface expression, with a modest reduction relative to WT (Fig. S20B–C). These mutants were used in subsequent concatenation and co-expression experiments. When incorporated into concatenated constructs, mutations that abolish ATP responsiveness in homotrimers did not significantly affect ATP-evoked currents in heterotrimers (pA/pF = 48.5 ± 9.8 and 41.6 ± 10.6; p > 0.05 versus WT rP2X3-3-2: 50.1 ± 10.5; one-way ANOVA with Dunnett’s post hoc test, F(2,19) = 0.1836; n = 6–9, Fig. 5F). These results indicate that site 1 does not contribute to ATP recognition in the P2X332 heterotrimer.

Furthermore, co-expression assays showed that the slowly desensitizing ATP-evoked currents of rP2X2^N288A^/3 and rP2X2^R290A^ remained inhibited by 0.3 μM AF-353 (inhibition ratio = 0.865 ± 0.032 and 0.845 ± 0.053 for rP2X2^N288A^/3 and rP2X2^R290A^/3, respectively; p > 0.05 vs. rP2X2^WT^/3: 0.855 ± 0.052; one-way ANOVA with Dunnett’s multiple comparisons test, F(2,19) = 0.02842; n = 3–6, Fig. 5G–H). Maximal current densities were not significantly different from rP2X2^WT^/3 (pA/pF = 251 ± 28 and 206 ± 77; p > 0.05 vs. rP2X2^WT^/3: 205 ± 29; F(2,13) = 0.3191; n = 5–6, Fig. 5H, right). Apparent ATP affinity was also unchanged (rP2X2^WT^/3 EC_50_ = 0.857± 0.268 μM; rP2X2^N288A^/3 EC_50_ = 1.39 ± 0.13 μM; n = 3, Figs.5J and S20D). These results indicate that ATP recognition at site 1 is not required for activation of the P2X2/3 heterotrimer.

The left flipper and dorsal fin domains, located adjacent to the ATP-binding pocket, undergo relative movements during channel opening and may contribute to conformational coupling. To assess their role at site 1, cysteine substitutions were introduced at rP2X2^283C^ and rP2X3^K210C^ (Fig. 5K). Reduction of the disulfide bond by DTT resulted in a modest increase in apparent ATP affinity (rP2X2^A283C^/P2X3^K210C^ EC_50_ = 12.2 ± 3.7 μM with DTT versus 3.48 ± 0.67 μM with H₂O₂; n = 3, Fig. 5K). Non-reducing SDS–PAGE revealed a trimeric band that disappeared following β-mercaptoethanol (β-Me) treatment, indicating the formation of inter-subunit disulfide bonds (Fig. S21A).

Voltage-clamp fluorometry (VCF) was performed to assess conformational changes of site 1 by incorporating the unnatural fluorescent amino acid (flUAA) Anap into rP2X2 residues constituting site 1 (Fig. S21B–C; see Methods). Using rP2X2^N288A^ as a reference (−1.57 ± 0.64 nm for rP2X2^N288Anap^/3, p > 0.05 versus rP2X2^G153Anap^/3; one-way ANOVA, F(6,44) = 6.292, n = 7, Fig. 5L-M), Anap was inserted at positions A292, L306, P106, and F291. Following heterotrimer activation, these positions exhibited significant rightward shifts in Anap maximal absorption relative to rP2X2^G153Anap^/3 (negative control)^50^ (4.75 ± 0.96 nm for rP2X2^N288A/A292Anap^/3, P < 0.001; 2.88 ± 0.81 nm for rP2X2^N288A/L306Anap^/3, P < 0.05; 2.57 ± 0.90 nm for rP2X2^N288A/P106Anap^/3, P< 0.05; 2.43 ± 0.72 nm for rP2X2^N288A/F291^Anap, P < 0.05; *vs.* −0.286 ± 0.29 nm; one-way ANOVA, F(6,44) = 6.292, n = 7–8, Fig. 5L-M). Anap at site 1 residue rP2X2^K308^ also showed a significant post-activation shift (4.29 ± 1.57 nm, P < 0.01). These results indicate that site I, composed of the P2X2 upper body–left flipper and P2X3 lower body–dorsal fin, is not required for ATP-mediated activation of P2X2/3 heterotrimers, but undergoes conformational changes during channel gating.

To evaluate the contributions of sites 2 and 3 to ATP-dependent activation of hP2X2/3 heterotrimers, we first verified that mutations rP2X2^K69A^ and P2X3^K63A^ abolish ATP recognition in rP2X2 or rP2X3 homotrimers while retaining proper membrane expression (Fig. S20E-H). Co-expression of rP2X2^K69A^ with rP2X3 to disrupt site 2 (Fig. 5D) significantly reduced ATP-evoked currents of rP2X2/3 heterotrimers (pA/pF = 300 ± 63 versus 48.9 ± 9.5 for rP2X2/3 versus rP2X2^K69A^/3, p < 0.01, unpaired t-test, n = 9, Fig. 5O).

In concatenated constructs, mutations rP2X3-2^K69A^-3 and rP2X3-2-3^K63A^, which individually eliminate ATP binding at sites 2 and 3, nearly abolished heterotrimer activation (pA/pF = 3.84 ±0.67 and 12.3 ±2.9 for rP2X3-2^K69A-^3 and rP2X3-2-3^K63A^, respectively; p < 0.001 vs. rP2X3-2-3: 48.2 ± 6.8; one-way ANOVA with Dunnett’s post hoc, F(2,15) = 30.39, n = 6, Fig. 5N). These results demonstrate that ATP binding at sites 2 and 3 is indispensable for activation of hP2X2/3 heterotrimers.

To further interrogate the contribution of site 3 to hP2X2/3 heterotrimer activation, we implemented a covalent occupancy approach using the concatenated mutant rP2X3-2-3^F174C^ (Fig. 5P), thereby selectively abrogating ATP engagement at site 3 (Fig. 5D). DTNB covalent modification at F174C markedly impaired ATP recognition in rP2X3^F174C^ homotrimers and rP2X3-2-3^F174C^ concatenated constructs, while sparing rP2X3-2-3^WT^ (I/I_control_ = 0.219 ± 0.073, 0.454 ± 0.109, and 0.833 ± 0.035, respectively; p > 0.05 for rP2X3^F174C^ vs. rP2X3-2-3^F174C^; p < 0.001 for rP2X3^F174C^ vs. rP2X3-2-3^WT^; p < 0.05 for rP2X3-2-3^F174C^ vs. rP2X3-2-3^WT^; one-way ANOVA with Dunnett’s multiple comparisons test, F (2, 7) = 19.87; Figs. 5P and S20I). MTSEA covalent modification yielded comparable results (I/I_control_ = 0.533 ± 0.116, 0.479 ± 0.045, and 0.887 ± 0.044 for rP2X3^F174C^, rP2X3-2-3^F174C^, and rP2X3-2-3^WT^, respectively; p > 0.05 for rP2X3^F174C^ vs. rP2X3-2-3^F174C^; p < 0.05 for rP2X3^F174C^ vs. rP2X3-2-3^WT^, and p < 0.01 for rP2X3-2-3^F174C^ vs. rP2X3-2-3^WT^; one-way ANOVA with Dunnett’s multiple comparisons test, F (2, 8) = 12.13, n = 3–5; Figs. 5P and S20I). These data collectively demonstrate that ATP binding at orthosteric sites 2 and 3 is requisite for hP2X2/3 heterotrimer activation (Fig. 5Q).

### P2X3–P2X2 interface amino acid differences in P2X332 heterotrimers slow desensitization and regulate gating efficiency compared with P2X3 homotrimers

Compared with P2X3 homotrimers, P2X2/3 heterotrimers show slower desensitization, with τ _activation_ intermediate between P2X2 and P2X3 (Fig. 6A). The apparent affinity of hP2X332 is slightly lower than that of hP2X3 (hP2X2/3 EC_50_ = 716 ± 120 nM; hP2X3 EC_50_ = 508 ± 83 nM; Fig. 6B), consistent with previous studies^45^. Both receptor types exhibit high Ca²⁺ permeability^52^; however, hP2X3 homotrimers desensitize rapidly following single or repeated stimulation. In contrast, P2X2/3 heterotrimers show markedly slower desensitization in the presence of Ca²⁺, allowing more sustained channel opening and enabling their involvement in diverse physiological and pathological processes^19, 53^. Understanding the structural basis for the reduced desensitization of P2X2/3 heterotrimers relative to P2X333 homotrimers remains an important objective.

**Figure 6.**
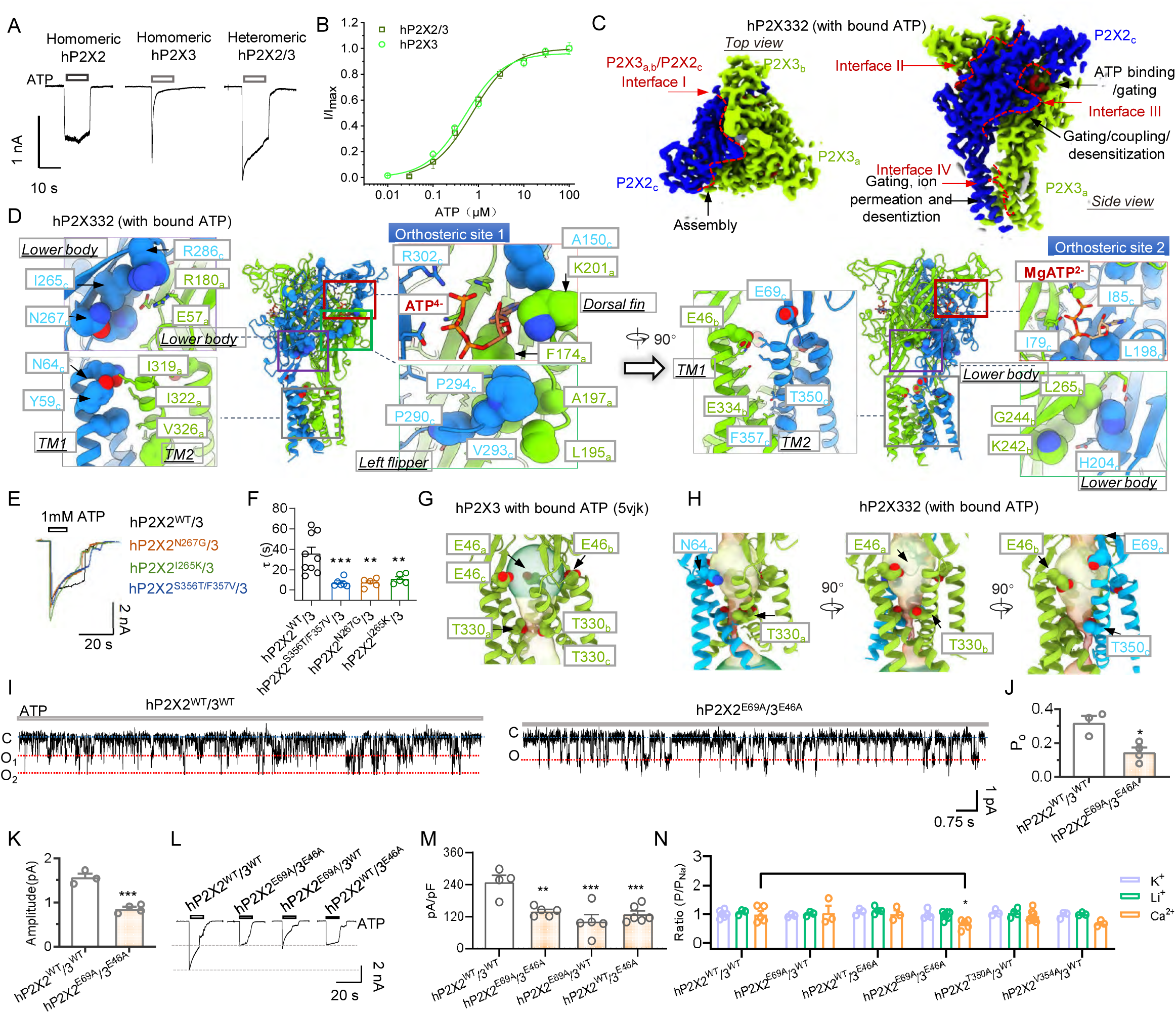
Subunit assembly interfaces of the hP2X332 heterotrimer and their roles in desensitization, conformational transmission, and ion permeation. **(A, B)** Representative current traces (A) and ATP concentration-response curves (B) for homomeric hP2X3 and hP2X332 heterotrimers (the identical data obtained from Fig. S11 B, E). **(C)** Structural depiction of subunit assembly interfaces in the hP2X332 heterotrimer. Red dashed lines indicate inter-subunit interfaces. **(D)** Residues located at the assembly interfaces, with key residues involved in ATP binding, gating, allosteric coupling, desensitization, and ion permeation highlighted as spheres. **(E, F)** Representative currents (E) and summary of τ_activation_ (F) from cells expressing hP2X332 and indicated mutants, evoked by 1 mM ATP. Each point represents an independent cell; **p < 0.01, ***p < 0.001 (one-way ANOVA with Dunnett’s multiple comparisons, F(3,21) = 9.371). (**G, H**) Comparison of cytoplasmic fenestrations in hP2X3 homotrimer (G) versus hP2X2/3 heterotrimer (H). Although both fenestrations form pathways between the receptor and lipid membrane, they differ in dimensions and structural configuration, contributing to distinct cytoplasmic ion permeation profiles. **(I–K)** Unitary currents recorded from outside-out patches at –60 mV in response to 0.1 μM ATP (I) and corresponding summaries of open probability (P₀, J) and amplitude (K); red and blue lines indicate full opening (O) and closing (C). *p < 0.05, ***p < 0.001 (unpaired two-tailed t-test). **(L, M)** Representative currents (L) and summary of current density (M) evoked by 100 μM ATP in cells expressing hP2X332 or mutants. **p < 0.01, ***p < 0.001 (one-way ANOVA with Dunnett’s test, F(3,16) = 10.61). **(N)** Ion permeation measurements for hP2X332 and mutants; *p < 0.05, ***p < 0.001 (two-way ANOVA with Dunnett’s test, F(10,68) = 1.811). Each circle represents an independent cell. Source data are provided.

Following P2X2 incorporation, the P2X332 heterotrimer establishes four distinct interfaces between head-to-tail arranged subunits (Fig. 6C, D). Interface I, involving stacking of the P2X3 and P2X2 upper body domains, primarily contributes to trimer assembly (Figs. 6C, D and S15). Interface II, where P2X3 head domains interdigitate with P2X2 upper body domains, supports trimer assembly and partially participates in ATP recognition and ATP-induced allosteric changes. Interface III, formed by the lower body domains of P2X3 and P2X2, is critical for ATP recognition, undergoes outward expansion during gating, and transmits conformational changes to the TM region, facilitating pore dilation. Interface IV, composed of stacked TM domains, contributes to pore formation and activation (Fig. 6D). Thus, desensitization of P2X332 currents following P2X2 incorporation is likely predominantly associated with interfaces III and V.

We assessed this hypothesis by replacing the divergent amino acids of the P2X2 subunit in hP2X332 with the corresponding residues from P2X3. The Interface IV mutant hP2X2^S356T/F357V^/3 exhibited significantly accelerated desensitization compared with hP2X2^WT^/3 (τ = 7.09 ± 1.6 *vs.* hP2X2^WT^/3: 35.6 ± 6.4 s, p < 0.001, one-way ANOVA followed by Dunnett’s multiple comparisons test, F (3, 21) = 9.371, n = 6-9, Fig. 6E-F). Interface III mutants hP2X2^N267G^/3 and hP2X2^I265K^/3 also showed significantly shortened desensitization times (τ = 7.26 ± 1.9 and 11.1 ± 2.2 s for hP2X2^N267G^/3 and hP2X2^I265K^/3 *vs.* hP2X2^WT^/3: 35.6 ± 6.4 s, p < 0.01, one-way ANOVA followed by Dunnett’s multiple comparisons test, F (3, 21) = 9.371, n = 5-9; Fig. 6E-F).

To further investigate the role of interface residues in gating efficiency, we generated chimeric constructs in which large portions of the lower body domains of P2X2 and P2X3 were swapped within P2X332, yielding rP2X3^ChI^ (G248-R275, rP2X2 numbering) and rP2X3^ChII^ (G310-S326, rP2X2 numbering) (Fig. SA-C). Co-expression confirmed functional heterotrimer formation. The rP2X2/3^ChII^ chimera exhibited a significant increase in apparent ATP affinity (I₀/I_max_= 0.316 ± 0.024 *vs.* WT: 0.178 ± 0.005; p < 0.01, unpaired t-test, n = 4–10; Fig. S22D-F), with a 5-fold leftward shift in EC_50_ (WT: 1.75 ± 0.36 µM; ChII: 0.306 ± 0.004 µM; Fig. S22G), suggesting involvement in activation gating. In contrast, rP2X2/3^ChI^, located in extracellular β11–β12, exhibited a 7-fold rightward EC_50_ shift (WT: 1.75 ± 0.36 µM; ChI: 14.9 ± 2.8 µM; Fig. S22G). Since these domains are adjacent to the TM region and distant from the ATP-binding site, the observed EC_50_ changes likely result from modulation of gating efficiency rather than direct effects on ATP recognition, although long-distance allosteric effects cannot be excluded.

### Acidic residues asymmetrically arranged at hP2X332 TM interfaces govern permeation of cations, including Na⁺ and Ca²⁺

In P2X receptors, ions typically access the channel via lateral fenestrations before traversing the central pore^28, 38^. In hP2X332, asymmetry in TM residue composition modifies this pathway. In P2X3 homotrimers, the symmetrically positioned acidic residue E46 mediates lateral permeation^26^ (Fig. 6G). In P2X2, N64 occupies the equivalent position, with an additional acidic residue, E69, disrupting symmetry in residue and electrostatic distribution compared with P2X3^54^ (Fig. 6H, right). Functionally, the double mutant hP2X2^E69A^/3^E46A^ showed approximately 50% reductions in single-channel conductance and open probability relative to WT hP2X332 (P₀ = 0.148 ± 0.028 *vs.* 0.320 ± 0.040, p < 0.05; amplitude = 0.855 ± 0.046 *vs.* 1.56 ± 0.09, p < 0.001; n = 3–4; Fig. 6I–K). Whole-cell current measurements confirmed significant reductions in both single and double mutants (pA/pF = 102 ± 26, 139 ± 7, and 129 ± 13 *vs*. 249 ± 26 for WT; p < 0.01 for single mutants; p < 0.001 for double mutant; one-way ANOVA with Dunnett’s test, F(3,16) = 10.61, n = 4–6; Fig. 6L, M).

While single mutants hP2X2^E69A^/P2X3^WT^ and hP2X2^WT^/P2X3^E46A^ did not significantly affect K⁺, Li⁺, or Ca²⁺ permeability (p > 0.05, one-way ANOVA followed by Dunnett’s multiple comparisons test, F (10, 55) = 0.9707, n = 3-7, Figs. 6N, S21D), the double mutant hP2X2^E69A^/P2X3^E46A^ showed a reduction in Ca²⁺ permeation (P_Ca²⁺/P_Na⁺ = 0.656 ± 0.072 versus 0.997 ± 0.123 in WT, p < 0.05, one-way ANOVA followed by Dunnett’s multiple comparisons test, F (10, 55) = 0.9707, n = 5, Figs. 6N and S21D). Other mutations had negligible effects relative to WT-hP2X332 (Figs. 6N and S21D). These results highlight the critical role of lateral portal residues P2X2-E69 and P2X3-E46 in directing cation entry and facilitating permeation along the pore. P2X2-specific residues N64 and E69 further modulate Ca²⁺ permeation, with the close positioning of E69 and E46 (Fig. 6H, right) forming a double carboxylate that enhances Ca²⁺ accumulation.

### Intrinsic heterogeneity of the hP2X332 heterotrimer enforces a fully asymmetric mode of action of the allosteric inhibitor AF-219 and ATP

Although two of the three ATP-binding pockets are essential for ligand recognition, it remains unclear whether the approved RCC drug Gefapixant (AF-219; Fig. 7A) follows a distinct recognition mechanism, and whether comparisons among binding pockets can reveal differences between P2X3 homotrimers and P2X2/3 heterotrimers^16, 35, 55^. To explore this, we determined the cryo-EM structure of the hP2X332/AF-219/ATP complex in the presence of both ligands (Figs. S23 and 7A, B). Surprisingly, despite the theoretical capacity of ATP and AF-219 to occupy three orthosteric and allosteric sites, respectively, only a single ATP⁴⁻ molecule was observed at orthosteric site 1, while AF-219 bound exclusively at the P2X3–P2X3 allosteric interface (Fig.7C). At this site, AF-219 interacts with residues S267, S272, S275, and V274 from the left flipper of one P2X3 subunit, and S178, L191, V61, and K176 from the adjacent subunit, through hydrogen bonding and hydrophobic contacts (Fig. 7C, lower)—consistent with previous findings^35^. The densities for both ligands are well defined (Fig. 7C, upper), whereas the remaining pockets are unoccupied (Fig. S24A–C). ∼1.7 μs CMD simulations further confirmed the stability of both ATP and AF-219 within the complex (Figs. S25 and S26).

**Figure 7.**
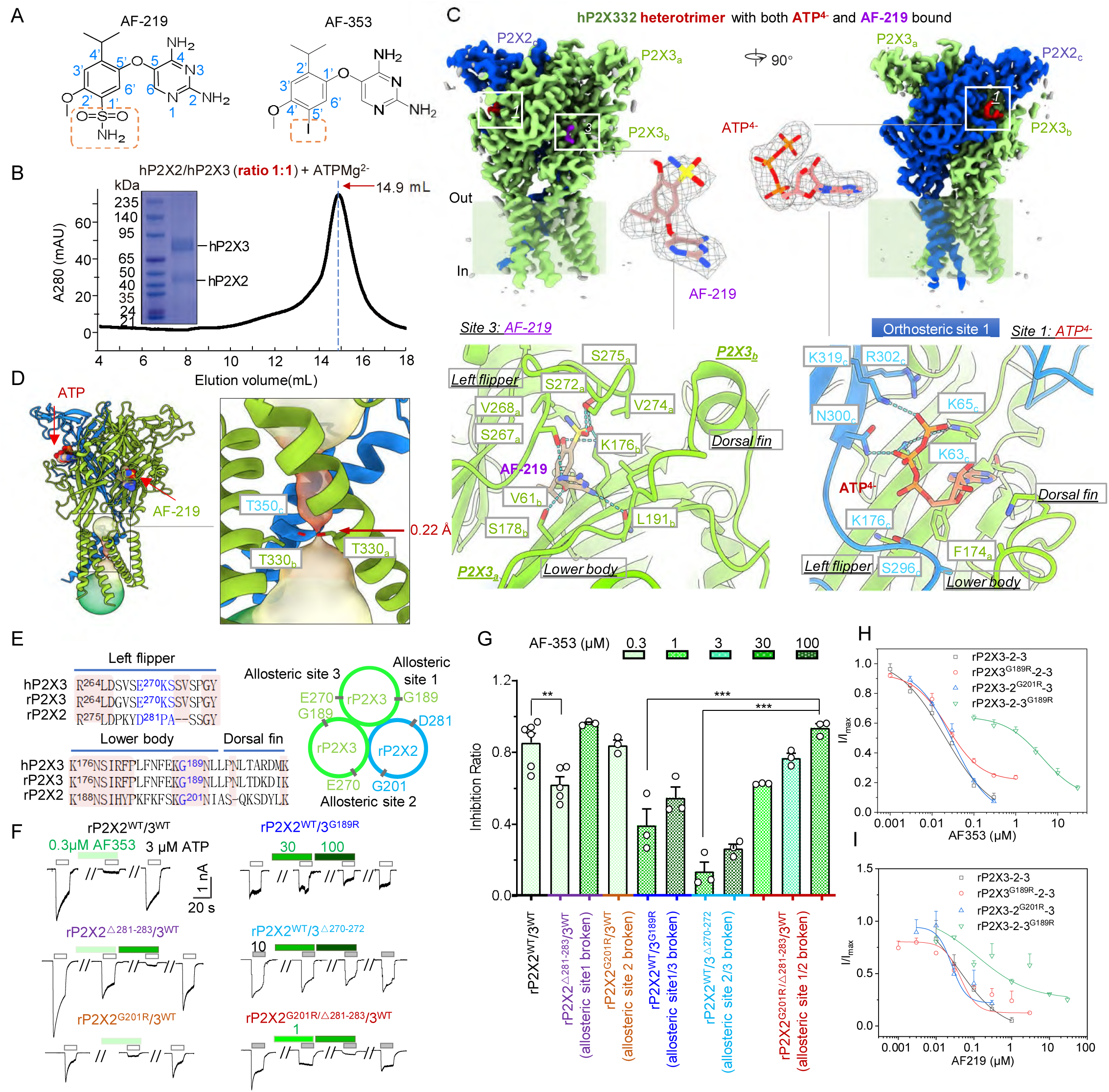
Cryo-EM structures of ATP-bound hP2X332 heterotrimers in complex with AF-219 and their unique ligand-binding stoichiometry. **(A)** Chemical structures of the allosteric modulators AF-219 and AF-353. **(B)** SEC profile and SDS-PAGE analysis of purified P2X2/P2X3 heteromeric complexes co-expressed at a 1:1 ratio in the presence of ATP and Mg²⁺. **(C)** Cryo-EM density map of the hP2X332 heterotrimer bound to both ATP and AF-219, viewed parallel to the membrane plane (upper). Close-up views of orthosteric (lower right) and allosteric (lower left) binding sites show the distinct binding conformations of ATP and AF-219. **(D)** Overall structure of the hP2X332 heterotrimer with ATP and AF-219 bound. Key residues forming the gating element are highlighted as stick models. **(E)** Sequence alignments and schematic representation of the three allosteric binding sites and their key residues in the rP2X332 heterotrimer. **(F, G)** Representative current traces (F) and summary data (G) illustrating the inhibitory effects of AF-353 on rP2X332 activation and indicated mutants. Each data point represents an independent cell. **(H, I)** Concentration–response curves of AF-353 (H) and AF-219 (I) on rP2X3-2-3 concatemers and mutants. Currents were evoked by 30 µM ATP and normalized to those in the absence of modulators. Solid lines indicate fits to the Hill equation. Source data are provided.

These results suggest that heterogeneity in hP2X332 establishes a unique allosteric coupling between the three left flipper–lower body domain pockets and the ATP-binding sites. Structural analysis shows that the hP2X332/AF-219/ATP complex adopts a conformation similar to the apo state (Fig. S24), with overlapping TM domains (Fig. S24D) and a fully closed pore defined by T350 (P2X2) and T330 (P2X3 ×2) (radius: 0.31 Å, Figs. 7D and S24E). The extracellular constriction sites remain unchanged (Fig. S24E). AF-219 binds at the P2X3–P2X3 interface, distorting the left flipper and compressing orthosteric site 3 (Fig. S24A), thereby disrupting ATP recognition. In contrast, left flipper sequence divergence prevents^35^ AF-219 binding at the P2X2–P2X3 interface (Fig. S24B), while structural rearrangements at the P2X3–P2X2 interface compress the allosteric pocket and impair ATP binding at site 2 (Fig. S24C). These coordinated conformational changes provide a compelling explanation for the observation of only a single AF-219 and one ATP molecule in the complex.

Several factors may underlie the observed conformation of the hP2X332/AF-219/ATP complex, including the use of truncated constructs in structural biology, and the conditions employed during protein purification and structural determination. Accordingly, it is imperative to assess whether RCC inhibitors are genuinely recognized by hP2X332^17, 55^ and whether their dysgeusia-associated side effects arise from the mechanism suggested by this structure—specifically, that binding of a single inhibitor molecule at the P2X3–P2X3 interface is sufficient. To address this, we employed a strategy consistent with the stoichiometric approach used for ATP-binding identification (Fig.5 D-P), introducing mutations in co-expression systems analogous to those applied in P2X3–2–3 concatenated constructs, with two key considerations: (1) AF-219 exhibits limited aqueous solubility and submicromolar affinity, which can constrain achievable concentrations in concentration–response assays; consequently, we employed the structurally related analogue AF-353 (Fig. 7A) for cross-validation. (2) Prior studies indicate that residue G189 in the hP2X3 lower body domain is critical for AF-219/AF353 access to its binding pocket, with substitution by a bulky arginine abolishing inhibitor activity, while deletion of residues 270–272 in the left flipper loop markedly reduces inhibitory potency^35^.

Mirroring ATP recognition, we numbered the three distinct AF-219-binding pockets within the hP2X332 heterotrimer: site 1, formed between the P2X2 left flipper and P2X3 lower body; site 2, at the P2X2 lower body–P2X3 left flipper interface; and site 3, between the P2X3 left flipper and lower body (Figs. 7E and S24A–C). To probe the functional significance of each site, we introduced mutations in residues comprising the left flipper and lower body of each allosteric site (Fig. 7E). Co-expression analyses reveal that disruption of site 1 (rP2X2^△281–283^/3^WT^) slightly reduced inhibition by 0.3 μM AF-353 to 62.2 ± 0.4%, compared with 85.5 ± 5.2% in WT (p < 0.01, one-way ANOVA, Tukey’s multiple comparisons test, F (10, 27) = 29.71, n = 3-6, Fig. 7F–G). However, increasing the AF-353 concentration restores inhibitory efficacy to 96.2 ± 0.7%. Disruption of site 2 (rP2X2^G201R^/3^WT^) did not significantly alter inhibition (84.2 ± 2.7% vs. WT, p < 0.05, Fig. 7F, G). Simultaneous disruption of sites 1 and 2 (rP2X2^△281–283/G201R^/3^WT^, 93.7 ± 2.3%) maintained 30 μM AF-353 efficacy. By contrast, mutational disruption of pocket 3 (rP2X2 ^WT^/3^△281–283^ or rP2X2 ^WT^/3^△270–272^, 39.4 ± 9.1% or 13.8 ± 5.3%) abrogated 30 μM AF-353 activity (p < 0.001, one-way ANOVA, Tukey’s multiple comparisons test, F (10, 27) = 29.71, Fig. 7F, G). These results provide compelling evidence for the pivotal role of site 3, in agreement with structural characterization (Fig. 7C).

Concatemer analyses revealed that mutation of G189 in pocket 3 induced a pronounced rightward shift in the concentration–response curves for both AF-353 and AF-219, with maximal inhibition unattainable even at saturating concentrations (IC_50_-AF-353 (nM) = 20.7 ± 7.8, 23.1 ± 2.8, 31.7 ± 6.7, and 4237 ± 971 for rP2X3-2-3, rP2X3^G189R^-2-3, rP2X3-2^G201R^-3, and rP2X3-2-3^G189R^; IC_50_-AF-219 (nM) = 55.6 ± 23.1, 55.2 ± 18.6, 25.9 ± 8.7, and 2183 ± 23 for rP2X3-2-3, rP2X3^G189R^-2-3, rP2X3-2^G201R^-3, and rP2X3-2-3^G189R^, respectively; Fig. 7H-I). These data corroborate the essential role of site 3 and demonstrate that binding of a single AF-219 or AF-353 molecule at the P2X3 left flipper–lower body interface is sufficient to inhibit the P2X2/3 heterotrimeric channel.

## Discussion

P2X receptors can form homotrimeric or heterotrimeric assemblies, yet the physiological and pharmacological diversity observed in native tissues cannot be fully explained by homotrimers alone, indicating the importance of heterotrimers. Among these, the P2X2/3 heterotrimer has been most extensively studied, with substantial biochemical and functional evidence; however, direct structural information has been lacking^46–47, 56–60^. This gap limits understanding of its assembly, in vivo composition, and function, and constrains the rational design and clinical development of selective P2X-targeted drugs. Here, we report the structures of the human P2X2/3 heterotrimer in ligand-free resting, ATP-bound open, and antagonist-bound states, defining its overall architecture and the gating cycle (Fig. 8). ATP recognition exhibits asymmetry: of three potential orthosteric sites, only two bind MgATP^2-^ directly (Fig.5D-P). Additionally, we elucidate the mechanism of the antitussive Gefapixant (AF-219), which achieves inhibition by occupying only an allosteric site between two adjacent P2X3 subunits. This study provides the first high-resolution structural and mechanistic insight into P2X2/3 heterotrimers and establishes a molecular framework for heterotrimer formation, supporting future functional studies and therapeutic development (Fig. 8).

**Figure 8.**
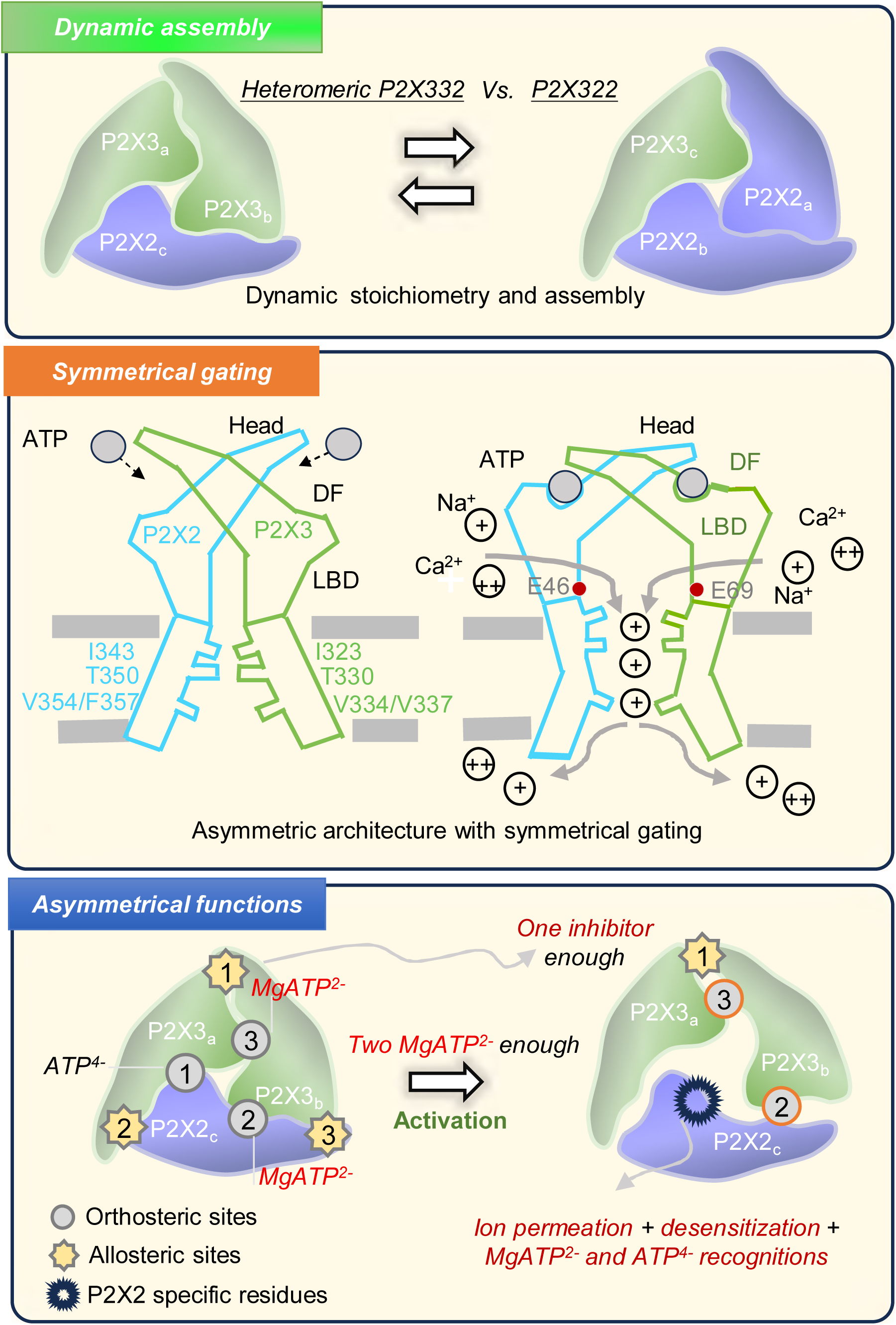
Dynamic assembly, symmetric gating, and asymmetric functions of the heteromeric P2X3/2 receptors. High-resolution cryo-EM structures of human P2X2/3 heterotrimers reveal two assembly configurations, P2X332 and P2X322. P2X332 shows asymmetry across three ATP- binding sites, differentially accommodating MgATP²⁻ and ATP⁴⁻; activation requires only two MgATP²⁻ molecules. Binding of a single allosteric antagonist, Gefapixant, selectively blocks MgATP²⁻ without affecting ATP⁴⁻, evidencing orthosteric–allosteric coupling. Transmembrane domain features establish a distinct Ca²⁺ permeation pathway. Functional assays show largely symmetric gating with modest desensitization differences. Our findings define a structural and mechanistic framework for heterotrimeric P2X receptor function and selective pharmacology.

P2X2/3 heterotrimers assemble into two configurations, P2X332 and P2X322, depending on the expression system and culture conditions. The P2X332 configuration predominates under physiological conditions, consistent with P2X3 expression levels that are 20–30 times higher than P2X2; nevertheless, even low P2X2 expression is sufficient to support P2X322 assembly. Environmental factors, including membrane composition, temperature, and the physicochemical properties of intra- and extracellular solutions, likely contribute to assembly regulation^2, 61–62^. Additional microenvironmental influences—such as post-translational modifications, chaperone-mediated folding, pH fluctuations, and osmotic changes—may further drive dynamic remodeling^2, 63–65^. Differences in P2X2 expression particularly affect heterotrimer stoichiometry and abundance^2, 59, 66^. Overall, microenvironment-dependent assembly may underlie the functional diversity of P2X2/3 heterotrimers and enable adaptive or pathological responses, a hypothesis that warrants further investigation.

Activation of homotrimeric P2X channels relies on the coordinated interplay between the left flipper and dorsal fin domains of adjacent subunits, representing a key allosteric event in channel opening^26, 28^. In the P2X332 heterotrimer, intrinsic subunit asymmetry gives rise to three ATP-binding pockets with distinct recognition modes: one at the P2X3–P2X3 interface and two at the heteromeric P2X2–P2X3 and P2X3–P2X2 interfaces^2^. Structural differences between these heteromeric interfaces lead to divergent pocket architectures and interaction networks. Our results identify two functional ATP-binding sites—the P2X3–P2X3 interface and the pocket formed by P2X3 head–upper body–left flipper with P2X2 lower body–dorsal fin—whereas the third pocket serves a non-canonical role in propagating conformational changes rather than directly binding ATP^2^. Although the physiological implications of this arrangement remain unclear, it may confer sensitivity to varying ATP concentrations across physiological and pathological states^23, 59^. Under normal conditions, the three pockets likely function cooperatively at low ATP levels, whereas under pathological conditions, elevated ATP may enable simultaneous engagement of all sites^23–24, 67^. We further propose that the functional landscape of P2X2/3 receptors is dynamically tuned by the relative expression of P2X2 and P2X3 subunits. Increased P2X3 expression during inflammation may favor homotrimer formation and heightened sensitivity, whereas increased P2X2 expression under homeostasis may stabilize heterotrimeric signaling. Such adaptability is unattainable by homotrimers alone. Thus, the functional interplay between homo- and heterotrimers is indispensable, and our study provides new insight into the molecular basis of their physiological and pharmacological diversity.

Unlike channel activation, desensitization of the P2X332 heterotrimer involves unique conformational rearrangements. We employed chimeric constructs, exchanging key segments and multi-domain regions between P2X2 and P2X3 homotrimers, to assess functional consequences in both homo- and heteromeric assemblies and to delineate structural domains critical for extracellular-to-intracellular signal propagation. At subunit interfaces, the β11, β12, and β14 strands of the left flipper regulate activation gating, while β11 and β12 also contribute to desensitization, with residues N267, I265 and F357 identified as determinants of desensitization. Consideration of transmembrane and intracellular contributions further highlights the P2X2–P2X3 interface as a key locus in gating transitions. Importantly, the transition from activation to desensitization reflects a global, dynamic process, orchestrated through cooperative interactions across multiple interface regions, transmitting signals from the extracellular head to the central core, TM helices, and intracellular tail. This inherent asymmetry is linked to the extensive P2X2 C-terminal domain, which was truncated in prior structural studies (residues S390 onward, 82 amino acids), leaving its precise role in P2X2/3 gating unresolved^34, 38, 53, 68–70^.

P2X3 receptors have emerged as key therapeutic targets for refractory chronic cough^71–73^. Our findings reveal that AF-219 engages the P2X332 heterotrimer through a single binding pocket at the P2X3–P2X3 interface, a site shared with P2X3 homotrimers. This structural conservation suggests that modifying AF-219 to achieve selective discrimination between P2X3 and P2X2/3 receptors is unlikely to succeed, making it difficult to dissociate antitussive efficacy from taste-related side effects. Consequently, this approach appears unsuitable for the development of truly selective P2X3 inhibitors. Although second-generation agents—such as S-600918, BLU-5937, and BAY1817080—have made progress in reducing taste disturbances, they remain constrained by diminished efficacy, restricted patient applicability, and an unclear mechanistic link between cough suppression and taste modulation^74–77^. Notably, these compounds converge on the same extracellular allosteric pocket (PCP)^37, 78–79^, in contrast to first-generation AF-219, which binds beneath the orthosteric ATP site^35^. Our recent work identifies a third extracellular allosteric site (IP–HD) within the head domain of the human P2X3 receptor^50^. Compounds such as quercetin and PSFL2915 that target this site lack selectivity between P2X3 homotrimers and heterotrimers, yet do not produce apparent taste disturbances in animal models, consistent with the absence of such side effects in clinical reports of quercetin. The mechanistic basis for these observations warrants further investigation.

Recent work has uncovered a previously unappreciated allosteric site at the upper TM region of the P2X3 receptor, where antagonists display remarkable affinity and selectivity^36^. Positioned at the subunit interface yet distinct from extracellular regions, this site differs fundamentally from those targeted by existing clinical drugs, opening new avenues for inhibitor design. Building on the structural framework of the P2X332 heterotrimer, future studies should focus on regions that diverge from the P2X3 homotrimer—including the head, left flipper, dorsal fin, and TM domains—with particular attention to the hybrid interface between P2X2 and P2X3 subunits. These regions may harbor exploitable determinants for selective targeting. Guided by the ATP recognition mechanism defined here, next-generation inhibitors could be engineered to directly disrupt ATP engagement at the central orthosteric site formed by the P2X3 head–upper body–left flipper and P2X2 lower body–dorsal fin, offering a strategy for selective heterotrimer inhibition. Complementary efforts to identify novel binding pockets and develop alternative allosteric approaches may ultimately enable precise discrimination between P2X2/3 heterotrimers and P2X3 homotrimers. Such strategies hold promises not only for addressing current limitations in refractory or unexplained chronic cough therapies but also for expanding into broader indications, including pulmonary fibrosis, hypertension, pain, and urological disorders.

Finally, The P2X332 heterotrimer integrates functional specialization at the subunit level: two P2X3 subunits ensure robust Na⁺ and K⁺ permeability, enabling rapid signal initiation during acute injury, while incorporation of a single P2X2 subunit preserves this rapid ion flux and, through coordinated electrostatic interactions within the TM region, enhances Ca²⁺ permeability. This may support localized Ca²⁺ accumulation or facilitate its transit through lateral membrane pathways. In addition, the slower desensitization of the heterotrimer prolongs Ca²⁺ entry. Ion selectivity is further dynamically regulated through modifiable sites on P2X subunits: phosphorylation of P2X2 during inflammation increases Ca²⁺ permeability to potentiate inflammatory and pain signaling, whereas dephosphorylation under basal conditions restores homeostatic balance and prevents tissue damage. This integrated mode of rapid Na⁺/K⁺ conductance and precise Ca²⁺ regulation is a unique property of the heterotrimer, unattainable by homotrimeric receptors.

Collectively, our structural and mechanistic characterization of the P2X2/3 heterotrimer provides new insight into P2X receptor function and offers a foundation for therapeutic innovation, while opening new avenues for atomistic studies of membrane protein biophysics.

## Materials and Methods

### Drugs and mutagenesis

Unless otherwise stated, all compounds were purchased from Sigma-Aldrich (USA). The hP2X3 plasmid was purchased from Open Biosystems. cDNAs for hP2X2 were synthesized by BGl Genomics (China) and subcloned into the pEGFP-N1 vector. The plasmids of rP2X3 and rP2X2 were generously supplied by Drs. Alan North and Lin-Hua Jiang. The mP2X3 construct was generated by site-directed mutagenesis using the rP2X3 plasmid as the template. The cDNAs of hP2X3 and rP2X3 were subcloned into the mCherry-pIRES2 vector. Co-expression systems mimicking the endogenous expression of P2X2/3 were constructed according to a previous report^47^: for hP2X2/3, the transfection ratio of hP2X2 to hP2X3 was set at 1:10, whereas for rP2X2/3, the ratio of rP2X2 to rP2X3 was adjusted to 1:5. Dual fluorescent labeling was employed to confirm the co-expression of P2X2 and P2X3 plasmids in HEK293 cells, with P2X3 tagged by the red fluorescent protein mCherry and P2X2 fused to the green fluorescent protein EGFP. Point mutations were introduced into the target plasmids using the QuikChange Site-Directed Mutagenesis Kit and verified by DNA sequencing. Chimeric constructs were generated using the ClonExpress II One-Step Cloning Kit (Vazyme).

### Cell culture and electrophysiology

Human embryonic kidney 293 (HEK293) and 293T (HEK293T) cells were purchased from the Shanghai Institutes for Biological Sciences. Cells were cultured in Dulbecco’s modified Eagle’s medium (DMEM; Corning, Manassas, USA) supplemented with 1% penicillin-streptomycin (Gibco, Grand Island, USA), 1% glutamate (Gibco) and 10% fetal bovine serum (FBS; PAN-Biotech GmbH, Adenbach, Germany), and incubated at 37°C in a humidified atmosphere of 5% CO₂ and 95% air. P2X2, P2X3, and P2X2/3 plasmids, together with their mutant constructs, were transfected into cells using either calcium phosphate precipitation or Hieff Trans Liposomal transfection reagent.

In this study, perforated, whole-cell, and single-channel patch-clamp configurations were utilized. The perforated patch-clamp approach was adopted for long-duration recordings involving repeated agonist applications, whereas whole-cell recordings were performed to assess peak current responses elicited by a single agonist application. All experiments were performed 24**-**48 hours post-transfection at room temperature (25 ± 2 °C). Patch pipettes were fabricated from borosilicate glass capillaries with a two-stage vertical puller (PC-100, Narishige, Japan), and the pipette resistance ranged from 3 to 5 MΩ; the extracellular solution was composed of 150 mM NaCl, 5 mM KCl, 10 mM glucose, 10 mM HEPES, and 10 mM EDTA, with the pH adjusted to 7.35**-**7.40. For whole-cell patch-clamp recordings, the intracellular solution contained 120 mM KCl, 30 mM NaCl, 0.5 mM CaCl₂, 1 mM MgCl₂, 10 mM HEPES, and 5 mM EGTA, with the pH adjusted to 7.2**-**7.4, while for perforated patch-clamp recordings, the intracellular solution consisted of 75 mM K₂SO₄, 55 mM KCl, 5 mM MgSO₄, and 10 mM HEPES, with the pH adjusted to 7.2. Data acquisition was performed using an Axopatch 200B amplifier (Molecular Devices) in conjunction with a 1550B digitizer (Molecular Devices), the holding potential was clamped at −60 mV before recording, and signals were sampled at 10 kHz and low-pass filtered at 2 kHz. Single-channel recordings were conducted in the outside-out configuration, recording pipettes pulled from borosilicate glass (World Precision Instruments) and fire-polished exhibited a resistance of 8**-**12 MΩ, the holding potential was set to −80 mV, single-channel data were sampled at 50 kHz, filtered at 2 kHz and further low-pass filtered at 200 Hz, and all data were collected and analyzed using Clampfit software (Molecular Devices).

In ionic selectivity experiments, patch pipettes were filled with a solution containing 150 mM NaCl and 10 mM HEPES. The bath solution used to determine the P_Cl_/P_Na_ permeability ratio consisted of 15 mM NaCl, 10 mM glucose, 10 mM HEPES, and 255 mM mannitol. For the determination of other cation permeability ratios relative to Na^+^, all bath solutions contained 10 mM glucose, 10 mM HEPES and 10 mM mannitol as fixed components, with the main cation salt adjusted accordingly: 150 mM KCl for the P_K_/_Na_ permeability ratio, 150 mM LiCl for the P_Li_/P_Na_ permeability ratio, and 100 mM CaCl_2_ for the P_Ca_/P_Na_ permeability ratio. Reversal potentials were determined by applying a voltage ramp and recording the voltage at zero current based on the current–voltage relationship. Permeability ratios were subsequently calculated from the reversal potential shift using the simplified Goldman–Hodgkin–Katz (GHK) equation for permeant ions^80^.

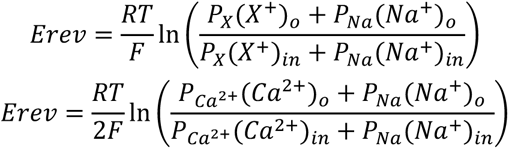

where [X^+^]_o_ and [Na^+^]_i_ are the extracellular and intracellular cation concentrations, respectively. *P_X_/P_Na_* is the permeability ratio of monovalent cations. *P_Ca_^2+^ /P_Na_* is the permeability ratio. *E_rev_* is the change in reversal potential when Na^+^ was replaced by tested cations. *F* is the Faraday constant. *R* is the gas constant. *T* is the absolute temperature.

### Fluorescent unnatural amino acid (flUAA) incorporation and the voltage-clamp fluorometry (VCF) analysis

As previously described^50^, HEK293T cells were cultured in medium supplemented with 20 μM L-Anap (AsisChem Inc.). Plasmids encoding AnapRS, the cognate tRNA, and wild-type (WT) or mutant channel constructs were co-transfected into cells using Lipofectamine® 3000 reagent. The L-Anap-supplemented medium was replaced every 8 h over a 24 h period, and subsequently switched to L-Anap-free medium for an additional 24 h to allow sufficient channel expression prior to experimentation. ANAP fluorescence was excited by a wLS LED light source (PHOTOMETRICS, Tucson, USA) equipped with BP340-390 excitation filters, DM410 dichroic mirrors, and BA420-IF emission filters. Emission spectra of ANAP were collected using an Acton SpectraPro SP-2150 spectrometer (Princeton Instruments) coupled with a Prime 95B CCD camera (Photometrics), and the emission peak was determined by fitting the spectra with a tilted Gaussian distribution^81^.

### Molecular cloning

The trimeric rP2X concatemer was constructed via standard molecular cloning techniques. Three rP2X gene fragments were tandemly cloned into the pcDNA3.1 vector with the incorporation of *SacⅡ* and *HindⅢ* restriction sites introduced (Fig. S19A). A Pro-Arg linker was inserted between the first and the second rP2X subunits, and a Lys-Leu linker was introduced between the second and the third subunits^82^. Additionally, the initiation codons of the second and third subunits were removed^83^, and His-tag, EE-tag, and Flag-tag were fused to the C-terminus of the three subunits in turn, respectively. Given the repetitive sequences present in the trimeric concatemer, the construction of site-directed mutants on this tandem vector was performed by first introducing the target mutations into individual monomeric subunits, followed by replacing the corresponding wild-type subunit in the concatemer with the mutated monomer via restriction digestion and ligation (Fig. S19B).

### Real-time quantitative PCR (qPCR)

qPCR was conducted using the standard curve method for absolute quantification. A 10-fold serial dilution of plasmid containing the *P2RX2* and *P2RX3* gene was used to generate a standard curve for real-time qPCR amplification using TB Green Premix Ex Taq (TaKaRa). The following primers were used to analyze the expression of *P2RX2* and *P2RX3* genes.

*P2rx2* forward: ACAGCCATAAGAAGTTCGA,reverse: CTGATGGAGGGCTGGGTGGCTGAT; *P2rx3* forward: ACAGCATCCGTTTCCCTCTC, reverse: ACCACATCCCCTACCCTCAA.

### Western blotting analysis

HEK293T cells were transfected with WT or mutant plasmids of P2X2, P2X3, and P2X2/3, washed three times with phosphate-buffered saline (PBS, pH 7.4), and then incubated with sulfo-NHS-LC-biotin (Pierce, Germany). The cells were incubated on ice at 4 °C for 30 min, with intermittent shaking every 10 min to ensure uniform labeling of membrane proteins. Glycine was added to terminate the biotinylation reaction. After an additional three washes with PBS, the cells were lysed in 200 μL of RIPA lysis buffer. The lysate was collected after scraping the cells from the culture dish and centrifuged at 12,000 rpm for 30 minutes at 4°C. A 20% (v/v) aliquot of the supernatant was mixed with SDS loading buffer for total protein quantification. The concentration of total protein was determined using the Enhanced BCA Protein Assay Kit (P0010, Beyotime). The remaining supernatant was incubated with NeutrAvidin-conjugated agarose resin (Thermo Scientific, USA) overnight at 4 °C, washed five times with cold PBS, and then eluted with SDS loading buffer to obtain the surface proteins fraction. Then, the protein samples were subjected to SDS-PAGE and transferred onto polyvinylidene difluoride (PVDF) membranes. The membranes were incubated overnight at 4°C with primary antibodies (anti-EE, anti-MYC, anti-eGFP, and anti-Na^+^/K^+^ ATPase, 1:3000; Sigma-Aldrich or anti-GAPDH, 1:3000; Sungene Biotech, China), secondary antibodies anti-rabbit IgG HRP-conjugated (1: 3000, Sungene Biotech, China) and anti-mouse IgG HRP-conjugated (1: 3000, Sungene Biotech, China) at 25°C for 2 h. The protein expression was visualized using electrochemiluminescence (ECL) reagents (HIGH, Thermo Scientific, USA) on the ImageQuant RT-ECL system (Tanon 5200, China).

### Protein expression and purification

Recombinant human P2X2(1-390) and human P2X3(6–364) were co-expressed in HEK293S GNTI⁻ cells (ATCC #CRL-3022) using the BacMam baculovirus delivery system. Recombinant baculovirus was generated in Sf9 insect cells and amplified through three rounds to produce P3 virus. HEK293S GNTI⁻ cells were cultured in Freestyle 293 medium (Gibco #12338018) supplemented with 1% fetal bovine serum (TransSerum #PS301-02) and infected with hP2X2 and hP2X3 viruses at a density of 3.0 × 10⁶ cells/mL. To enhance protein expression, sodium butyrate (Coolaber #CS9931-100g) was added to a final concentration of 10 mM at 12–16 hours post-infection. Cultures were then incubated at 30 °C for 60 hours before harvesting by centrifugation.

To isolate the P2X2/P2X3 heteromeric complex, different virus infection ratios (P2X2:P2X3 = 1:1, 1:5, and 1:10) were tested. At a 1:1 ratio, only the heterotrimer composed of one P2X2 and two P2X3 (denoted P2X322) was obtained. At a 1:10 ratio, only the heterotrimer composed of two P2X2 and one P2X3 (denoted P2X332) was obtained. In contrast, a 1:5 infection ratio yielded a mixture of P2X322 and P2X332 heteromeric complexes, as detailed in the Cryo-EM structure determination section.

Cell pellets from 1 L culture were resuspended in 40 mL of Buffer A (20 mM HEPES pH 7.4, 150 mM NaCl, 0.15 mg/mL DNase I, 100 µM ATP, 100 µM MgCl₂) containing protease inhibitors (1.5 µg/mL leupeptin, 1.5 µg/mL pepstatin A, 1 mM AEBSF, 1 mM benzamidine, 1 mM PMSF). Cells were lysed by sonication and solubilized for 1 hour at 4 °C with continuous stirring in Buffer A supplemented with 0.75% n-dodecyl-β-d-maltopyranoside (DDM, Anatrace), 0.25% (w/v) lauryl maltose neopentyl glycol (LMNG, Anatrace), and 0.025% (w/v) cholesteryl hemisuccinate (CHS, Anatrace). The lysate was centrifuged at 70,000 × g for 30 minutes at 4 °C, and the supernatant was filtered through a 0.45 µm polystyrene membrane (NEST). The solubilized protein was incubated with 2 mL of STarm Streptactin Beads 4FF (Smart-Lifesciences #SA092100) for 30 minutes at 4 °C with gentle agitation. Beads were washed with 50 mL of Buffer B (20 mM HEPES pH 7.4, 150 mM NaCl, 100 µM ATP, 100 µM MgCl₂, 0.05% LMNG, 0.005% CHS), and bound protein was eluted with Buffer B containing 5 mM biotin (Beyotime). The eluate was subsequently applied to Anti-Flag affinity beads (Smart-Lifesciences). After 30 minutes of incubation, the beads were washed with Buffer B, and the P2X2/3 heteromeric complex was eluted using Buffer B supplemented with 0.2 mg/mL Flag peptide (GenScript). Final purification was achieved by size-exclusion chromatography on a Superose 6 Increase 10/300 GL column (Cytiva) pre-equilibrated with Buffer C (20 mM HEPES pH 7.4, 150 mM NaCl, 0.02% glycol-diosgenin (GDN, Anatrace)). For the apo receptor sample, cells were resuspended in hypotonic buffer (20 mM HEPES pH 7.4, 1 mM EDTA, plus protease inhibitors) and incubated for 1 hour at 4 °C. During membrane solubilization, 1 U/mL apyrase (MCE) and 1 mM CaCl₂ were added to hydrolyze residual ATP.

Peak fractions from size-exclusion chromatography were pooled and concentrated to approximately 10 mg/mL using a 50 kDa molecular weight cut-off filter (Millipore). For ligand-bound samples, the protein was incubated with either 200 µM AF-219 (MCE) or 100 µM ATP plus MgCl₂ on ice for 30 minutes prior to cryo-EM grid freezing.

### Cryo-EM *s*ample preparation and data acquisition

Purified protein (4.5 μL) was applied to glow-discharged (10 s) Quantifoil R1.2/1.3 300-mesh copper grids (Electron Microscopy Sciences). Grids were blotted and plunge-frozen in liquid ethane using a Vitrobot Mark IV (Thermo Fisher Scientific) set to 4 °C and 100% humidity, with a blot time of 4–6 s and blot force of 1. Micrographs were acquired on a 300 kV Titan Krios microscope (Thermo Fisher Scientific) equipped with K3 Summit direct electron detector (Gatan) and Bioquantum energy filter (Gatan), operating in counting mode. Detailed data collection parameters for each dataset are provided in Table S1. All datasets were processed according to the workflow described below.

### Cryo-EM structure determination

Cryo-EM data processing workflows are summarized in Fig. S3. All data processing was performed in cryoSPARC v.4^84^ and RELION 4.1^85^. Movie frames were gain-corrected, motion-corrected, and dose-weighted using MotionCor2^86^. Contrast transfer function (CTF) parameters were estimated in cryoSPARC using Patch CTF; micrographs with an estimated resolution better than 4.5 Å were retained. Particles were automatically picked using the Blob picker in cryoSPARC.

For the dataset of the P2X2/3 complex in the presence of ATP and Mg²⁺, a total of 5,029,741 particles were extracted from 7,866 micrographs (3× binned) and subjected to three rounds of reference-free 2D classification. A subset of 200,035 particles displaying well-defined structural features was used for ab initio reconstruction. All extracted particles were then processed through heterogeneous refinement using the ab initio model as an initial reference. After three iterative rounds of heterogeneous refinement, false-positive particles and low-quality reconstructions were effectively removed. From the resulting set, 447,110 particles were re-extracted at full pixel size and refined using non-uniform refinement under C1 symmetry, yielding a 3D reconstruction at 2.9 Å global resolution. The particles were subsequently subjected to Bayesian polishing in RELION 4.1, followed by an additional round of heterogeneous refinement to further improve particle quality. A final set of 349,034 high-quality particles was subjected to 3D classification in RELION 4.1 (initial resolution: 6 Å; symmetry: C1; number of classes: 2; T = 16), yielding two reconstructions exhibiting subtle structural differences. Notably, the region between residues D158 and I164 forms a loop in P2X3 but adopts a β-sheet conformation in P2X2. This distinguishing structural feature enabled us to unambiguously assign the two maps to P2X3-rich (P2X332) and P2X2-rich (P2X322) conformations. Both maps were imported into cryoSPARC as reference volumes for a subsequent heterogeneous refinement (initial resolution: 6 Å; symmetry: C1; number of classes: 2). Particles corresponding to each class were separately subjected to global and local CTF refinement followed by non-uniform refinement in cryoSPARC v.4, resulting in final reconstructions at overall resolutions of 2.68 Å (P2X332) and 2.95 Å (P2X322). The accuracy of structural assignments between P2X2 and P2X3 was further validated by analyzing residues with significant side-chain density that differ between the two subunits (see Fig. S3, S13, and S23).

For the P2X2/3 complex dataset bound to inhibitor AF-219, a total of 7,053,134 particles were extracted from 5,683 micrographs (3× binned) and subjected to three rounds of reference-free 2D classification. From these, a subset of 233,496 particles displaying well-resolved structural features was selected for ab initio reconstruction. All extracted particles then underwent heterogeneous refinement using the ab initio model as a reference. After three iterative rounds of heterogeneous refinement, false-positive particles and low-quality reconstructions were effectively removed. A total of 408,368 particles were subsequently re-extracted at full pixel size and processed by non-uniform refinement under C1 symmetry, yielding a reconstruction at 3.1 Å global resolution. These particles were further improved by Bayesian polishing in RELION 4.1, followed by one additional round of heterogeneous refinement. A final set of 380,230 high-quality particles was subjected to 3D classification in RELION 4.1 (initial resolution: 6 Å; symmetry: C1; number of classes: 2; T = 16), which yielded two reconstructions with no appreciable structural differences; both were assigned to the P2X332 conformational state. These particles were then transferred to cryoSPARC v.4 for global and local CTF refinement, followed by non-uniform refinement, resulting in a final map at an overall resolution of 2.87 Å. The structural assignment was validated by analyzing residues exhibiting distinct side-chain densities between P2X2 and P2X3 subunits (see Fig. S23).

For the P2X2/3 complex in apo state, a total of 3,980,835 particles were extracted from 4,730 micrographs (3× binned) and subjected to three rounds of reference-free 2D classification. From these, a subset of 213,736 particles displaying well-resolved structural features was selected for ab initio reconstruction. All extracted particles then underwent heterogeneous refinement using the ab initio model as a reference. After five iterative rounds of heterogeneous refinement, false-positive particles and low-quality reconstructions were effectively removed. A total of 130,771 particles were subsequently re-extracted at full pixel size and processed by non-uniform refinement under C1 symmetry, yielding a reconstruction at 3.7 Å global resolution. These particles were further improved by Bayesian polishing in RELION 4.1, followed by three additional round of heterogeneous refinement. A final set of 59,230 high-quality particles was subjected to 3D classification in RELION 4.1 (initial resolution: 6 Å; symmetry: C1; number of classes: 2; T = 16), which yielded two reconstructions with no appreciable structural differences; both were assigned to the P2X332 conformational state. These particles were then transferred to cryoSPARC v.4 for global and local CTF refinement, followed by non-uniform refinement, resulting in a final map at an overall resolution of 3.3 Å. The structural assignment was validated by analyzing residues exhibiting distinct side-chain densities between P2X2 and P2X3 subunits (see Fig. S3, S13, and S23).

Atomic models were initially built using an AlphaFold-predicted structure as a reference^87^. Iterative real-space refinement was carried out in Coot^88^, followed by further real-space refinement in PHENIX^89^. The final models exhibit good stereochemistry and show strong correlation with the cryo-EM density, as indicated by Fourier shell correlation (FSC) values (Fig. S3, S13, and S23, and Table S1). Structural figures were prepared using UCSF Chimera^90^, ChimeraX^91^, and PyMOL (www.pymol.org).

### Molecular dynamic (MD) simulations

As previously reported^30, 50^, energy-minimized models of hP2X223-ATP, hP2X332-ATP, *apo* hP2X332, hP2X332-ATP-AF-219, and hP2X332-AF-219 served as starting structures for all MD simulations. A large 1-palmitoyl-2-oleoyl-sn-glycero-3-phosphocholine (POPC) bilayer at 300 K, provided by the System Builder module in Desmond^92^, was constructed to establish an appropriate membrane environment according to the P2X receptor of the OPM database (https://opm.phar.umich.edu)^93^, enabling correct embedding of the transmembrane (TM) region of hP2X2/3. The hP2X223-ATP-POPC, hP2X332-ATP-POPC, hP2X332-ATP-AF-219-POPC, hP2X332-AF-219-POPC, and apo hP2X332-POPC systems were subsequently solvated with simple point charge (SPC) water molecules. Counterions were subsequently added to neutralize the net negative charge of the system. NaCl (150 mM) was supplemented into the simulation box to mimic physiological ionic strength. The default relaxation protocol implemented in Desmond was applied to each system before the production run. In brief: (1) 100 ps NVT (constant number, volume, and temperature) simulation with Brownian dynamics at 10 K, with heavy atoms of the solute restrained; (2) 12 ps NVT simulation using a Berendsen thermostat at 10 K with small time steps, with heavy atoms of the solute restrained; (3) 12 ps NPT (constant number, pressure, and temperature) simulation using a Berendsen thermostat and barostat at 10 K and 1 atm, with heavy atoms of the solute restrained; (4) 12 ps simulation using a Berendsen thermostat and barostat at 300 K and 1 atm, with heavy atoms of the solute restrained; (5) 24 ps simulation using a Berendsen thermostat and barostat at 300 K and 1 atm, without any restraints. Following equilibration, MD simulations were performed for approximately 0.5 μs. Long-range electrostatic interactions were computed using the smooth particle mesh Ewald (SPME) approach. Trajectories were saved at an interval of 200 ps, and other default parameters in Desmond were adopted for conventional MD simulations^94^. All simulations employed the OPLS-2005 all-atom force field^95^, which is compatible with proteins, ions, lipids, and SPC water. The Simulation Interaction Diagram (SID) module in Desmond^50^ was utilized to analyze the binding interactions between SIV/ATP and hP2X3. All MD simulations were conducted on a Dell T7920 workstation equipped with an NVIDIA Tesla K40C GPU, or a Caowei 4028GR server equipped with an NVIDIA Tesla K80 GPU. System preparation, trajectory analysis, and visualization were performed on a Dell T7500 graphics workstation with 12 CPU cores.

### Animals

All experiments were performed on adult mice aged 8–12 weeks. Wild-type (WT) C57BL/6 mice were purchased from Huachuang Sino Company (Taizhou, China). All mice were housed and bred in the specific pathogen-free animal facility of China Pharmaceutical University. The experimental protocols were approved by the Animal Ethics Committee of China Pharmaceutical University (YSL-202507013), and all animal procedures were performed in compliance with the National Institutes of Health Guide for the Care and Use of Laboratory Animals.

### Complete Freund’s adjuvant (CFA) model

Inflammatory pain was induced by complete Freund’s adjuvant (CFA)^43^. Briefly, 1 mg/kg CFA was prepared as a 1:1 oil-saline emulsion and administered via intraplantar injection into the right hind paw.

### Isolation and culture of dorsal root ganglion (DRG) neurons for electrophysiological recording and qPCR analysis

A midline incision was made along the dorsal surface of mice using surgical scissors, and the skin and muscle layers were carefully dissected to expose the vertebral column. Vertebrae were gently removed to access the spinal cord, and L4–L6 dorsal root ganglia were meticulously harvested and collected in ice-cold saline solution. The isolated DRG tissues were minced into small fragments and digested with Type II collagenase (Sigma-Aldrich). Following centrifugation, the supernatant was discarded, and the cell pellets were resuspended in fresh DMEM culture medium, seeded onto poly-D-lysine (PDL)-coated culture dishes, and incubated at 37°C in a 5% CO₂ incubator for subsequent electrophysiological recording and qPCR analysis.

For electrophysiological recordings, the extracellular solution was prepared with 150 mM NaCl, 5 mM KCl, 10 mM glucose, and 10 mM HEPES, with the pH adjusted to 7.35–7.40. The intracellular pipette solution contained 30 mM KCl, 2 mM CaCl₂, 1 mM MgCl₂, 10 mM HEPES, and 110 mM K-gluconate, with the pH adjusted to 7.2–7.4. For qPCR analysis, total RNA was extracted immediately after cell adherence, and reverse transcription was performed using the EZ-press RNA purification kit (B0004-plus, EZBioscience) and PrimeScript RT kit (RR047A, TaKaRa), respectively, in strict accordance with the manufacturer’s protocols.

## Supporting information

Compressed package includes PDB validation report, electron density map and the structural model in PDB format.

## Data analysis

All electrophysiological recordings were analyzed using Clampfit 10.6 (Molecular Devices). All results are expressed as means ± SEM. Statistics analyses were performed using one-way analysis of variance (ANOVA) or unpaired Student’s t-test, with P < 0.05 considered statistically significant unless otherwise stated. The concentration-response curves were fitted using the *Hill* equation: I/I_max_ = 1/(1+( EC_50_ or IC_50_/[ATP or inhibitor])^n^), where I is the normalized current at a given concentration of ATP or inhibitor, I_max_ is the maximum normalized current, EC_50_ or IC_50_ is the concentration of the ATP or inhibitor exhibiting the half-maximum effect and n is the *Hill* coefficient^30^.

## Acknowledgments

This work was supported by the National Key Research and Development Program of China (2025YFA1308903 to Y. Y), National Natural Science Foundation of China (32371289 to Y. Y, 82474171 to C. G, 32401027 to S.M.Y and 32370046 to C.W.), Gansu Science and Technology Leading Talent Project (26RCKA010, to Y. Y), Xing Yao Leading Scholars of China Pharmaceutical University (2021) to Y. Y, CAMS Innovation Fund for Medical Sciences (2021-I2M-3-001 to Y. Y), Medical Innovation and Development Project of Lanzhou University (lzuyxcx-2022-156 to Y. Y), the Postdoctoral Fellowship Program of CPSF (No. GZC20252599 to D. W.), Guangdong Basic and Applied Basic Research Foundation (2024A1515030085 to C.W.), Startup of Shenzhen Institute of Advanced Technology, Chinese Academy of Sciences (to C.W.) and Hunan Provincial Natural Science Foundation (2024JJ6242 to X. Y).

## Author Contributions Statement

Y. Y. and C. W. designed the project; D. W., X. Y., A. Z., X. C. and Y. G. performed cell culture, patch-clamp recording, Western blotting; C. W., W. C., and B. C. performed Cryo-EM experiments; M. S. performed VCF; M. S. and A. Z. performed ion permeation experiments; D. W., A. Z., and Y. G. performed animal experiments; Y. Y., and P. C. did MD simulations; Y. Y., C. W., D. W., M. S., C. G., and C. L. analyzed data; and Y. Y., D. W., C. G., M. S., and M. Z. wrote the manuscript. All authors discussed the results and commented on the manuscript.

---

**Figure S1.**
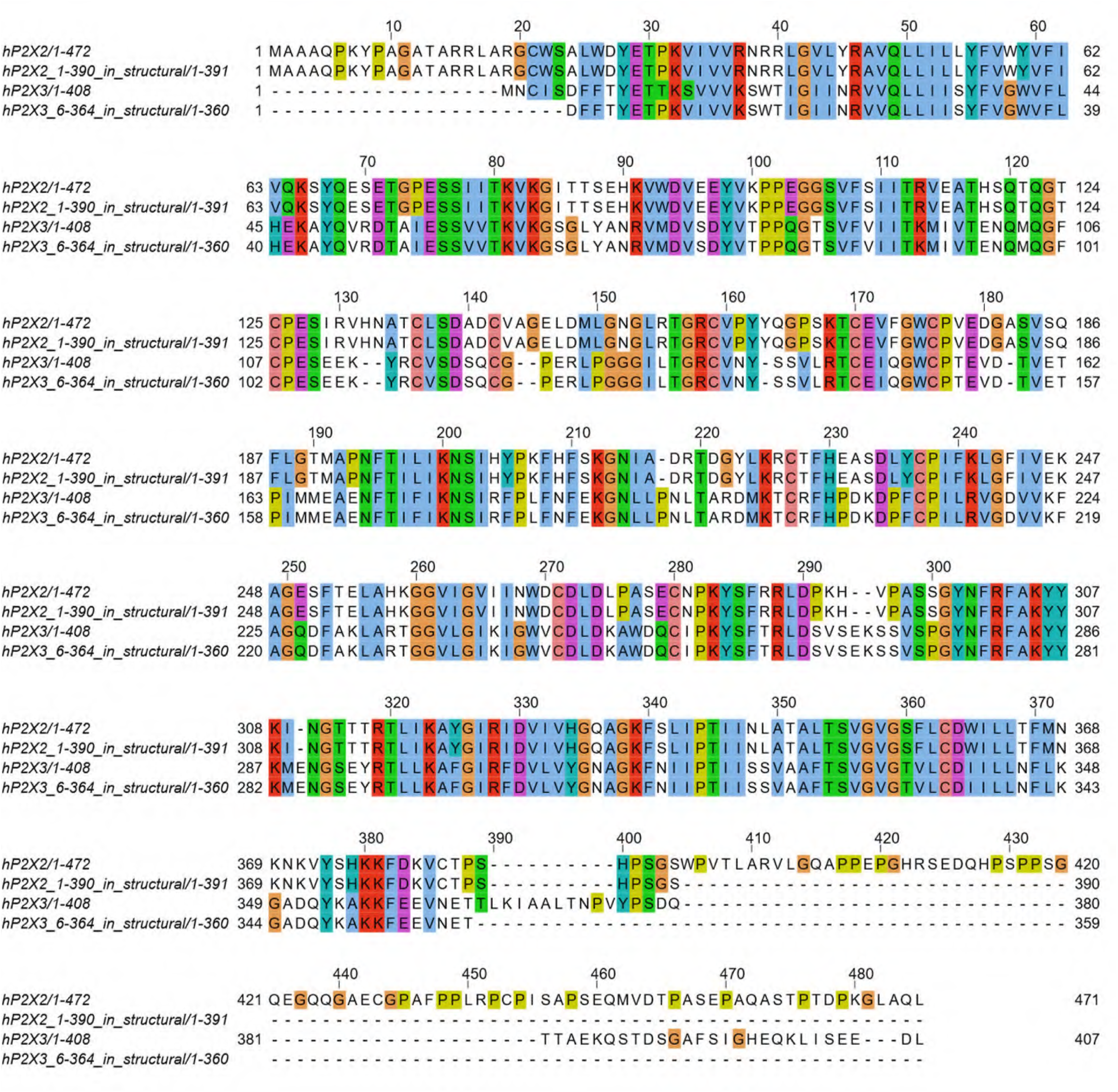
Multiple sequence alignments of human P2X2 (hP2X2), human P2X3 (hP2X3), recombinant hP2X2 (residues 1–390), and recombinant hP2X3 (residues 6–364) constructs employed in structural analyses. UniProt accession numbers for the corresponding P2X subtypes are: hP2X2 (Q9UBL9), rat P2X2 (rP2X2; P49653), mouse P2X2 (mP2X2; Q8K3P1), hP2X3 (P56373), rat P2X3 (rP2X3; P49654), and mouse P2X3 (mP2X3; Q3UR32). Sequence alignments were generated using TCoffeeWS and subsequently analyzed and visualized in Jalview. Residue coloring was applied according to the Clustal scheme within Jalview, reflecting both physicochemical properties of amino acid side chains and the degree of positional conservation across the alignment.

**Figure S2.**
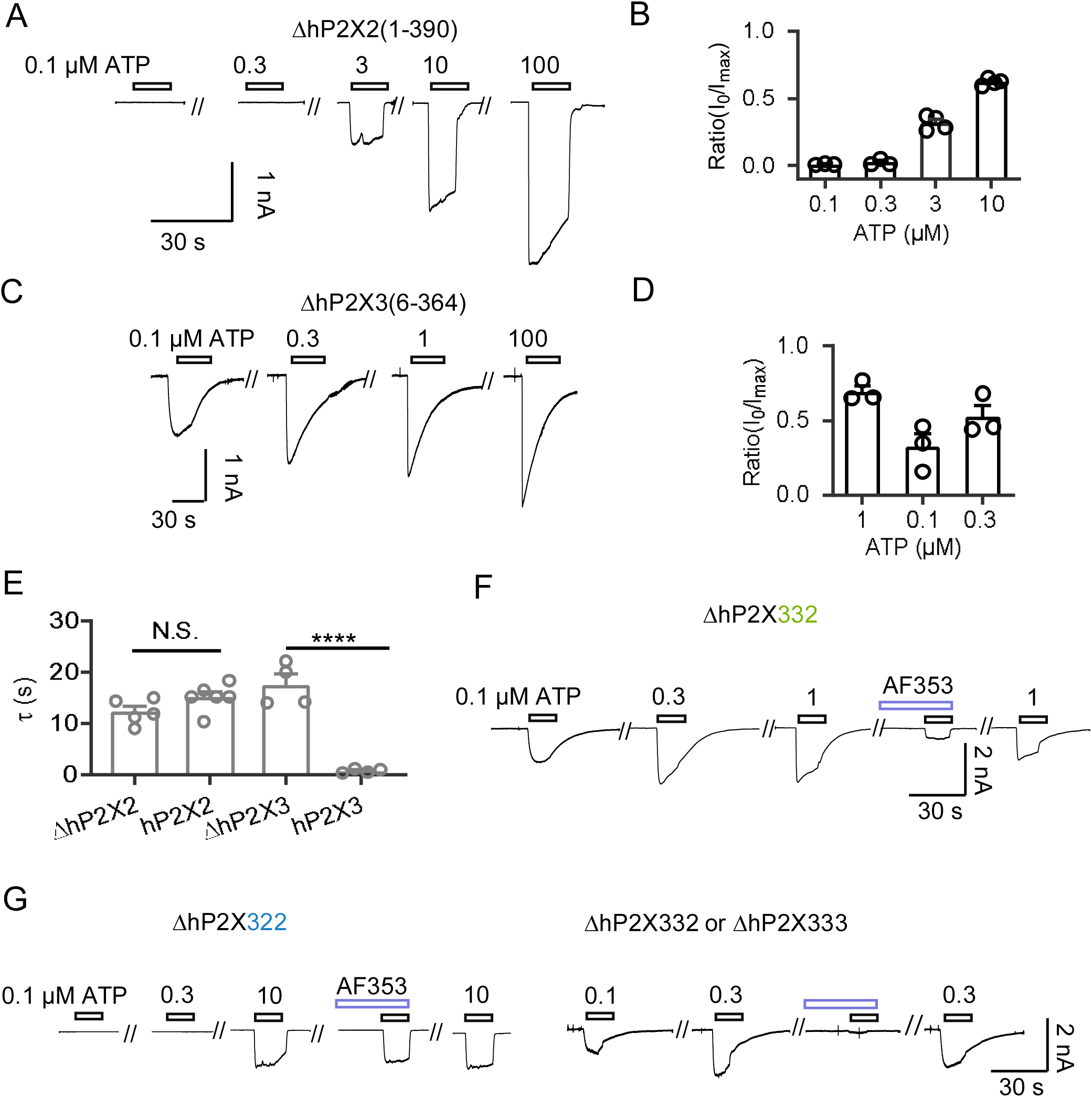
Current recordings of recombinant human P2X2 (ΔhP2X2, residues 1–390) and ΔhP2X3 (residues 6–364) used in structural studies. **(A, B)** Representative current traces (A) and pooled data (B) from cells expressing ΔhP2X2, evoked by 0.1–100 μM ATP. Each circle represents an individual cell. **(C, D)** Representative current traces (C) and pooled data (D) from cells expressing ΔhP2X3, evoked by 0.1–100 μM ATP. Each circle represents an individual cell. **(E)** Summary of activation time constants (τ_activation). N.S., not significant; ***P < 0.001; one-way ANOVA with Tukey’s multiple comparisons test (F(3,15) = 31.48). **(F, G)** Representative current traces from cells co-expressing ΔhP2X2 and ΔhP2X3. Homo- and heterotrimers were distinguished based on ATP-evoked current kinetics and sensitivity to AF-353. AF-353 inhibits P2X3 homotrimers and P2X332 heterotrimers, but not P2X322 heterotrimers. All summary data are presented as mean ±s.e.m. Source data are provided.

**Figure S3.**
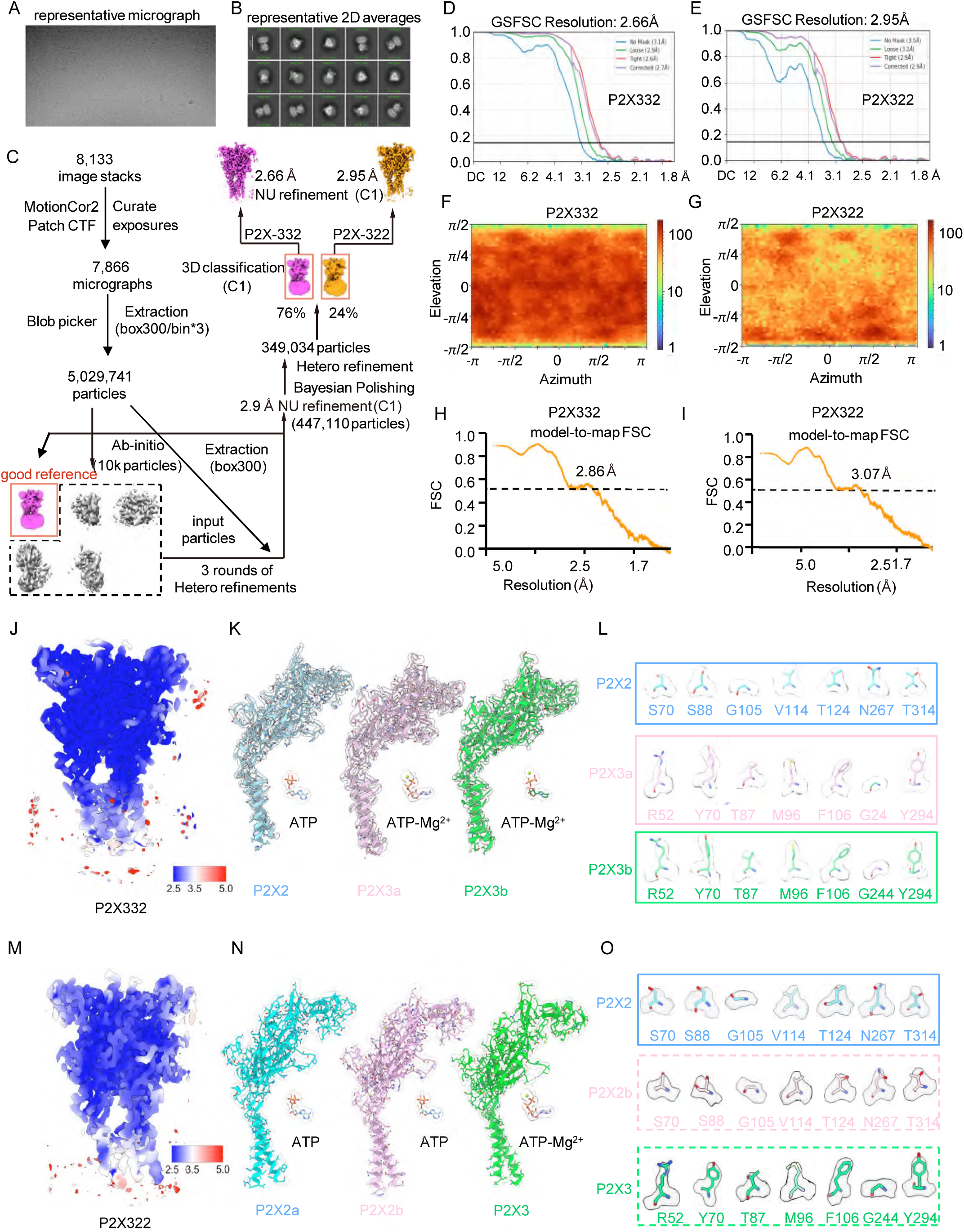
Cryo-EM analysis of the P2X2/P2X3 heteromeric complex co-expressed at a 1:5 ratio in the presence of ATP and Mg²^+^. (**A**) Representative cryo-EM micrograph. (**B**) Selected 2D class averages. (**C**) Data processing workflow for 3D reconstruction. (**D, E**) Fourier shell correlation (FSC) curves from the gold-standard refinement. (**F, G**) Angular distribution of particles used in the final 3D reconstruction. (**H, I**) FSC curves between the final atomic model and the corresponding sharpened map. (**J, M**) Local resolution estimation of the sharpened cryo-EM map. (**K, N**) Representative cryo-EM density for key structural features in each subunit and ATP. (**L, O**) Analysis of residues exhibiting distinct side-chain densities between P2X2 and P2X3 subunits.

**Figure S4.**
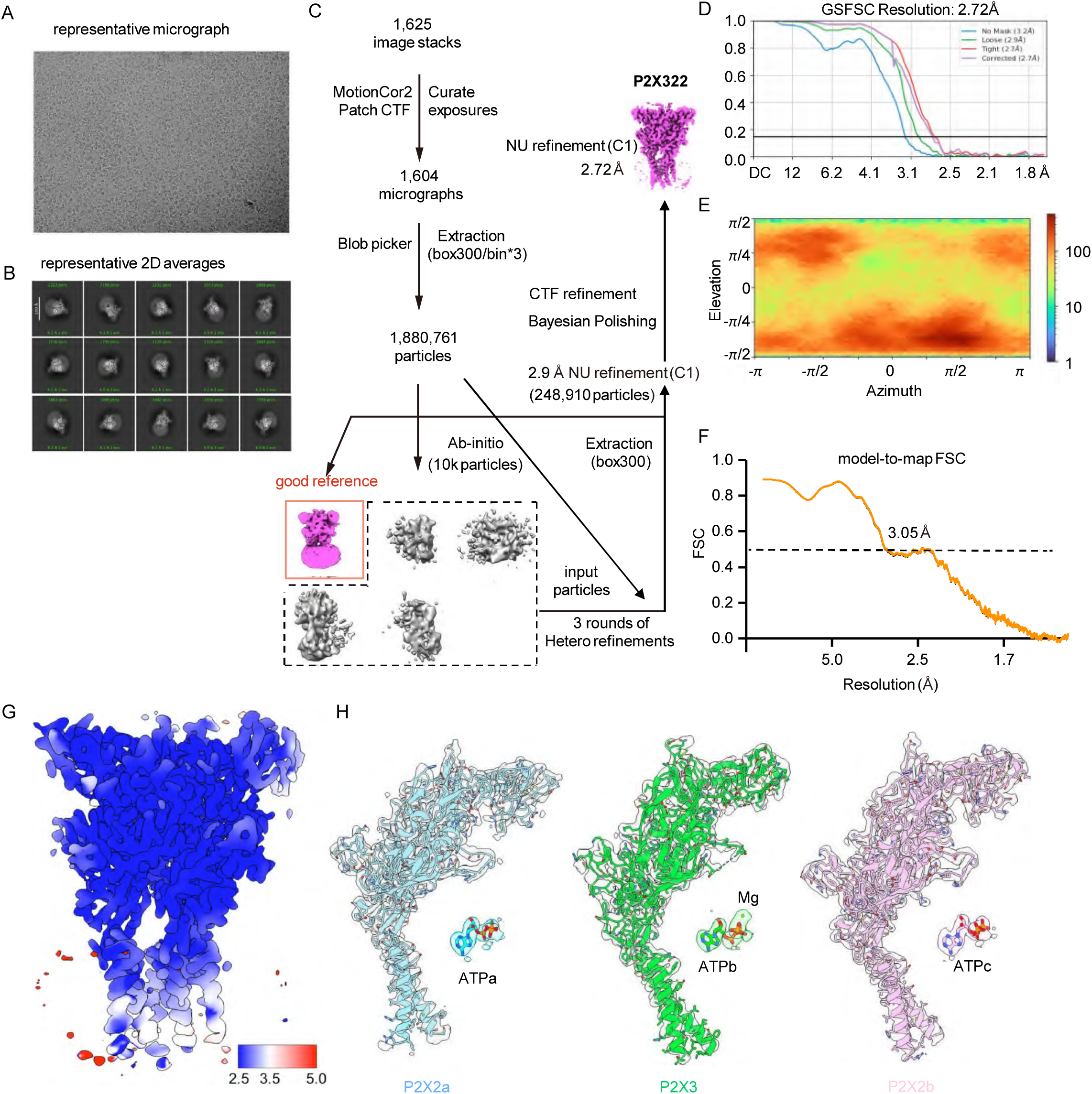
Cryo-EM analysis of the P2X2/P2X3 heteromeric complex co-expressed at a 1:1 ratio in the presence of ATP and Mg²^+^. (**A**) Representative cryo-EM micrograph. (**B**) Selected 2D class averages. (**C**) Data processing workflow for 3D reconstruction. (**D**) Fourier shell correlation (FSC) curves from the gold-standard refinement. (**E**) Angular distribution of particles used in the final 3D reconstruction. (**F**) FSC curves between the final atomic model and the corresponding sharpened map. (**G**) Local resolution estimation of the sharpened cryo-EM map. (**H**) Representative cryo-EM density for key structural features in each subunit and ATP.

**Figure S5.**
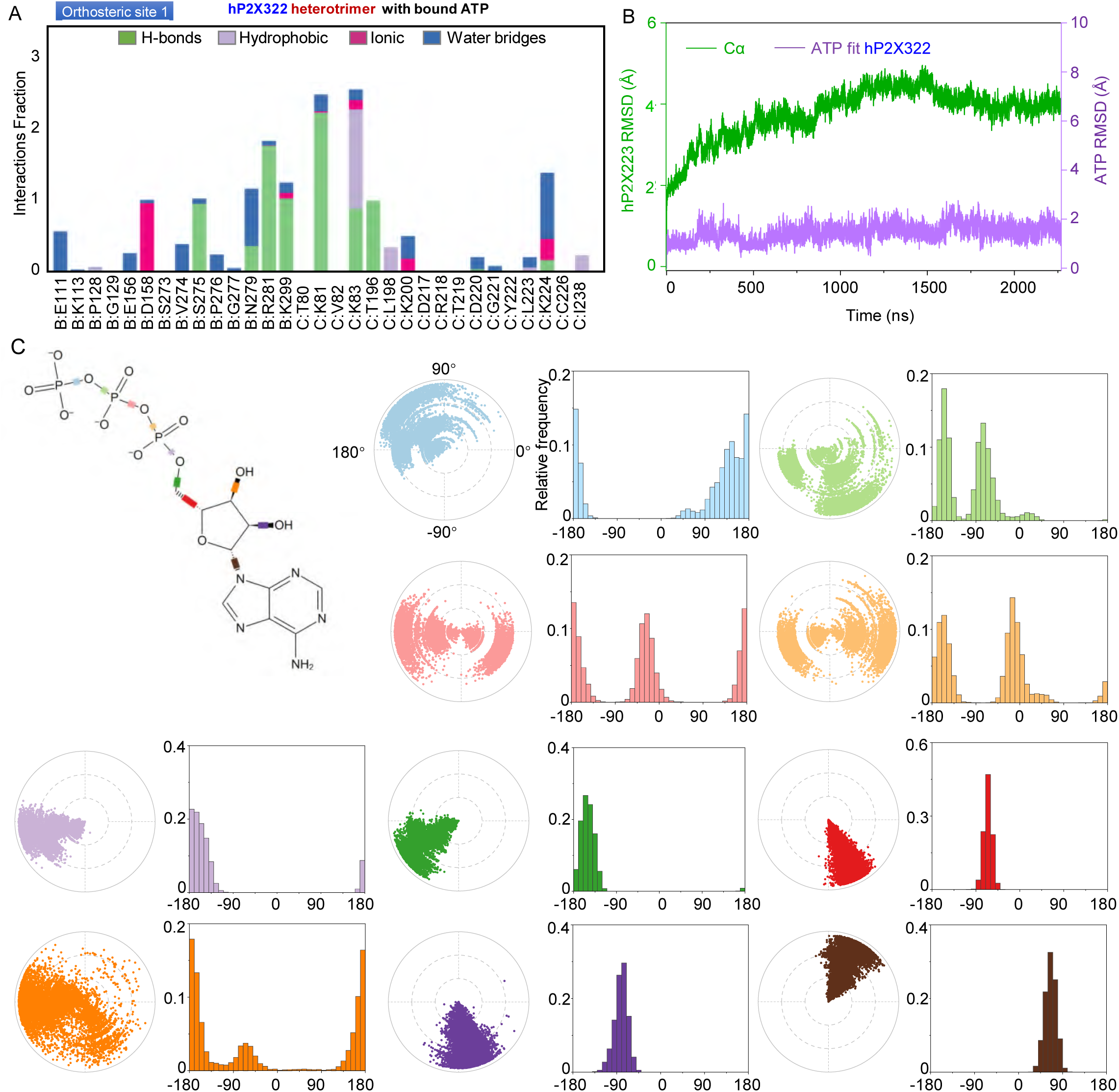
Conventional molecular dynamics (MD) simulations to study the interaction of hP2X322 heterotrimer in complex with ATP at orthosteric site 1. (**A**) The interaction between hP2X322 and ATP was monitored throughout the MD simulations. hP2X322/ATP interactions were classified into four types: hydrogen bonds (green), hydrophobic (light purple), ionic interactions (pink), and water bridges (blue). The stacked histograms are normalized over the course of the trajectory. (**B**) Backbone root-mean-square deviation (RMSD) analysis of the binding process of ATP to hP2X322 throughout the MD simulation. (**C**) Torsion plots of ATP summarizing the conformational evolution of each rotatable bond every 10 ns throughout the simulated trajectory. 2D schematics of ATP are shown as color-coded rotatable bonds. The radial plots represent the conformation of the torsion bodies. The center of the radial plot represents the beginning of the simulation, plotting the temporal evolution in the radial direction outward. The histogram summarizes the data of the corresponding radial plot, which represents the probability density of the torsion. The relationship between histogram and torsional potential gives insight into the conformational strain that the ligand underwent to maintain the hP2X322-bound conformation.

**Figure S6.**
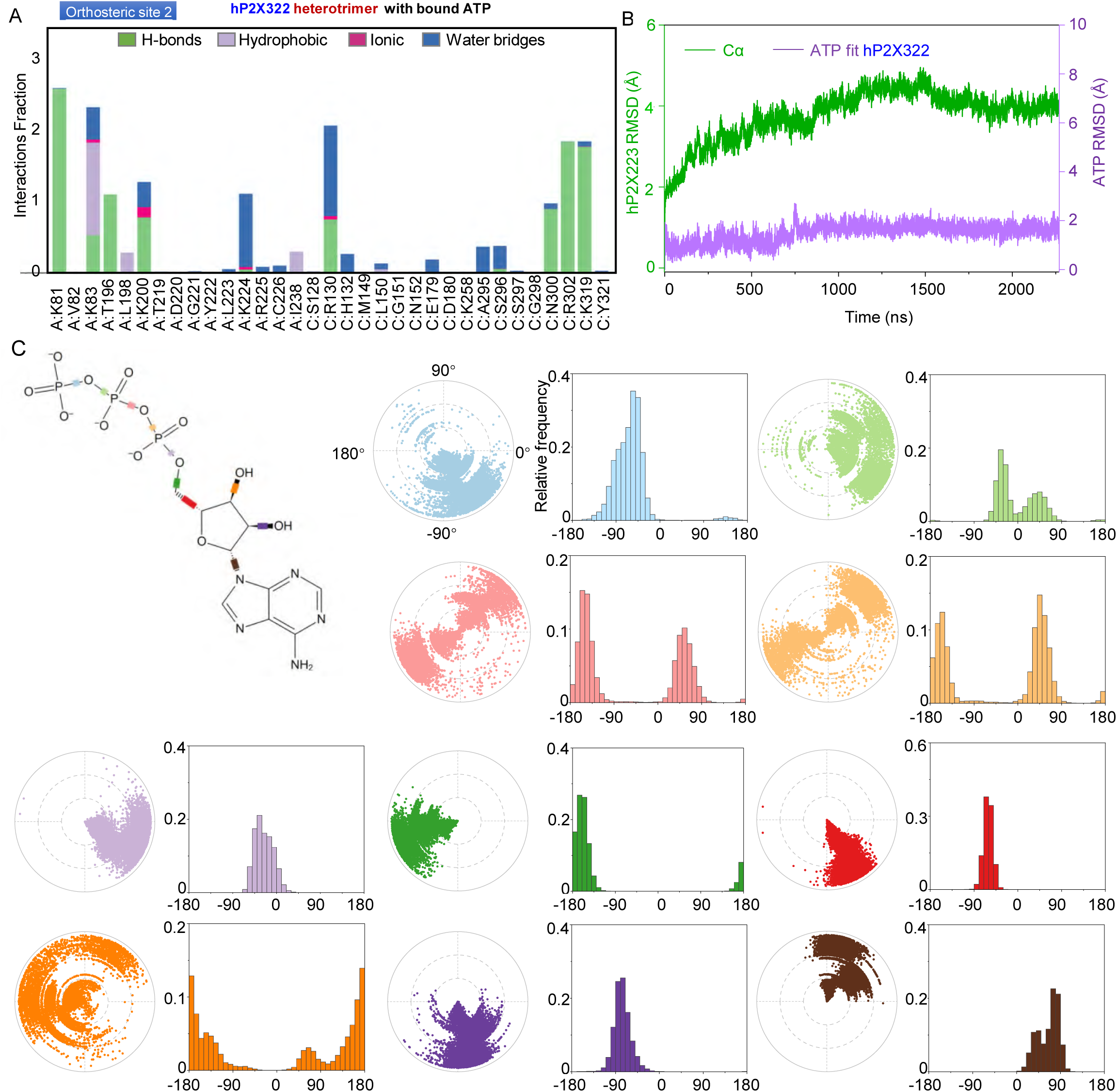
Conventional MD simulations to study the interaction of hP2X322 heterotrimer in complex with ATP at orthosteric site 2. (**A**) The interaction between hP2X322 and ATP was monitored throughout the MD simulations. hP2X322/ATP interactions were classified into four types: hydrogen bonds (green), hydrophobic (light purple), ionic interactions (pink), and water bridges (blue). The stacked histograms are normalized over the course of the trajectory. (**B**) RMSD analysis of the binding process of ATP to hP2X322 throughout the MD simulation. (**C**) Torsion plots of ATP summarizing the conformational evolution of each rotatable bond every 10 ns throughout the simulated trajectory. Two-dimensional schematics of ATP are shown as color-coded rotatable bonds. The radial plots represent the conformation of the torsion bodies. The center of the radial plot represents the beginning of the simulation, plotting the temporal evolution in the radial direction outward. The histogram summarizes the data of the corresponding radial plot, which represents the probability density of the torsion. The relationship between histogram and torsional potential gives insight into the conformational strain that the ligand underwent to maintain the hP2X322-bound conformation.

**Figure S7.**
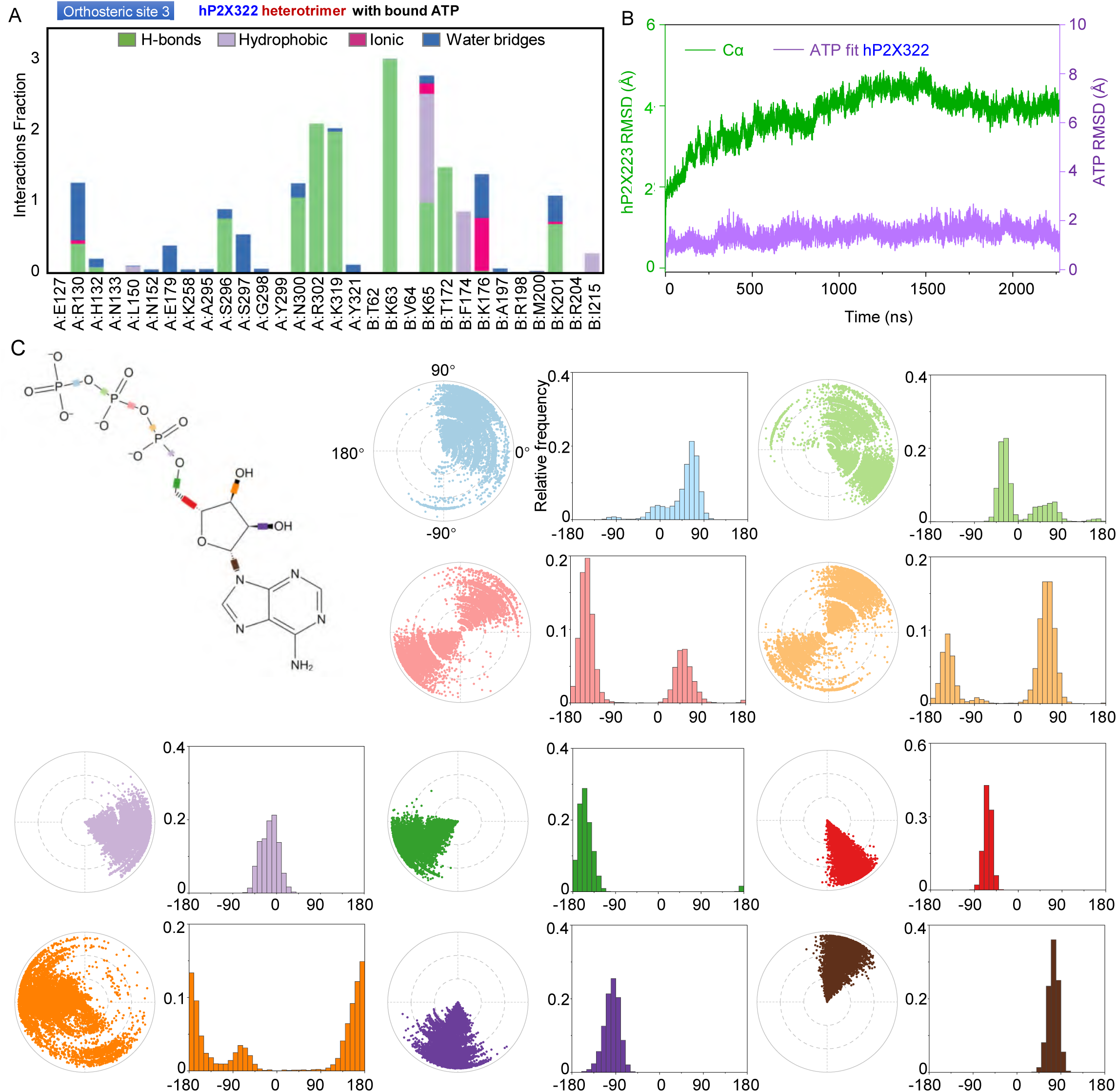
Conventional MD simulations to study the interaction of hP2X322 in complex with ATP at orthosteric site 3. (**A**) The interaction between hP2X322 and ATP was monitored throughout the MD simulations. hP2X322/ATP interactions were classified into four types: hydrogen bonds (green), hydrophobic (light purple), ionic interactions (pink), and water bridges (blue). The stacked histograms are normalized over the course of the trajectory. (**B**) RMSD analysis of the binding process of ATP to hP2X322 throughout the MD simulation. (**C**) Torsion plots of ATP summarizing the conformational evolution of each rotatable bond every 10 ns throughout the simulated trajectory. Two-dimensional schematics of ATP are shown as color-coded rotatable bonds. The radial plots represent the conformation of the torsion bodies. The center of the radial plot represents the beginning of the simulation, plotting the temporal evolution in the radial direction outward. The histogram summarizes the data of the corresponding radial plot, which represents the probability density of the torsion. The relationship between histogram and torsional potential gives insight into the conformational strain that the ligand underwent to maintain the hP2X322-bound conformation.

**Figure S8.**
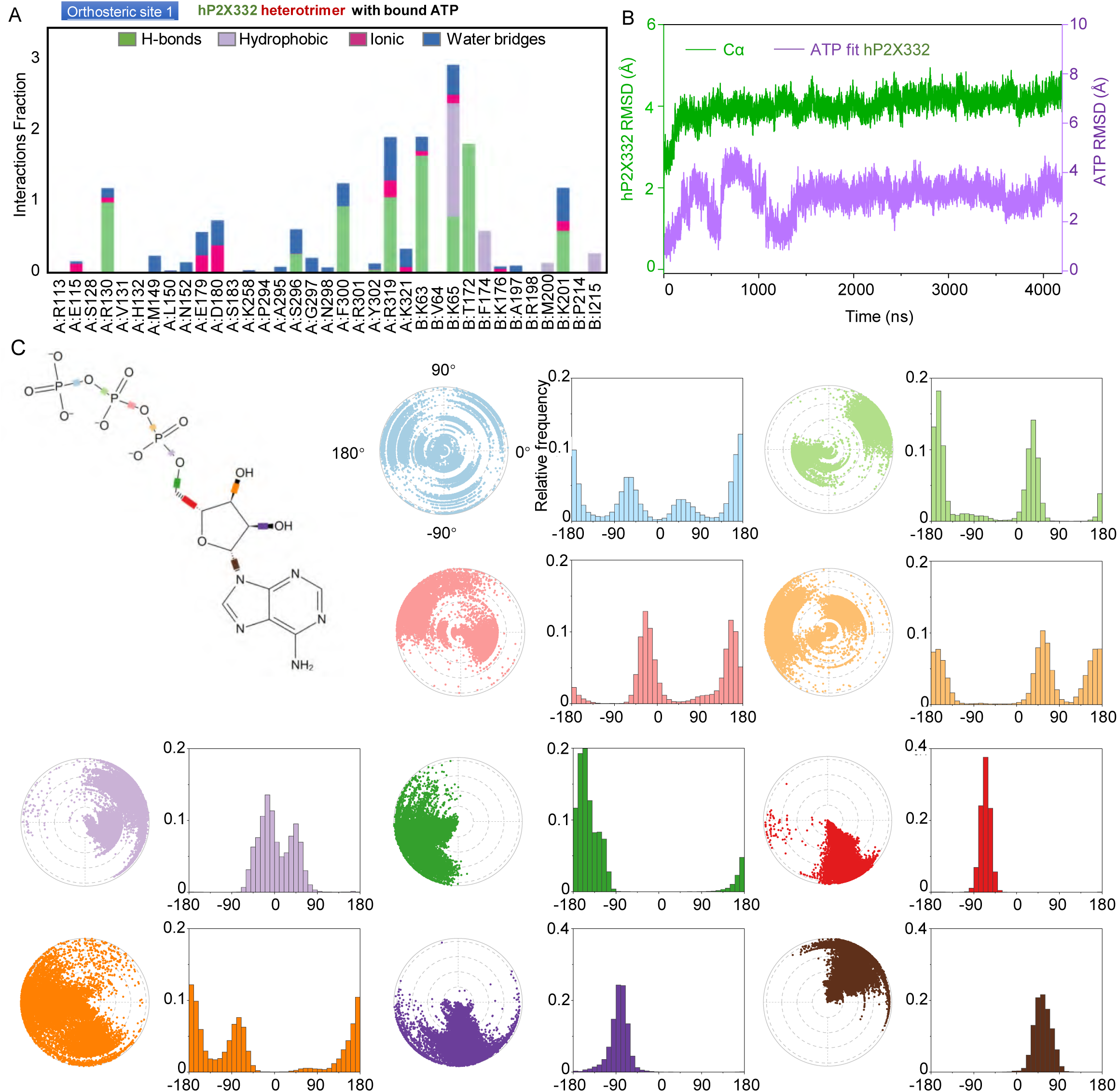
Conventional MD simulations to study the interaction of hP2X332 heterotrimer with ATP at orthosteric site 1. (**A**) The interaction between hP2X332 and ATP was monitored throughout the MD simulations. hP2X332/ATP interactions were classified into four types: hydrogen bonds (green), hydrophobic (light purple), ionic interactions (pink), and water bridges (blue). The stacked histograms are normalized over the course of the trajectory. (**B**) RMSD analysis of the binding process of ATP to hP2X332 throughout the MD simulation. (**C**) Torsion plots of ATP summarizing the conformational evolution of each rotatable bond every 10 ns throughout the simulated trajectory. Two-dimensional schematics of ATP are shown as color-coded rotatable bonds. The radial plots represent the conformation of the torsion bodies. The center of the radial plot represents the beginning of the simulation, plotting the temporal evolution in the radial direction outward. The histogram summarizes the data of the corresponding radial plot, which represents the probability density of the torsion. The relationship between histogram and torsional potential gives insight into the conformational strain that the ligand underwent to maintain the hP2X332-bound conformation.

**Figure S9.**
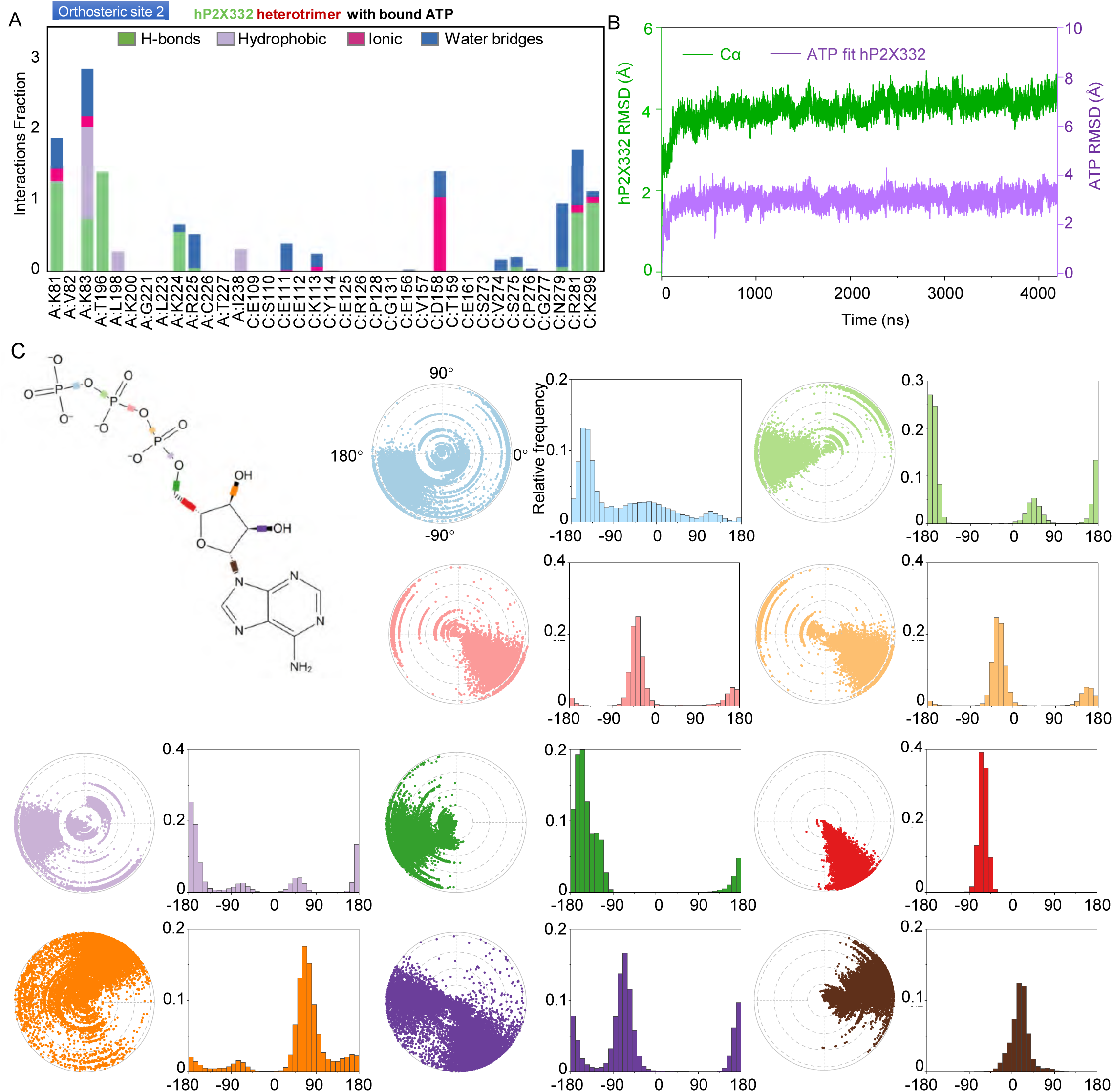
Conventional MD simulations to study the interaction of hP2X332 heterotrimer with ATP at orthosteric site 2. (**A**) The interaction between hP2X332 and ATP was monitored throughout the MD simulations. hP2X332/ATP interactions were classified into four types: hydrogen bonds (green), hydrophobic (light purple), ionic interactions (pink), and water bridges (blue). The stacked histograms are normalized over the course of the trajectory. (**B**) RMSD analysis of the binding process of ATP to hP2X332 throughout the MD simulation. (**C**) Torsion plots of ATP summarizing the conformational evolution of each rotatable bond every 10 ns throughout the simulated trajectory. Two-dimensional schematics of ATP are shown as color-coded rotatable bonds. The radial plots represent the conformation of the torsion bodies. The center of the radial plot represents the beginning of the simulation, plotting the temporal evolution in the radial direction outward. The histogram summarizes the data of the corresponding radial plot, which represents the probability density of the torsion. The relationship between histogram and torsional potential gives insight into the conformational strain that the ligand underwent to maintain the hP2X332-bound conformation.

**Figure S10.**
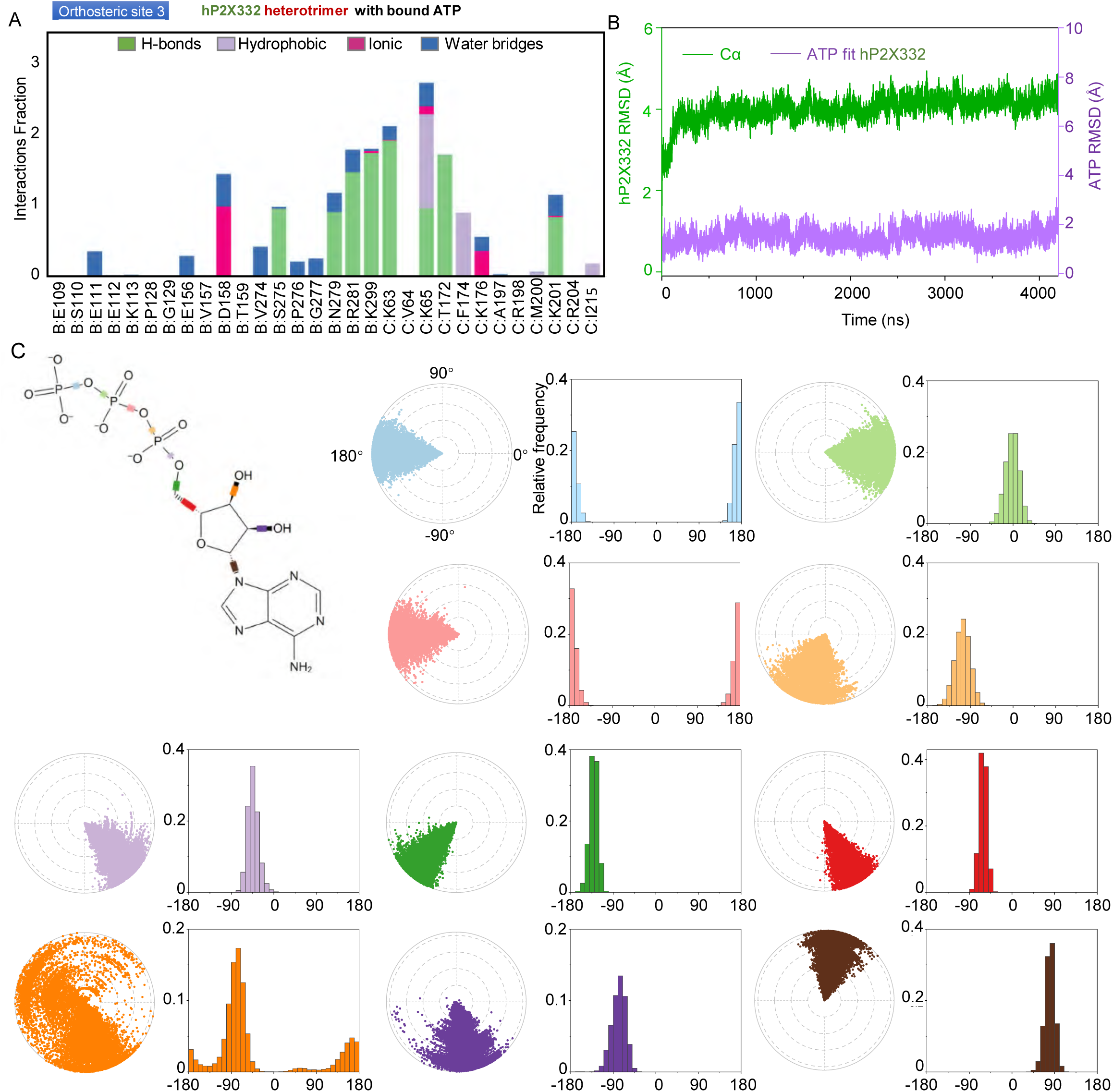
Conventional MD simulations to study the interaction of hP2X332 heterotrimer with ATP at orthosteric site 3. (**A**) The interaction between hP2X332 and ATP was monitored throughout the MD simulations. hP2X332/ATP interactions were classified into four types: hydrogen bonds (green), hydrophobic (light purple), ionic interactions (pink), and water bridges (blue). The stacked histograms are normalized over the course of the trajectory. (**B**) RMSD analysis of the binding process of ATP to hP2X332 throughout the MD simulation. (**C**) Torsion plots of ATP summarizing the conformational evolution of each rotatable bond every 10 ns throughout the simulated trajectory. Two-dimensional schematics of ATP are shown as color-coded rotatable bonds. The radial plots represent the conformation of the torsion bodies. The center of the radial plot represents the beginning of the simulation, plotting the temporal evolution in the radial direction outward. The histogram summarizes the data of the corresponding radial plot, which represents the probability density of the torsion. The relationship between histogram and torsional potential gives insight into the conformational strain that the ligand underwent to maintain the hP2X332-bound conformation.

**Figure S11.**
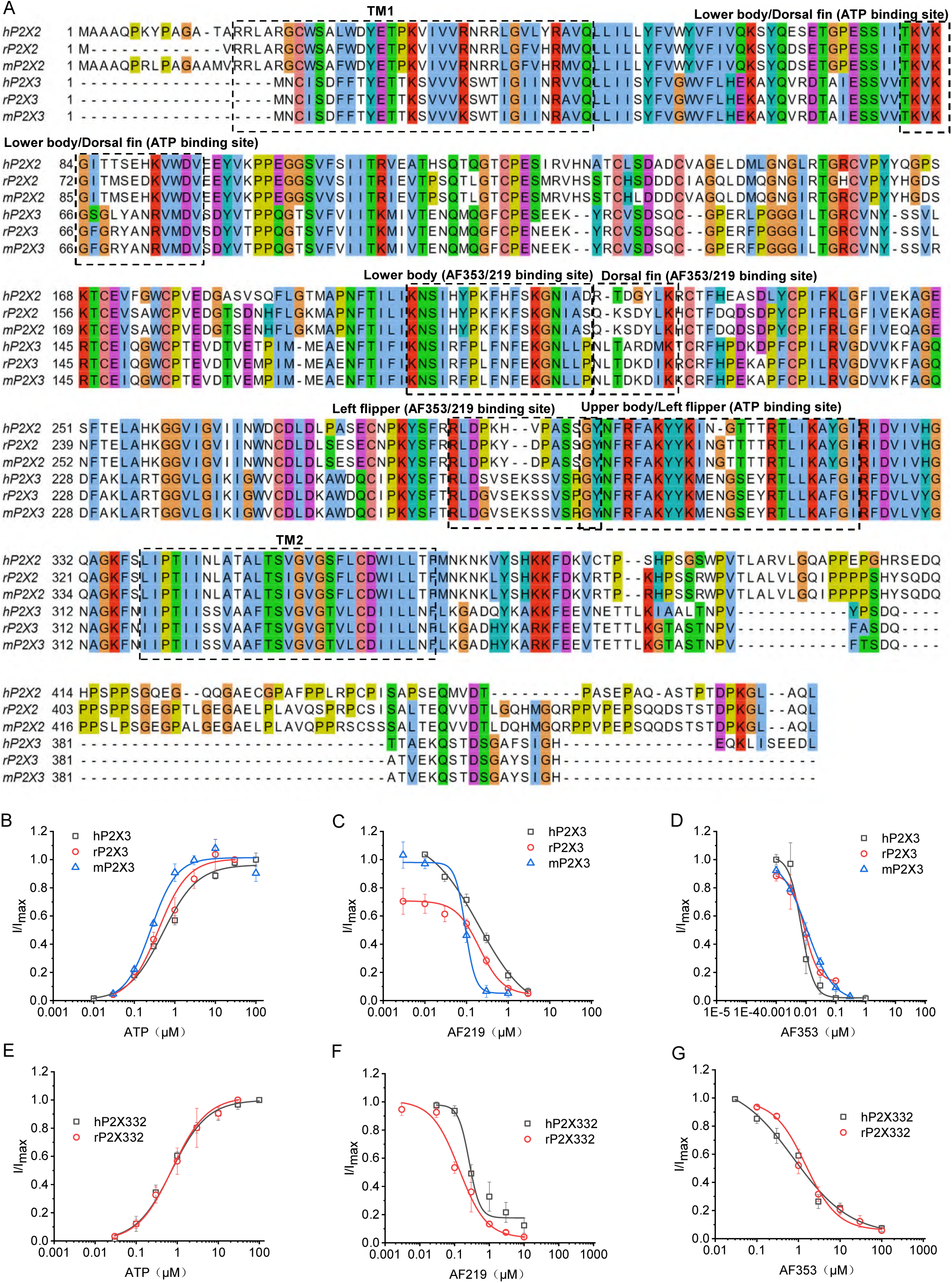
Activation and inhibition profiles of ATP, AF-219 and AF-353 on P2X3 homotrimers and P2X332 heterotrimers across species. (**A**) Sequence alignments of human P2X2 (hP2X2), rat P2X2 (rP2X2), mouse (mP2X2), hP2X3, rP2X3 and mP2X3. (**B**) Concentration-response curves of ATP on the current amplitude in hP2X3, rP2X3, and mP2X3. (**C, D**) Concentration-response curves of hP2X3, rP2X3, and mP2X3 to AF-219 (C) and AF-353 (D). Currents were evoked by 1 µM ATP for P2X3. (**E**) Concentration-response curves of ATP on the current amplitude in hP2X332 and rP2X332 heterotrimer. (**F, G**) Concentration-response curves of hP2X3332 and rP2X332 heterotrimer to AF-219 (F) and AF-353 (G). Currents were evoked by 1 µM ATP. Each solid line is a fit of the Hill equation. All summary data are expressed as mean ± SEM (n = 3-5). Source data are provided as a Source Data file. UniProt accession numbers are provided in the legend of Fig. S1.

**Figure S12.**
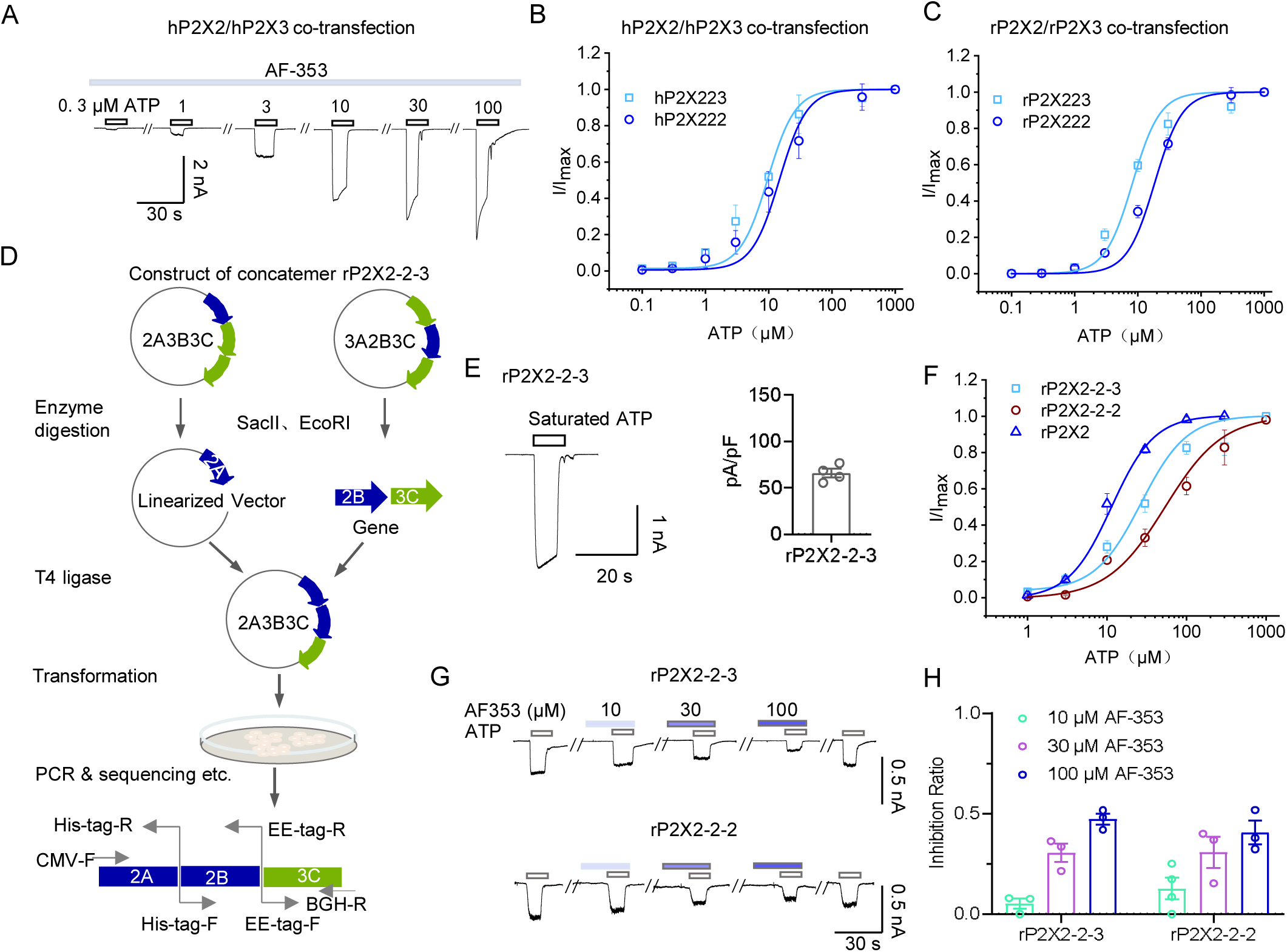
Design and validation of rat P2X2-2-3 (rP2X2-2-3) concatemers and P2X2/3 con-transfection systems. (**A**) Whole-cell patch-clamp recordings showed that AF-353 fails to inhibit the currents mediated by P2X2/3 co-transfection. (**B, C**) Concentration-response curves of ATP on the current amplitude in hP2X2, hP2X2/3 con-transfection (B), rP2X2, and rP2X2/3 con-transfection. Each solid line is a fit of the Hill 1 equation. (**D**) Construction of wild-type (WT) rP2X2-2-3 concatemers. (**E**) The current characteristics and current density of rP2X2-2-3 concatemers evoked by ATP. (**F**) Concentration-response curves of ATP on the current amplitude in rP2X2, rP2X2-2-2, and rP2X2-2-3 concatemers. (**G, H**) Representative currents (G) and pooled data inhibited by AF-353 (H) recorded from cells transfected with rP2X2-2-2 and rP2X2-2-3 concatemers. Data are shown as circles from different experimental units.

**Figure S13.**
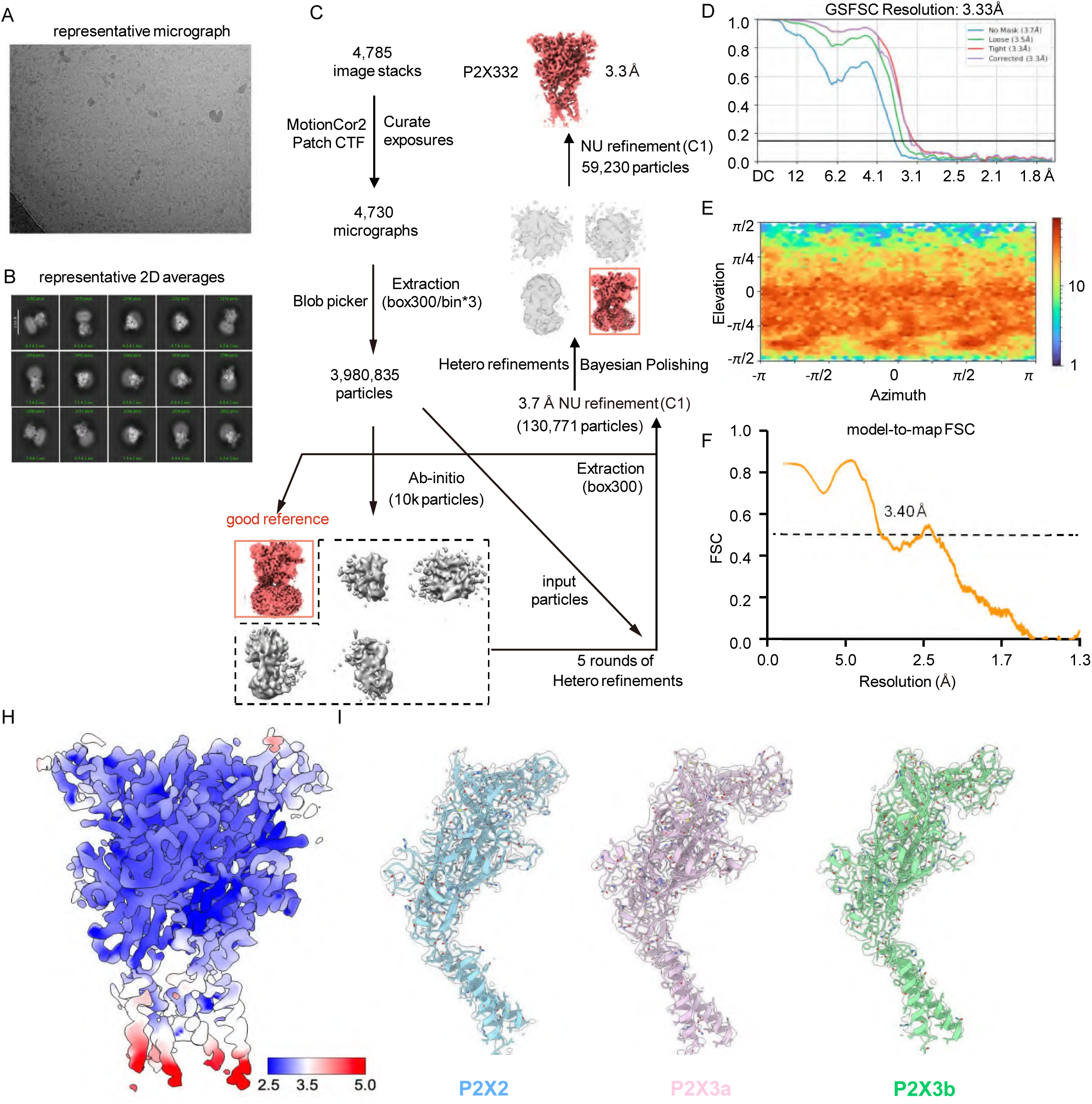
Cryo-EM analysis of the P2X2/P2X3 heteromeric complex co-expressed at a 1:10 ratio. (**A**) Representative cryo-EM micrograph. (**B**) Selected 2D class averages. (**C**) Data processing workflow for 3D reconstruction. (**D**) FSC curves from the gold-standard refinement. (**E**) Angular distribution of particles used in the final 3D reconstruction. (**F**) FSC curves between the final atomic model and the corresponding sharpened map. (**G**) Local resolution estimation of the sharpened cryo-EM map. (**H**) Representative cryo-EM density for key structural features in each subunit.

**Figure S14.**
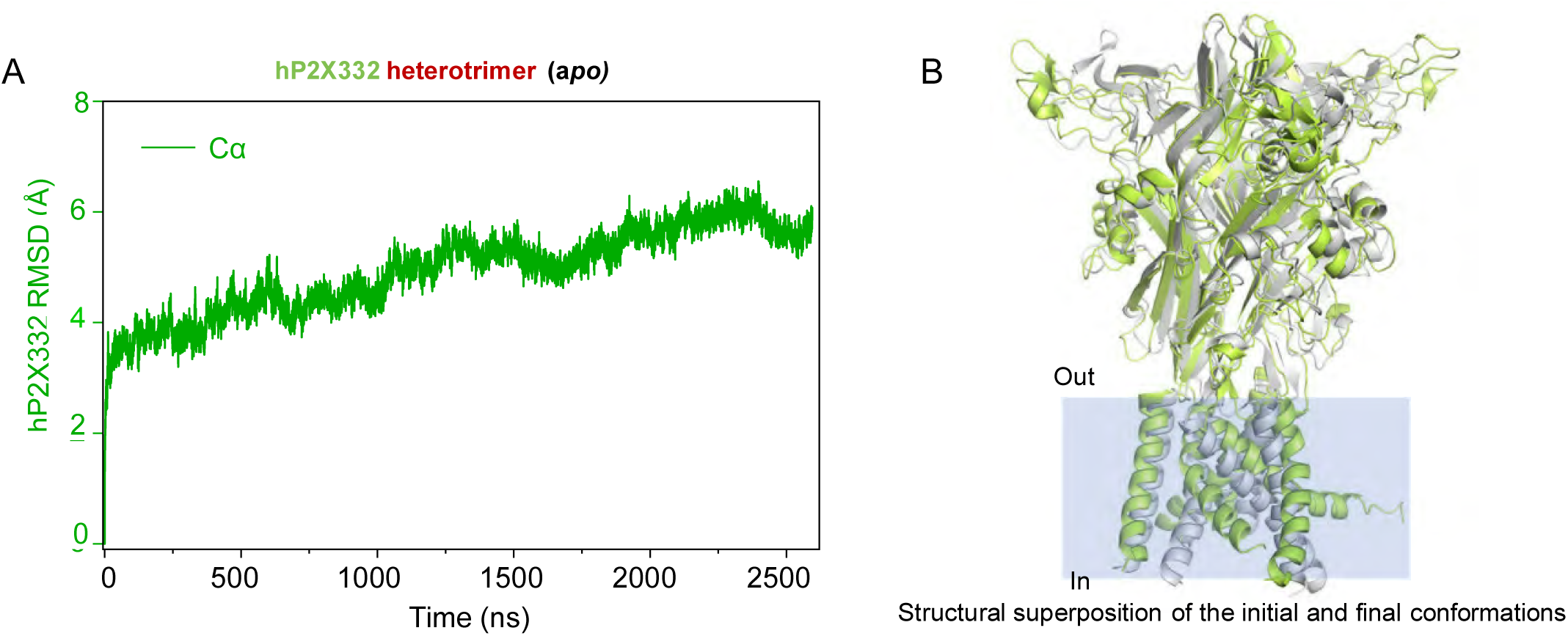
Conventional MD simulations of the *apo* hP2X332 heterotrimer. (**A**) Time evolution of the RMSD of hP2X322 throughout the simulation trajectory. (**B**) Structural superposition of the initial and final conformations, illustrating conformational changes observed during the simulation.

**Figure S15.**
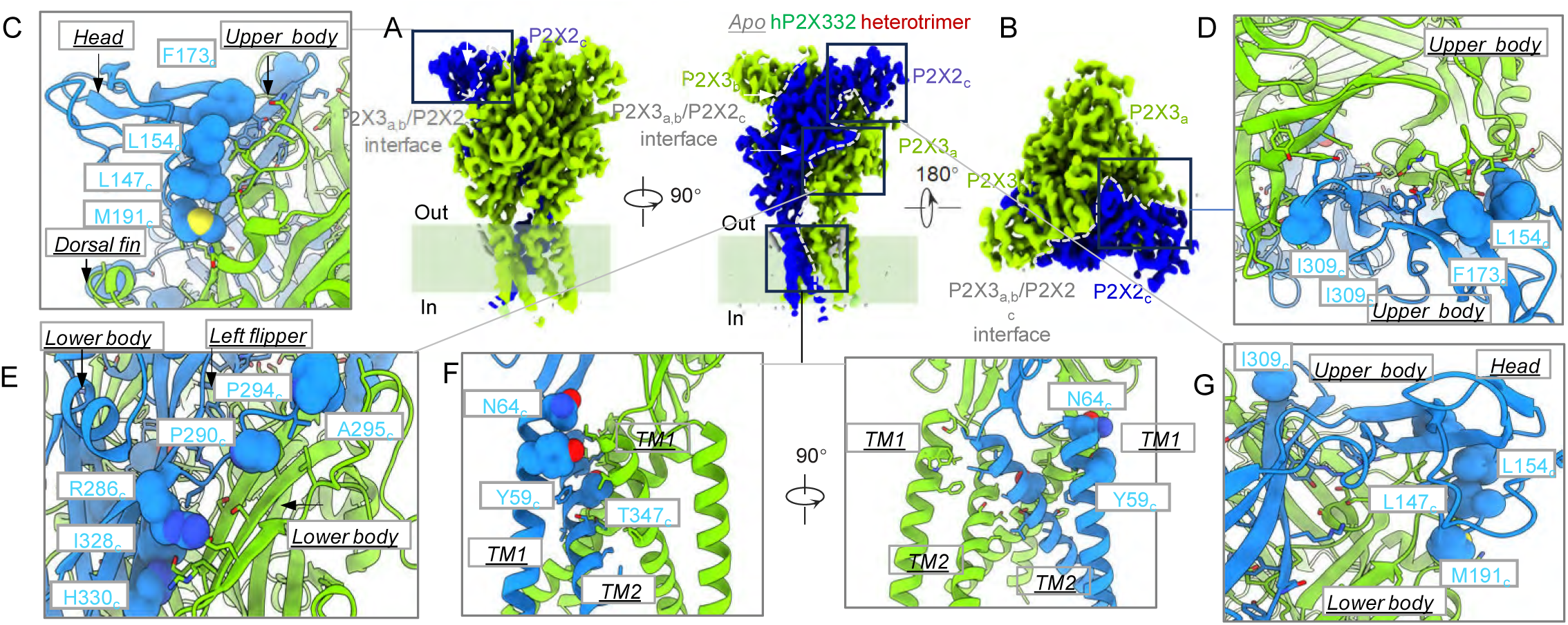
Cryo-EM structure of the apo hP2X332 heterotrimer highlighting non-conserved residues at the P2X2–P2X3 subunit interfaces. **(A, B)** Cryo-EM density map of the hP2X332 heterotrimer viewed parallel to the membrane plane (A) and from the membrane plane (B) to visualize subunit interfaces. Gray dashed lines indicate inter-subunit interfaces. **(C–G)** Residues at the subunit assembly interfaces. Interface residues are shown as sticks, with spheres indicating non-conserved residues between P2X2 and P2X3.

**Figure S16.**
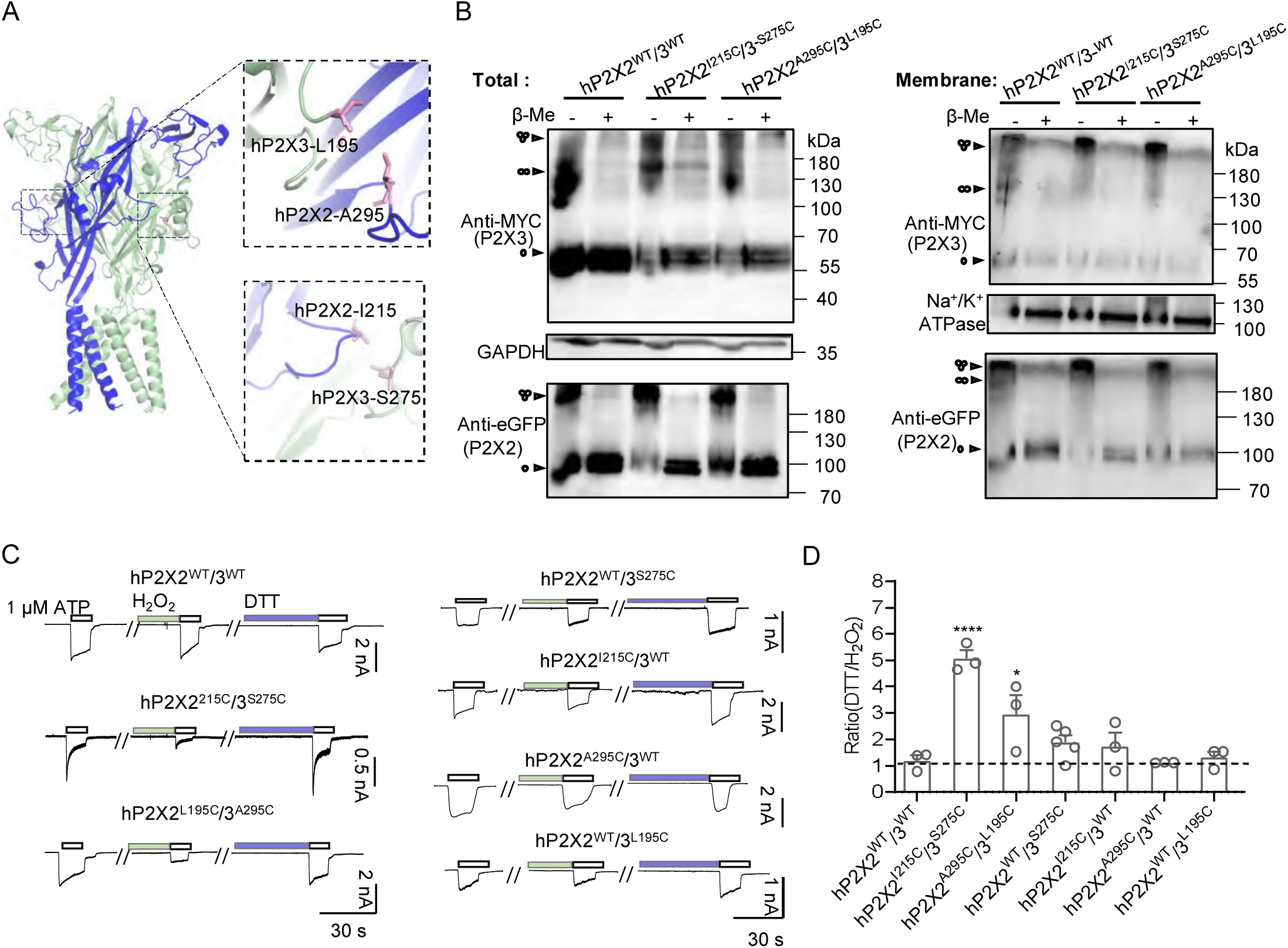
Functional role of orthosteric site 1 in the activation of the hP2X332 heterotrimer via conformational modulation rather than direct ATP binding. **(A)** Orthosteric sites 1 and 2 in the hP2X332 heterotrimer, highlighting residues targeted for cysteine cross-linking (shown as sticks). **(B)** Non-reducing Western blot analysis of heterotrimers containing double cysteine mutants I215C (DF domain, P2X2) / S275C (LF domain, P2X3) and A295C (LF domain, P2X2) / L195C (DF domain, P2X3), both predominantly forming trimers. WT channels display minor dimer formation and comparatively lower trimer levels. **(C, D)** Representative ATP-evoked currents (C) and normalized pooled data (D) from cells expressing hP2X332 mutants. Disulfide bonds were induced with 0.3% H₂O₂ and reduced with 10 mM DTT. Data in (D) represent the ratio of ATP-evoked current post- to pre-DTT treatment. Each circle indicates an independent cell. Statistical analysis: one-way ANOVA with Dunnett’s multiple comparisons test (F(6,16) = 13.00); *p < 0.005, ***p < 0.001 versus hP2X2^WT^/3^WT^. Source data are provided.

**Figure S17.**
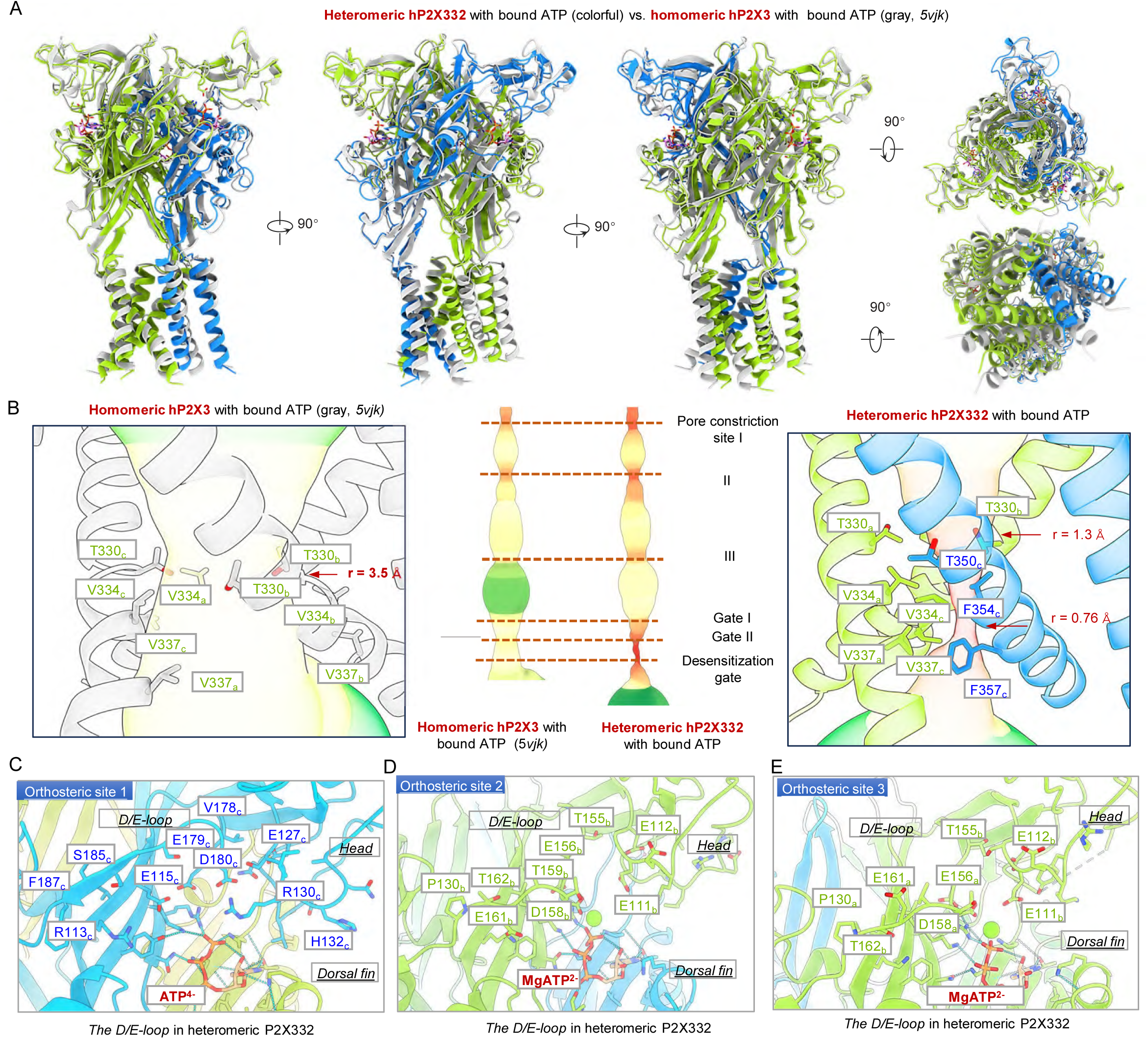
Structural comparison of ATP-bound desensitized heteromeric hP2X332 and ATP-bound open-state homomeric hP2X3. **(A)** Superimposition of the heteromeric hP2X332 structure (colored, this study) with the homomeric hP2X3 ATP-bound structure (gray, PDB 5VJK). **(B)** Close-up views of the channel gate: heteromeric hP2X332 in the desensitized state (right) and homomeric hP2X3 in the open state (left). The central pore was analyzed using MOLE. Structural alignment shows that the extracellular domain (ECD) conformation of the desensitized heterotrimer closely resembles that of the open-state homomer, as also indicated by overlapping pore constriction sites in the ECD. **(C–E)** Structural details of the D/E-loop at the three orthosteric ATP-binding sites in hP2X332. ATP molecules and D/E-loop residues are depicted as sticks. The D/E-loop, in conjunction with head domain residues, defines the chemical environment of each orthosteric site, dictating whether the pocket preferentially binds ATP⁴⁻ or MgATP²⁻.

**Figure S18.**
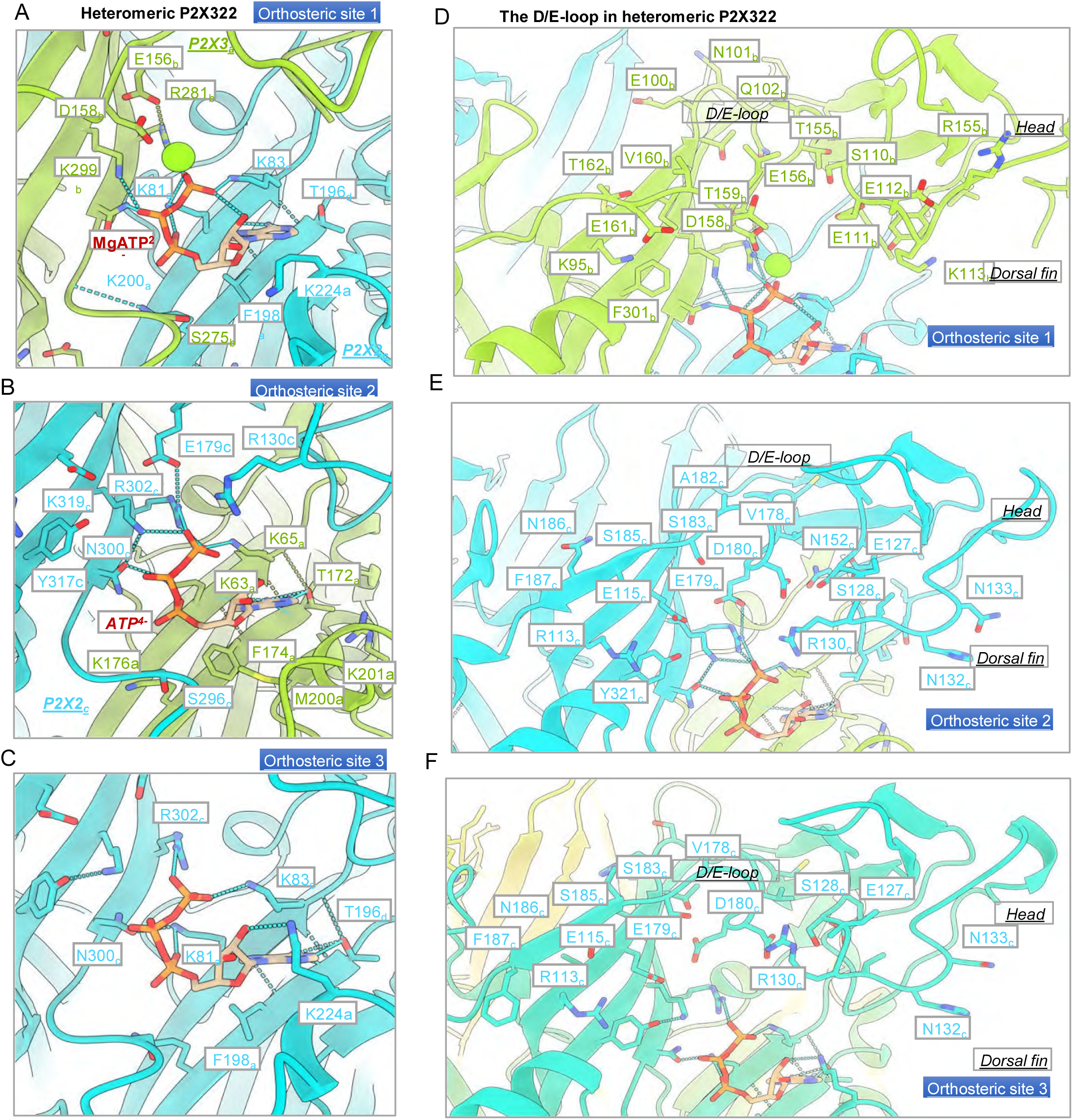
Structural analysis of the three orthosteric ATP-binding sites (A–C) and the D/E-loop (D–F) in the hP2X322 heterotrimer. ATP and critical residues are depicted as sticks for clarity. Mg²⁺ ions are highlighted by green circles in panels A and D. ATP adopts the MgATP²⁻ form in orthosteric site 1 (A) and ATP⁴⁻ in sites 2 and 3 (B, C, E, F). The D/E-loop, in combination with head domain residues, establishes the chemical properties of each orthosteric pocket, dictating whether the pocket preferentially binds ATP⁴⁻ or MgATP²⁻.

**Figure S19.**
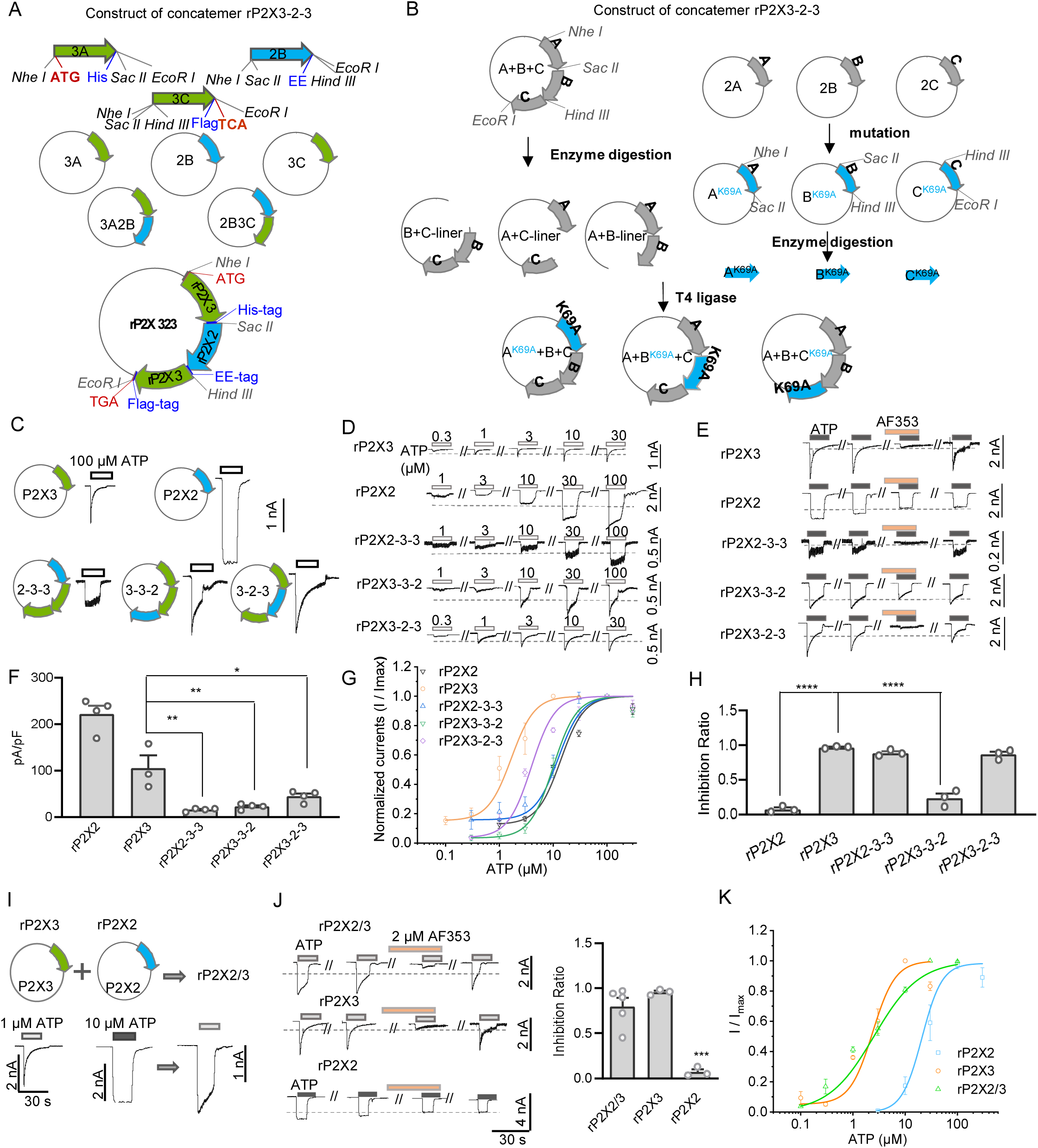
Design and validation of rP2X3-2-3 concatemers and rP2X3/2 con-transfection systems. (**A, B**) Construction of WT (A) and mutated (B) rP2X3-2-3 concatemers. (**C**) The current characteristics of homomeric rP2X2 and homomeric rP2X3, and rP2X3-2-3 rP2X3-3-2, and rP2X2-3-3 concatemers evoked by ATP. (**D, E**) Representative currents evoked by ATP (D) and inhibited by AF-353 (E) recorded from cells transfected with rP2X2, rP2X3, rP2X3-2-3, rP2X3-3-2, and rP2X2-3-3 concatemers. (**F**) Current density of rP2X2, rP2X3, rP2X3-2-3, rP2X3-3-2, and rP2X2- 3-3 concatemers evoked by 1 mM ATP. Data are shown as circles from different experimental units. *: p < 0.05, **: p < 0.01, one-way ANOVA with Dunnett’s multiple comparison test (F (4, 14) = 45.77). (**G**) Concentration-response curves of ATP on the current amplitude in rP2X2, rP2X3, rP2X3-2-3, rP2X3-3-2, and rP2X2-3-3 concatemers. Each solid line is a fit of the Hill 1 equation. (**H**) Inhibition ratio of AF-353 on rP2X2, rP2X3, rP2X3-2-3, rP2X3-3-2, and rP2X2-3-3 concatemers evoked by 1 mM ATP. Data are shown as circles from different experimental units. ***: p < 0.01, one-way ANOVA with Dunnett’s multiple comparison test (F (4, 10) = 117.0). (**I**) Construction (upper) and current characteristics (lower) of rP2X3/2 (con-transfection). (**J**) Representative current traces (left) and pooled data (right) illustrating the inhibitory effects of AF-353 on the activation of rP2X2, rP2X3, and rP2X3/2 heterotrimer (con-transfection). Each circle represents an independent cell. (**K**) Concentration-response curves of ATP on the current amplitude in rP2X2, rP2X3 and rP2X3/2 (con-transfection). Each solid line is a fit of the Hill 1 equation.

**Figure S20.**
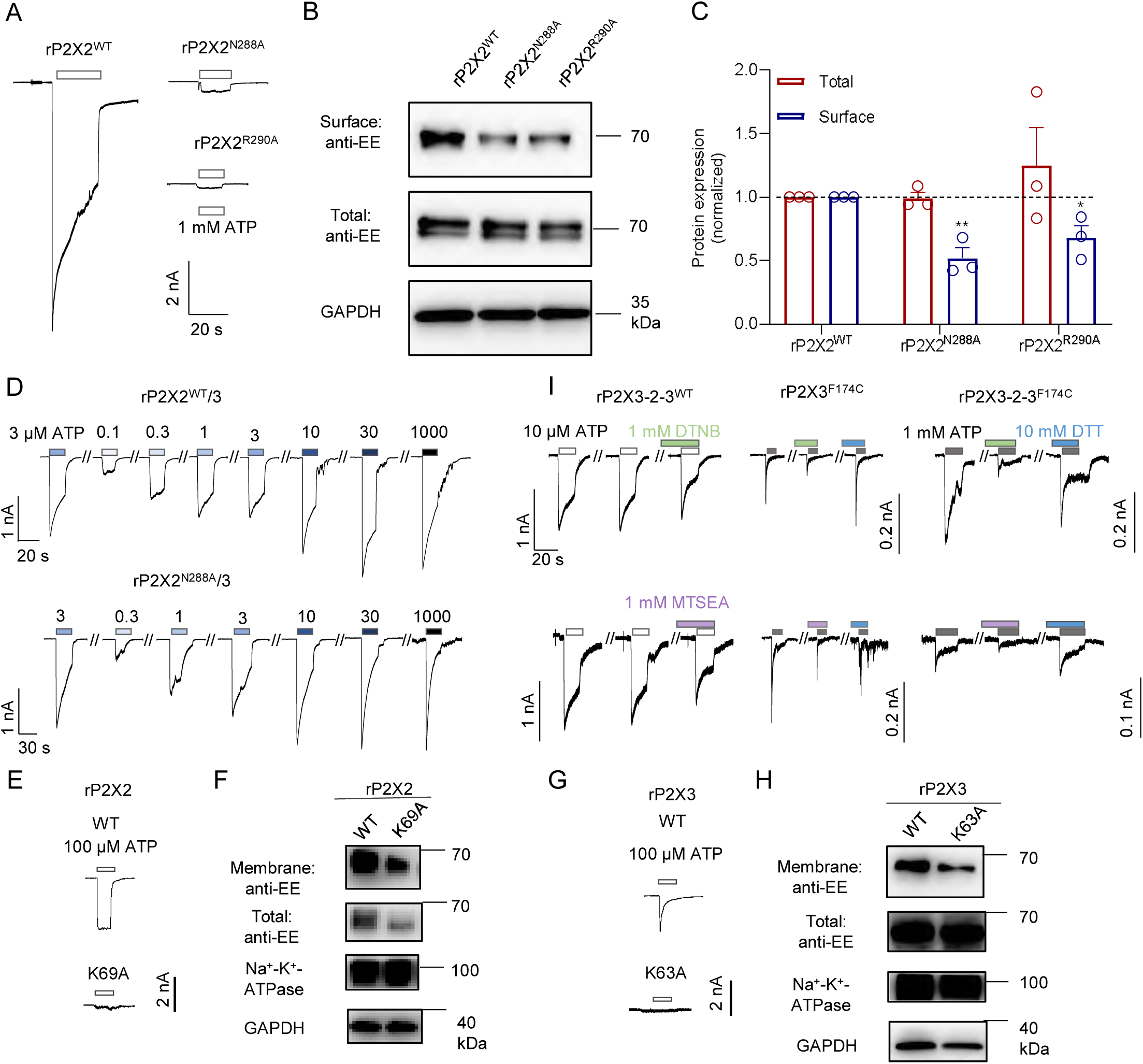
Effects of orthosteric site mutations on protein expression and receptor activation. **(A)** Representative current traces of WT rP2X2 and its mutants in response to 1 mM ATP. **(B, C)** Expression levels (B) and pooled data (C) of WT rP2X2 and its mutants in total proteins and surface proteins. Data are shown as circles from different experimental units. *p < 0.05, **p < 0.01 compared to rP2X2^WT^, one-way ANOVA with Dunnett’s multiple comparison test (F (2, 6) = 11.23). **(D)** Representative current traces of rP2X3/2 and its mutants. **(E)** Representative current traces of WT rP2X2 and its mutants in response to 100 μM ATP. **(F)** Expression levels of WT rP2X2 and its mutants in total proteins and surface proteins. **(G)** Representative current traces of WT rP2X3 and its mutants in response to 100 μM ATP. **(H)** Expression levels of WT rP2X3 and its mutants in total proteins and surface proteins. **(I)** Representative current traces of DTNB and MTSEA modification at cysteine mutation sites on rP2X3-2-3 concatemers.

**Figure S21.**
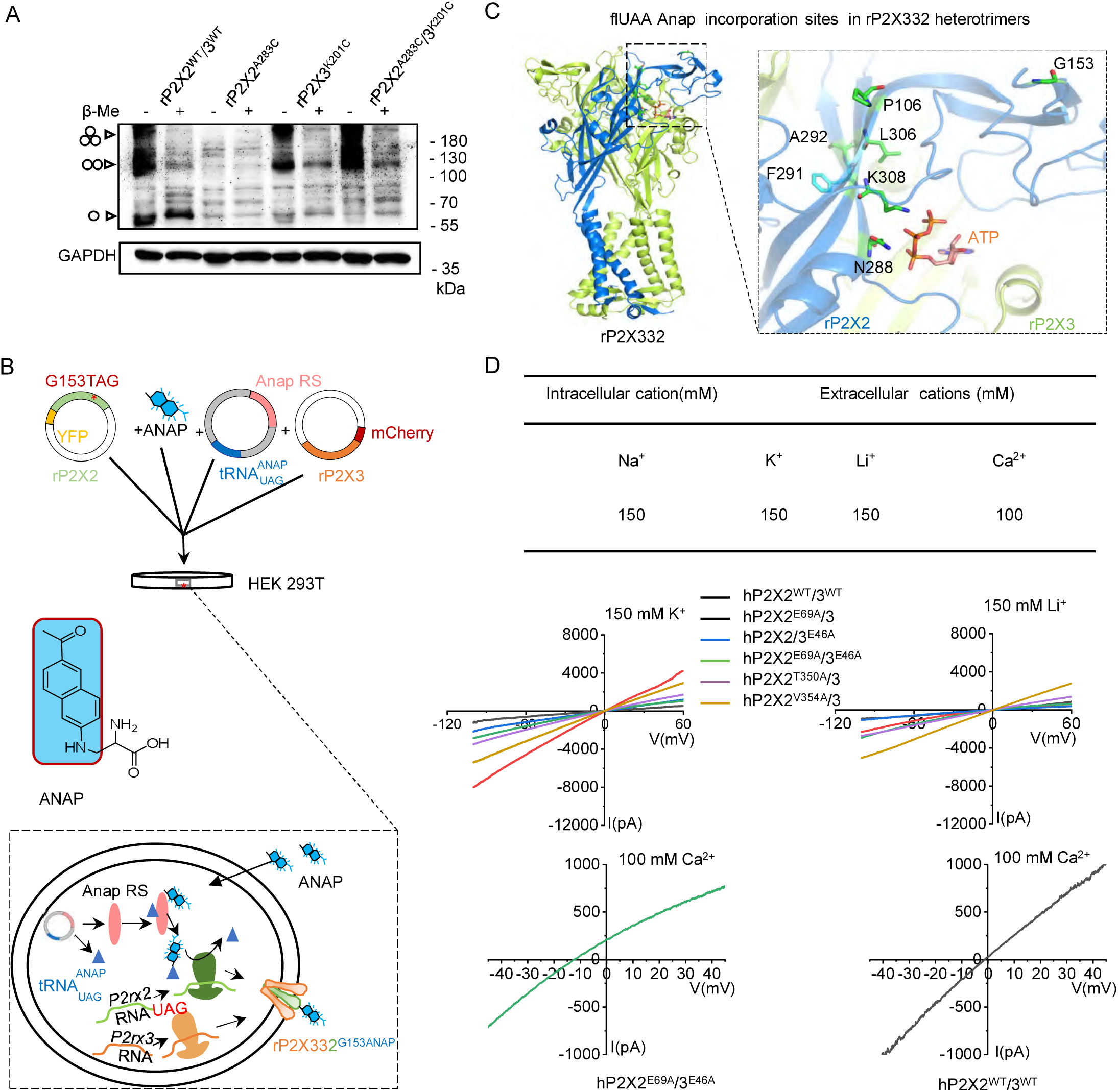
Design and validation of disulfide cross-linking, voltage-clamp fluorometry (VCF), and ion permeation assays. (**A**) Hetero-oligomers of rP2X332 containing double cysteine mutations A283C (left flipper domain, P2X2 subunit) and K201C (dorsal fin domain, P2X3 subunit) predominantly form trimers or dimers in non-reducing Western blots. (**B**) Schematic of fluorescent unnatural amino acid (flUAA) incorporation and VCF. The chemical structure of L-ANAP and the strategy for its incorporation into P2X3 are shown. Plasmids encoding ANAP tRNA synthetase and cognate tRNAs, together with plasmids carrying the receptor gene, were transiently transfected into HEK293T cells, which were then cultured in ANAP-containing medium for ≥24 h to ensure site-specific integration. (**C**) Location of selected sites at orthosteric site 1 of rP2X3₂₂ and corresponding negative control. (**D**) Solution compositions, formulas for calculating K⁺, Li⁺, and Ca²⁺ permeabilities, and a step-voltage protocol from −100 mV to +60 mV used to generate current–voltage (I–V) relationships for ion permeation assays.

**Figure S22.**
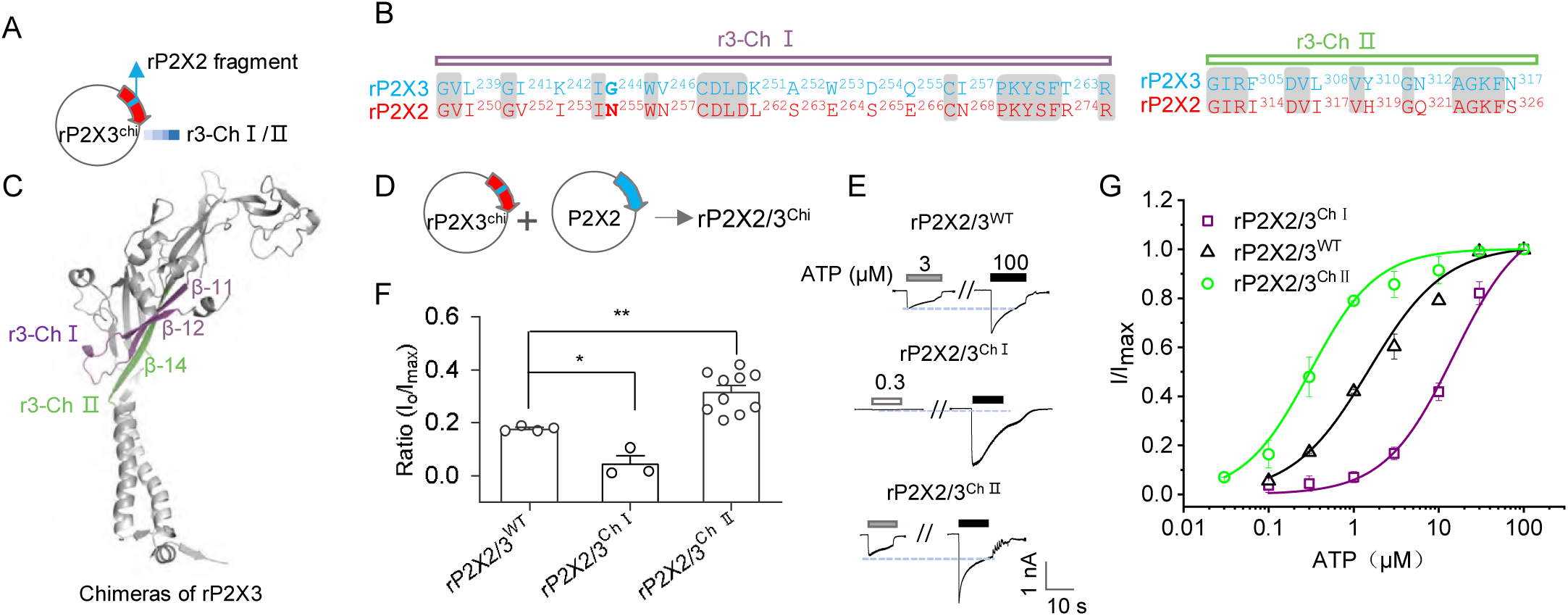
Contributions of subunit assembly interfaces to gating efficiency in the hP2X332 heterotrimer. **(A)** Schematic of rP2X3 chimeras. **(B)** Sequence alignments of rP2X2 and rP2X3. Identical residues are highlighted in gray. **(C)** Positions of Chimera I (Chi I) and Chimera II (Chi II) on rP2X3. **(D)** Schematic of the rP2X2/3 chimera (rP2X2/3^Chi^) heterotrimer. **(E, F)** Representative currents (E) and pooled data (F) from cells expressing rP2X2/3^WT^ and rP2X2/3^Chi^ heterotrimers. *p < 0.05, **p < 0.01, one-way ANOVA with Tukey’s multiple comparisons test (F(2,14) = 22.64). **(G)** Concentration-response curves of ATP for rP2X2/3^WT^ and rP2X2/3^Chi^ heterotrimers.

**Figure S23.**
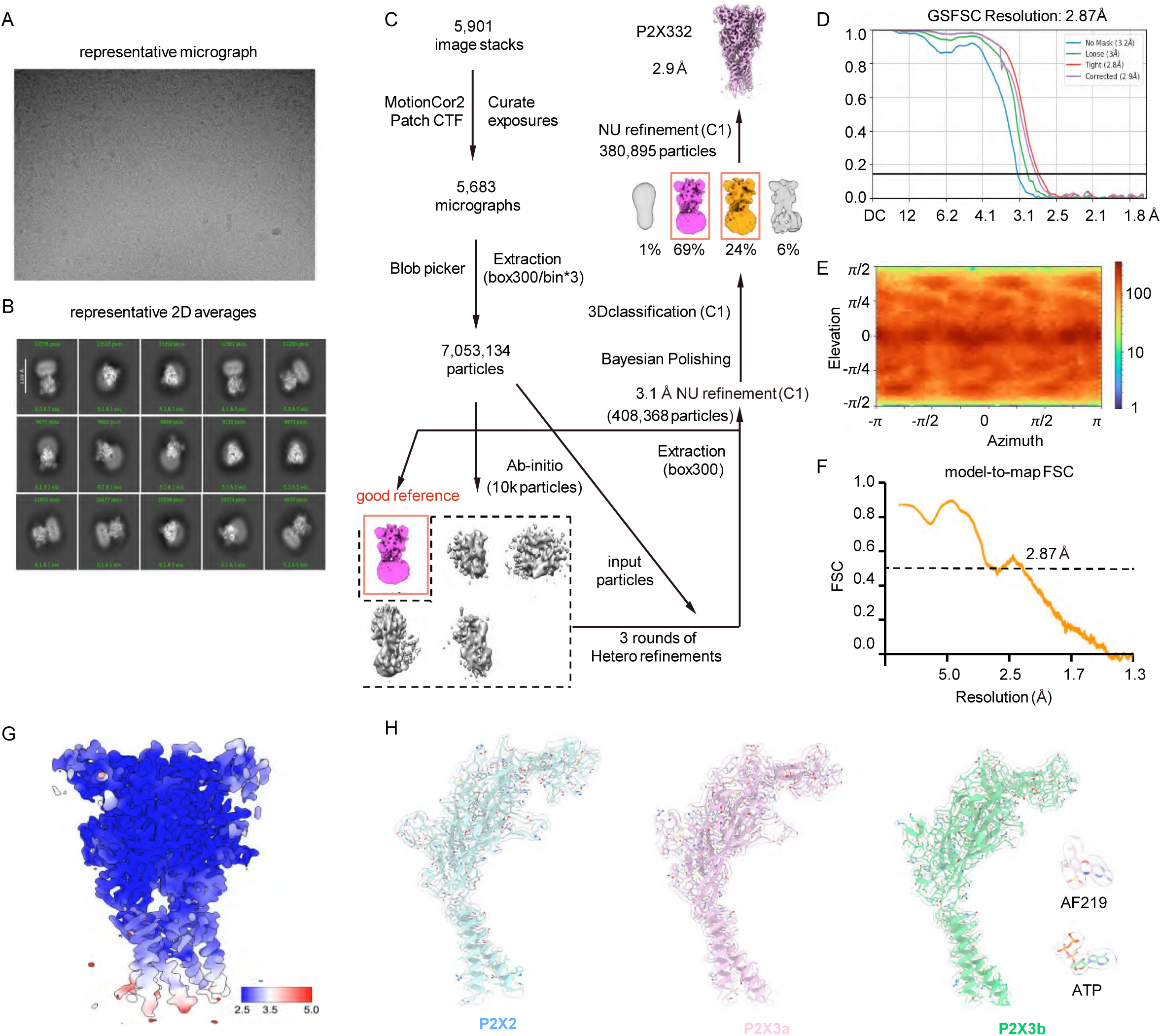
Cryo-EM analysis of the P2X2/P2X3 heteromeric complex co-expressed at a 1:10 ratio in the presence of inhibitor AF-219. (**A**) Representative cryo-EM micrograph. (**B**) Selected 2D class averages. (**C**) Data processing workflow for 3D reconstruction. (**D**) Fourier shell correlation (FSC) curves from the gold-standard refinement. (**E**) Angular distribution of particles used in the final 3D reconstruction. (**F**) FSC curves between the final atomic model and the corresponding sharpened map. (**G**) Local resolution estimation of the sharpened cryo-EM map. (**H**) Representative cryo-EM density for key structural features in each subunit, ATP and AF-219.

**Figure S24.**
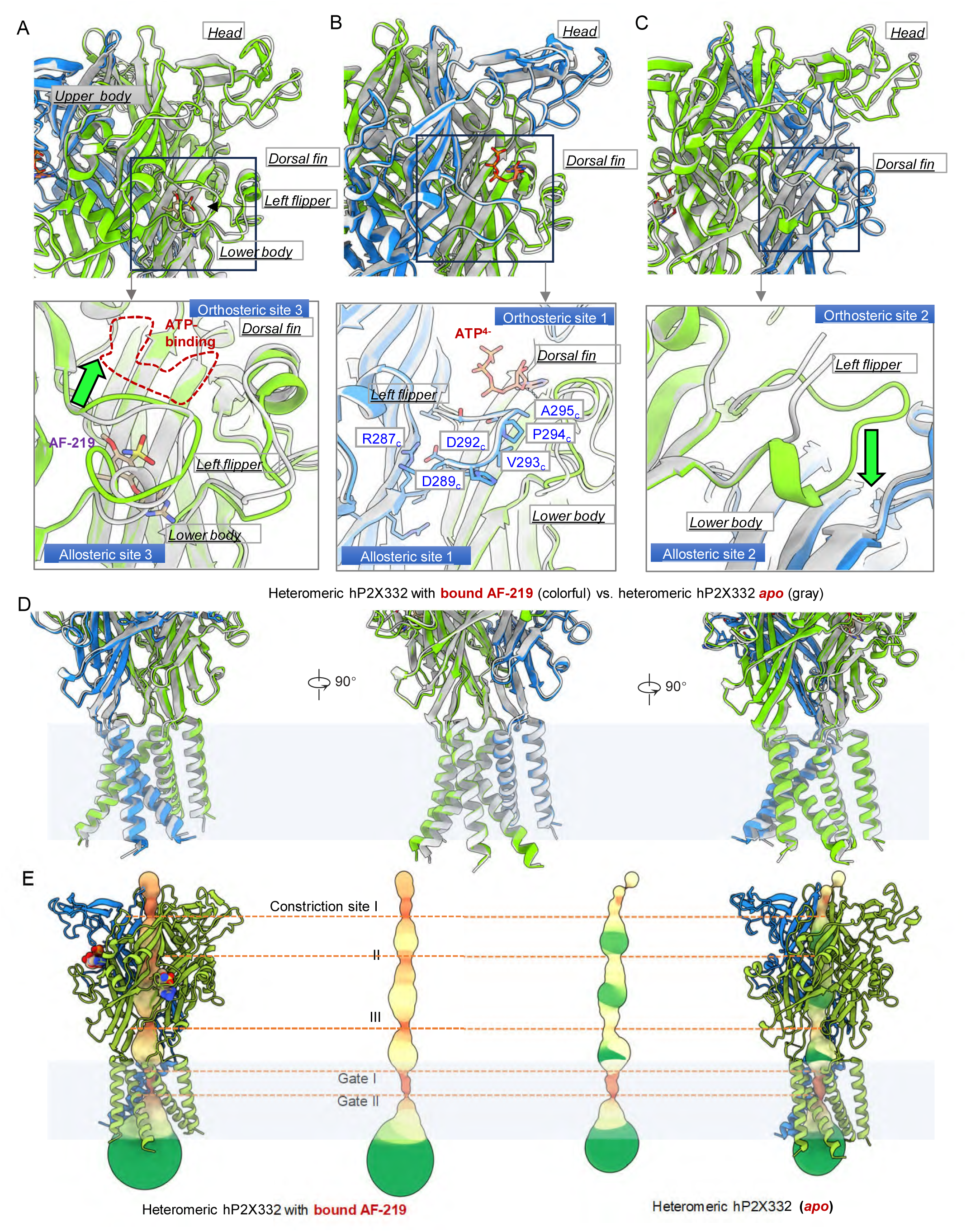
Structural comparison between AF-219-bound and apo hP2X3332 heterotrimers. (**A–C**) Detailed views of the three allosteric binding sites in hP2X332, with AF-219 and critical residues shown as sticks for clarity. (**D**) Superimposition of the AF-219-bound heterotrimer (color) and the apo heterotrimer (gray) to illustrate conformational changes.(**E**) Close-up views of the channel gate in the AF-219-bound (left) and apo (right) states. Channel geometry was calculated using the MOLE program.

**Figure S25.**
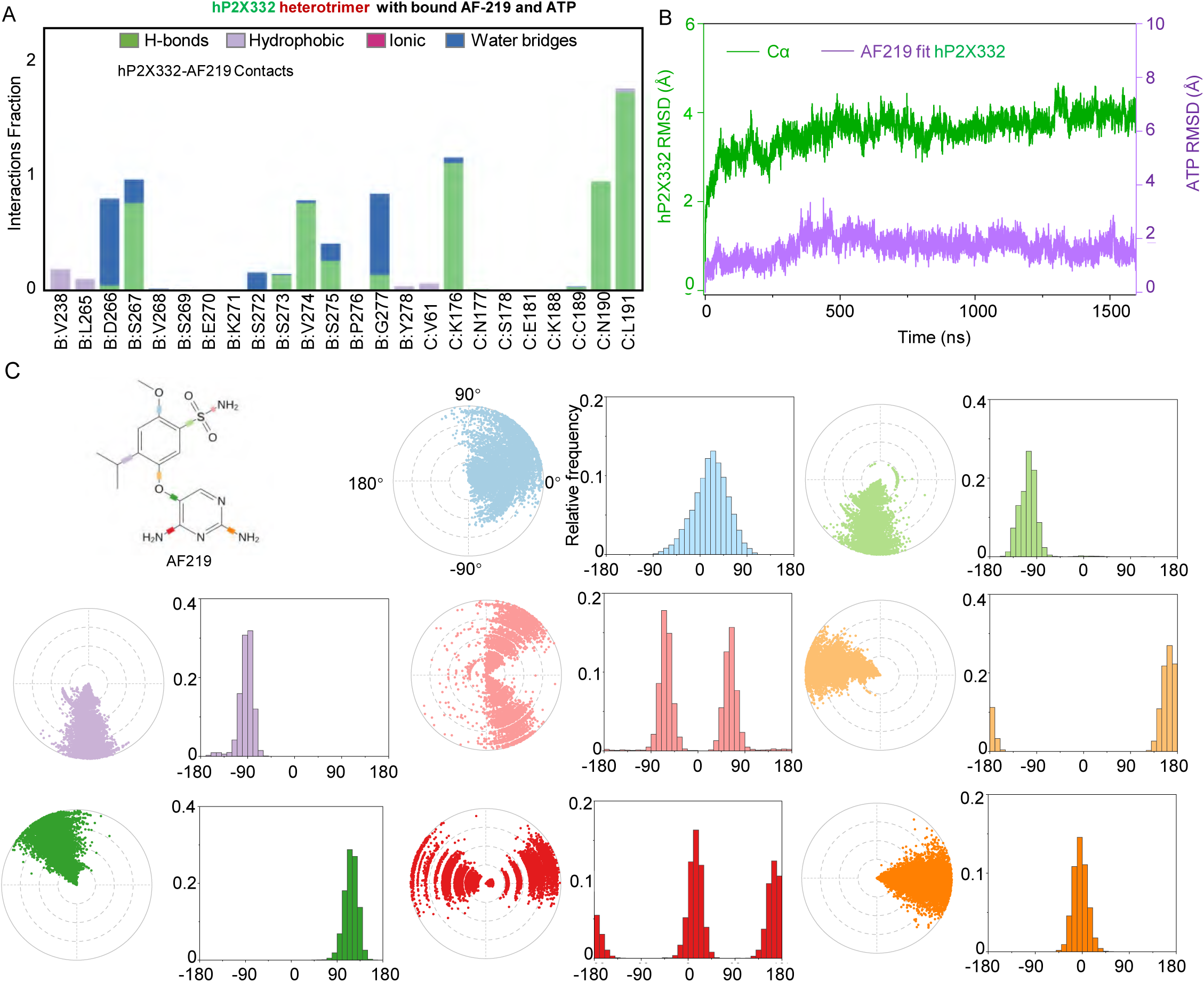
Conventional MD simulations to study the interaction of hP2X332 heterotrimer with AF-219. (**A**) The interaction between hP2X332 and AF-219 was monitored throughout the MD simulations. hP2X332/ AF-219 interactions were classified into four types: hydrogen bonds (green), hydrophobic (light purple), ionic interactions (pink), and water bridges (blue). The stacked histograms are normalized over the course of the trajectory. (**B**) RMSD analysis of the binding process of AF-219 to hP2X332 throughout the MD simulation. (**C**) Torsion plots of AF-219 summarizing the conformational evolution of each rotatable bond every 10 ns throughout the simulated trajectory. 2D schematics of ATP are shown as color-coded rotatable bonds. The radial plots represent the conformation of the torsion bodies. The center of the radial plot represents the beginning of the simulation, plotting the temporal evolution in the radial direction outward. The histogram summarizes the data of the corresponding radial plot, which represents the probability density of the torsion. The relationship between histogram and torsional potential gives insight into the conformational strain that the ligand underwent to maintain the hP2X332-bound conformation.

**Figure S26.**
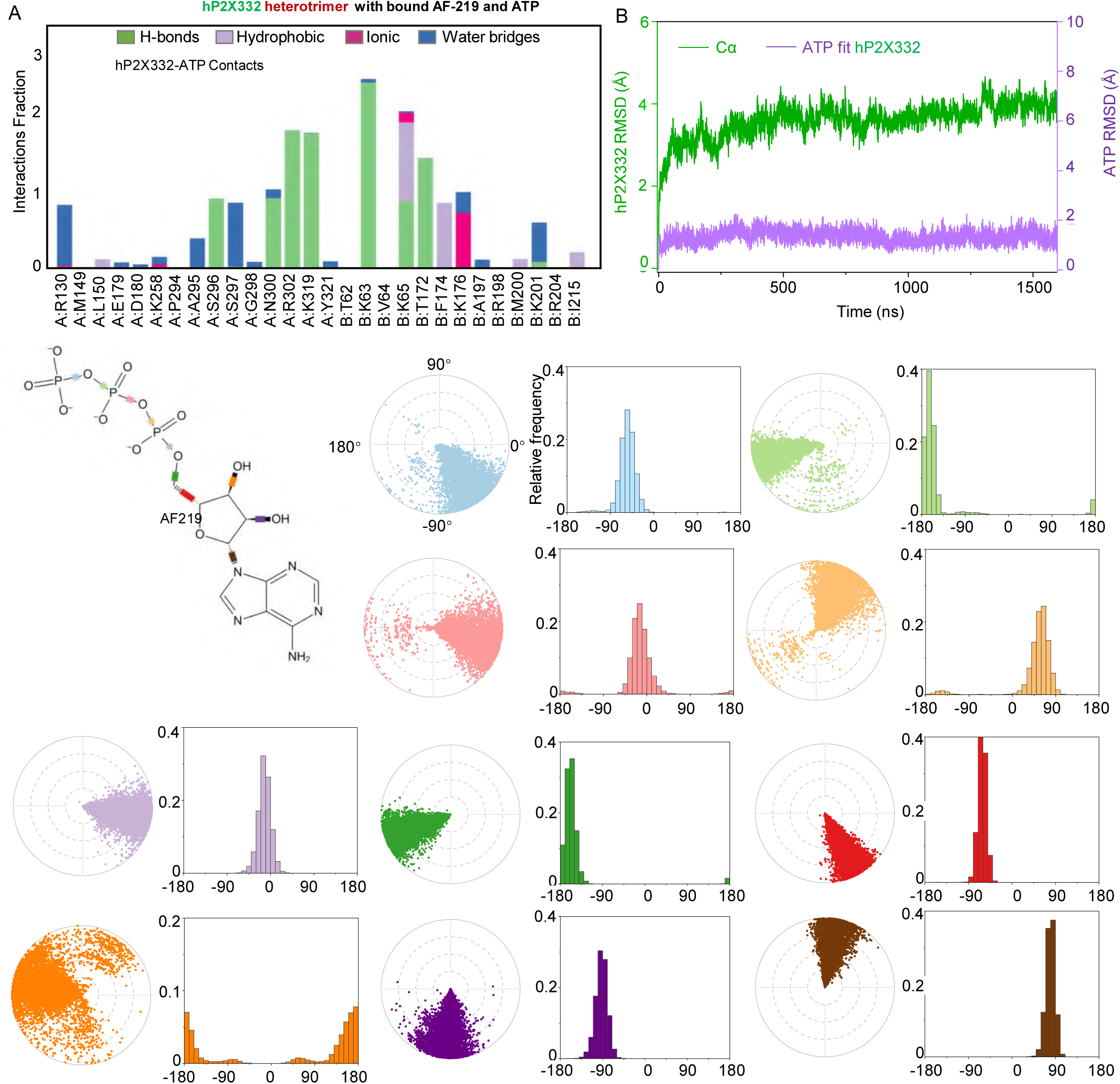
Conventional MD simulations to study the interaction of hP2X332 heterotrimer with ATP. (**A**) The interaction between hP2X332 and ATP was monitored throughout the MD simulations. hP2X332/ATP interactions were classified into four types: hydrogen bonds (green), hydrophobic (light purple), ionic interactions (pink), and water bridges (blue). The stacked histograms are normalized over the course of the trajectory. (**B**) RMSD analysis of the binding process of ATP to hP2X332 throughout the MD simulation. (**C**) Torsion plots of ATP summarizing the conformational evolution of each rotatable bond every 10 ns throughout the simulated trajectory. Two-dimensional schematics of ATP are shown as color-coded rotatable bonds. The radial plots represent the conformation of the torsion bodies. The center of the radial plot represents the beginning of the simulation, plotting the temporal evolution in the radial direction outward. The histogram summarizes the data of the corresponding radial plot, which represents the probability density of the torsion. The relationship between histogram and torsional potential gives insight into the conformational strain that the ligand underwent to maintain the hP2X332-bound conformation.

**Table S1.**
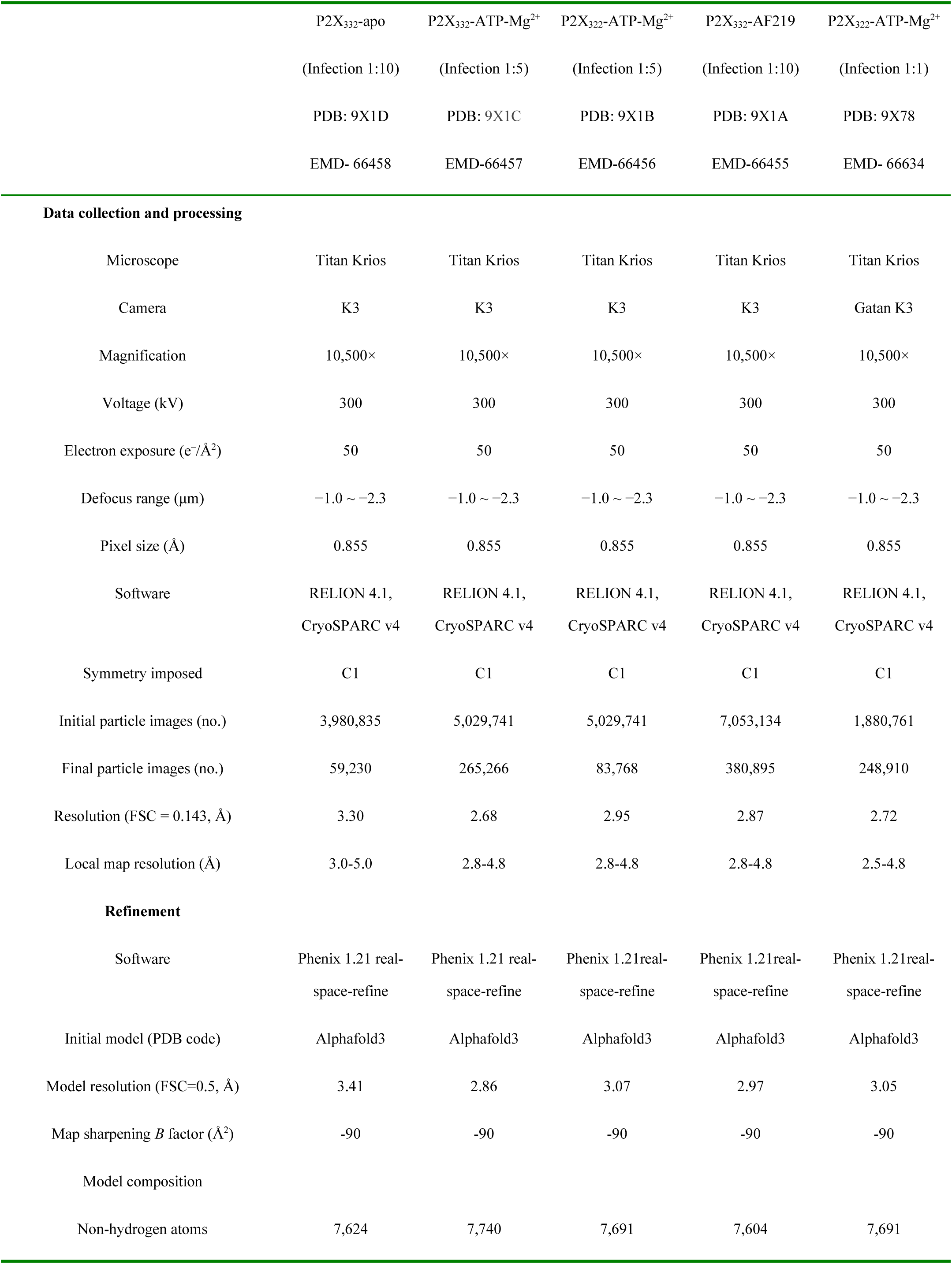
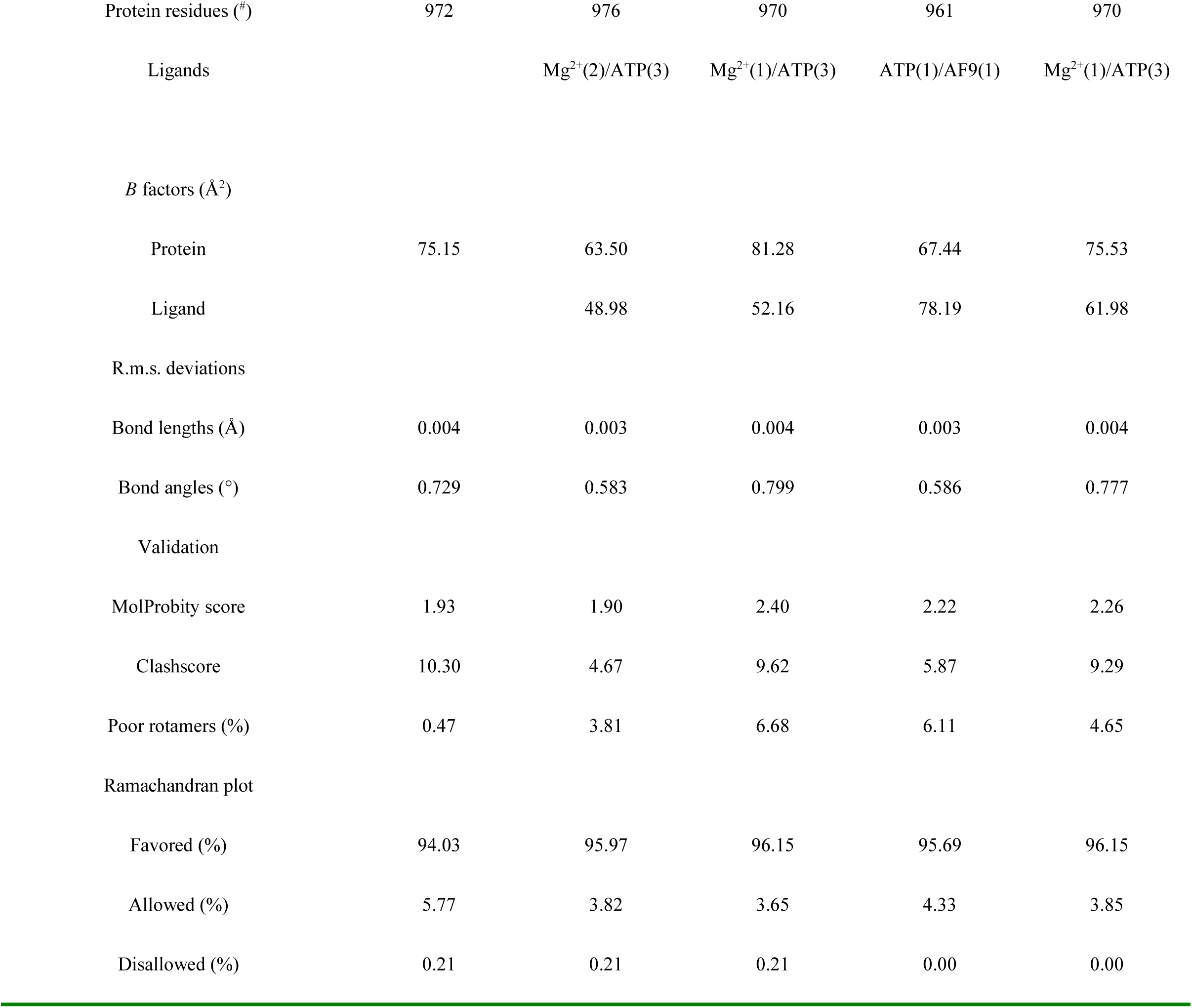
Cryo-EM data collection, refinement and validation statistics.

